# Fine scale mapping of genomic introgressions within the *Drosophila yakuba* clade

**DOI:** 10.1101/152421

**Authors:** David A. Turissini, Daniel R. Matute

**Author notes:** Correspondence: Biology Department, University of North Carolina, 250 Bell Tower Road, Chapel Hill, 27599.

## Abstract

The process of speciation involves populations diverging over time until they are genetically and reproductively isolated. Hybridization between nascent species was long thought to directly oppose speciation. However, the amount of interspecific genetic exchange (introgression) mediated by hybridization remains largely unknown, although recent progress in genome sequencing has made measuring introgression more tractable. A natural place to look for individuals with admixed ancestry (indicative of introgression) is in regions where species co-occur. In west Africa, *D. santomea* and *D. yakuba* hybridize on the island of São Tomé, while *D. yakuba* and *D. teissieri* hybridize on the nearby island of Bioko. In this report, we quantify the genomic extent of introgression between the three species of the *Drosophila yakuba* clade (*D*. *yakuba, D. santomea*), *D. teissieri*). We sequenced the genomes of 86 individuals from all three species. We also developed and applied a new statistical framework, using a hidden Markov approach, to identify introgression. We found that introgression has occurred between both species pairs but most introgressed segments are small (on the order of a few kilobases). After ruling out the retention of ancestral polymorphism as an explanation for these similar regions, we find that the sizes of introgressed haplotypes indicate that genetic exchange is not recent (>1,000 generations ago). We additionally show that in both cases, introgression was rarer on *X* chromosomes than on autosomes which is consistent with sex chromosomes playing a large role in reproductive isolation. Even though the two species pairs have stable contemporary hybrid zones, providing the opportunity for ongoing gene flow, our results indicate that genetic exchange between these species is currently rare.

**AUTHOR SUMMARY:** Even though hybridization is thought to be pervasive among animal species, the frequency of introgression, the transfer of genetic material between species, remains largely unknown. In this report we quantify the magnitude and genomic distribution of introgression among three species of *Drosophila* that encompass the two known stable hybrid zones in this genetic model genus. We obtained whole genome sequences for individuals of the three species across their geographic range (including their hybrid zones) and developed a hidden Markov model-based method to identify patterns of genomic introgression between species. We found that nuclear introgression is rare between both species pairs, suggesting hybrids in nature rarely successfully backcross with parental species. Nevertheless, some *D. santomea* alleles introgressed into *D. yakuba* have spread from São Tomé to other islands in the Gulf of Guinea where *D. santomea* is not found. Our results indicate that in spite of contemporary hybridization between species that produces fertile hybrids, the rates of gene exchange between species are low.

## INTRODUCTION

When two species hybridize, produce fertile hybrids and persist, three outcomes are possible. First, genes from one of the species might be selected against in their hybrids thus removing “foreign” genes from the gene pool, with the rate of removal being proportional to the product of the population size and the strength of selection [1-4]. Second, some alleles will have no fitness effects and may be retained in the population or lost due to drift. Finally, some introduced genes could be maintained in the population because they are advantageous ([2], [3]; but see [4] for additional possibilities). Such introgressed alleles can be a source of novel genetic (and phenotypic) variation.

The frequency and fate of introgressed alleles has been investigated in only a few cases (e.g., [5-11] among many others; reviewed in [12,13]), and the susceptibility of genomes to introgression is the target of lively debate among evolutionary biologists. Obtaining conclusive evidence about the magnitude of introgression is difficult and has led to two general views in speciation research. Some maintain that genomes are co-adapted units that can tolerate very little foreign contamination [14-16]. Others argue that closely related species differ only in a few distinct genomic regions responsible for reproductive isolation and can not only tolerate considerable introgression elsewhere [2,17] but may even benefit from it [18-20]. In reality, both instances occur, but to understand how prevalent introgression is during the speciation process, we require systematic assessments of the rate and identity of introgressions in varied biological systems

Current efforts to detect introgression have found the process to be pervasive in nature (e.g., [13,21-23]). Yet, one of the main limitations of this inference is that most models of introgression are tailored to detect recent introgression where introgressed haplotypes are found in large, contiguous blocks [24]. Powerful analytic tools such as HAPMIX [25], ELAI [26], ChromoPainter [27] and others heavily rely on linkage disequilibrium or phased genomic data which makes them inapplicable to many organisms [28-30]. A second limitation is that some methods [31-33] will estimate the amount of introgression but not specific introgressed genomic regions, precluding the measuring the frequency of introduced segments. Ideally methods for detecting introgressions would be able to identify introgressed segments within individuals and not need haplotype information and/or phased genotypes which might not be available for all taxa.

Even though *Drosophila* has been a premier system for studying how reproductive isolation evolves, until recently interspecific gene flow within the taxon has been understudied because hybrid zones were either unknown or uncharacterized. Yet, neither gene flow, nor hybrid zones are absent in the *Drosophila* genus [8,34-36].

The *D. yakuba* species clade is composed of three species (*D. yakuba*, *D. santomea,* and *D. teissieri*) whose last common ancestor is thought to have existed ~1.0 million years ago (MYA) [37]. *Drosophila yakuba* is a human-commensal that is widespread throughout sub-Saharan Africa and is also found on islands in the Gulf of Guinea [38-40]. *Drosophila teissieri,* like *D. yakuba,* is also distributed across large portions of the continent but is largely restricted to forests with *Parinari* (Chrysobalanaceae) trees [41-43]. *Drosophila santomea* is restricted to the island of Sao Tomé in the Gulf of Guinea. *Drosophila yakuba* also lives on São Tomé and occurs at low elevations (below 1,450 m) and is mostly found in open and semidry habitats commonly associated with agriculture and human settlements [39,44]. In contrast, *D. santomea* is endemic to the highlands of São Tomé where it is thought to exclusively breed on endemic figs (*Ficus chlamydocarpa fernandesiana,* Moraceae; [45]). *Drosophila yakuba and D. santomea* produce sterile male and fertile female hybrids, and the two species co-occur in a hybrid zone in the midlands on the mountain Pico de São Tomé [38,44,46]. Backcrossed females and some males are fertile [47,48]. Oddly, a second stable hybrid zone composed exclusively of hybrid males occurs on top of Pico de São Tomé largely outside the range of the two parental species [49].

Within *Drosophila,* the *D. santomea/ D. yakuba* hybrid zone is the best studied for at least three reasons. First, it has the highest known frequency of hybridization: on average, 3-5% of *yakuba* clade individuals collected in the midlands of São Tomé are F1 hybrids [44]. Second, the hybrid zone is stable and has persisted since its discovery in 1999 [38,44,49,50] which makes it one of the two stable hybrid zones in the genus (along with *D. yakuba/D. teissieri,* see below). Third, F1 hybrids are easily identified by their characteristic abdominal pigmentation [51,52]. Advanced intercrosses are harder to identify since pigmentation patterns regress toward the parental species in just one or two generations of backcrossing [52].

*Drosophila teissieri is* the sister species to the *D. yakuba/ D. santomea* (*yak/sah*) dyad. It is distributed throughout tropical Africa and is thought to have occupied a much larger range before humans expanded into the forests of Sub-Saharan Africa [43,53]. Even though it breeds at higher elevations (over 500m), it is commonly found in the same locations where *D. yakuba* is collected [41,43,54]. The species is thought to be a narrow specialist of the ripe fruit of *Parinari* [41,53]. The nuclear genomes of *D. yakuba and D. teissieri* differ by numerous fixed inversions, which were long thought to preclude hybridization ([55]but see [37,56]). Nonetheless, *D. teissieri* does produce hybrids with *D. yakuba and D. santomea* in the laboratory [57]. F1 females (from both reciprocal directions of the cross) and some backcrossed individuals are fertile [57]. Field collections have also found a stable and narrow hybrid zone between *D. yakuba* and *D. teissieri in* the highlands on the island of Bioko at the interface between cultivated areas and secondary forest [54].

Across both the *yak/san* and *D. yakuba/D. teissieri*(*yak/tei*) hybrid zones, little is known about the genomic and geographic distributions of introgression. Genealogies from two mitochondrial genes (*COII* and *ND5;* 1,777 bp) show *D. yakuba* and *D. santomea* individuals interspersed, especially for individuals collected in the hybrid zone of São Tomé. A mitochondrial genome survey still shows admixture but to a lesser extent [58]. Additionally, mitochondrial divergence between the three species is much lower than expected given the levels of divergence observed for the nuclear loci [37,56,58]. The discrepancy has been interpreted as mitochondrial introgression resulting in the homogenization of the mitochondrial genome within the clade.

Despite this emphasis on mitochondrial introgression, little is known about the extent of nuclear introgression between *D. yakuba* and *D. santomea.* Preliminary genetic analyses [37,39] found evidence of gene flow for two autosomal loci that showed low levels of divergence relative to the other typed loci. Beck et al. [59] also found nuclear introgression from *D. yakuba* into *D. santomea* of genes coding for nuclear pore proteins that interact with mitochondrial gene products. No study has however addressed the possibility of gene flow between *D. yakuba* and *D. teissieri,* and no systematic genomic effort has addressed the magnitude of gene flow between *D. yakuba and D. santomea.* We focus on measuring whether, similar to the mitochondrial genome, the nuclear genomes within the *yakuba* species group show evidence of introgression.

To characterize introgression within the *yakuba* species group we developed a new statistical framework to identify introgressed regions of the genome. Since linkage disequilibrium in *Drosophila* usually decays fast (on the order of a few hundred base pairs; [60] but see [61,62]), we were not able to use available LD-based methods to detect introgression. Our method (Int-HMM) relies on the identification of stretches of differentiated SNPs, and uses a hidden Markov Model (HMM) approach to identify introgressed regions from unphased whole genome sequencing data. The framework does not require pre-identified pure-species samples from allopatric regions, and is able to identify introgressions on the order of 1kb with low false positive rates (<1%). We used this model to quantify the magnitude of introgression between *D. yakuba/ D. santomea* and *D. yakuba/ D. teissieri.* We found that nuclear introgression is rare between the two species pairs despite hybrids being identified in nature. We also found that some alleles that have introgressed from *D. santomea* into *D. yakuba* have spread from São Tomé to other islands in the Gulf of Guinea where *D. santomea* is not currently found.

## RESULTS

### Molecular divergence and approximate species divergence times

*Drosophila yakuba* and *D. santomea* had been previously estimated to have diverged ~393,000 years ago [51] and ~500,000 years ago [37]. Bachtrog et al. [37] also estimated the divergence time between *D. yakuba* and *D. teissieri* to be ~1 million years ago. However, these estimates were based on only a few nuclear loci (N ~ 15 DNA fragments). We estimated the divergence times using the number of synonymous substitutions (K_s_) from 14,267 genes [57] using the same approach as Llopart et al. [63]. We had previously estimated K_s_ between *D. yakuba* and *D. santomea* as 0.0479 and between *D. yakuba* and *D. teissieri* as 0.1116 [57]. We then compared these K_s_ values to the K_s_ of 0.1219 between *D. melanogaster*and *D. simulans* [64], which are estimated to have diverged 3 million years ago [65]). Assuming comparable substitutions rates between the two groups, we obtained estimated divergence times of 1.18 million years ago for the *D. yakuba - D. santomea* split and 2.75 million years ago for the divergence time between *D. yakuba* and *D. teissieri.* The level of divergence between the latter pair is surprising considering that *D. yakuba* and *D. teissieri* produce fertile F1 females and have a stable hybrid zone on the island of Bioko, while species with similar divergence (e.g., *D. melanogaster and D. simulans* whose divergence time is estimated to be between 3 an 5MYA; [66,67]) produce sterile or inviable hybrids ([68]; Table S10 in [57]).

### PCA

We used principle component analyses (PCA) to investigate genomic divergence among all three *D. yakuba*-clade species. Analyses were completed separately for the *X* chromosome and the autosomes (Figure S1) as sex chromosomes and autosomes often experience different demography and selection patterns [69,70]. Principal components (PCs) 1, 2, and 3 separate the species for both the *X* and autosomes. Collectively the first three PCs explain 77.8% of the variation for the autosomes and 79.9% of the variation for the *X* chromosome. Among the three species, *D. yakuba* exhibited the most variation for all 3 principle components. We did not observe any overlap between the species. Four *D. santomea* lines were slightly more similar to *D. yakuba* for both PC 1 and 2 on the autosomes and were not included in the donor population when selecting markers for the *san-* into-*yak* HMM analysis (see below).

### Detecting Evidence of Introgression

#### (i) Patterson’s D statistic

We first explored the occurrence of introgression using multiple versions of the Patterson’s D statistic (i.e., ABBA BABA test, [31,33,71]). Because the test requires potentially admixed populations and a population without gene flow of the recipient species, we tested for gene flow from *D. santomea* into *D. yakuba.* We used *D. yakuba* from the continent as the outgroup and *D. yakuba* from São Tomé as the potential recipient. (*Drosophila santomea* has a relatively small range, and we have been unable to find bona fide allopatric populations.) We found significant introgression between *D. yakuba* and *D. santomea* but the average direction of introgression depends on the choice of outgroup (Table 1). If *D. teissieri* is the outgroup of the test, the most common direction of introgressions *is D. santomea* into *D. yakuba* (*san-into-yak*). If *D. melanogaster* is the outgroup of the test, the most common direction of introgressions *is D. yakuba* into *D. santomea* (*yak-*into-*san*). All of our other analyses (see below) however indicate that introgression is more common in the *yak-into-san* direction indicating that *D. melanogaster reads* might not map well to the *D. yakuba* genome due to the increased divergence between *D. melanogaster* and *D. yakuba* (Ks~0.26 [72]; especially at regions less constrained by selection such as intergenic regions); the choice of outgroup is clearly relevant.

**TABLE 1.** D-statistic variations (D [31,32] and f_d_ [71]) show evidence for admixture between *D. yakuba* (mainland, São Tomé hybrid zone—HZ—, and other islands—Bioko and Principe—) and *D. santomea* and between *D. yakuba and D. teissieri.* Note, that the negative numbers of D indicate that the average direction of the introgression goes from the population assigned as putatively recipient to the population assigned as putatively donor.

| Allopatric population | Recipient population | Donor population | outgroup | D | Z-score | $f_d$ |
| --- | --- | --- | --- | --- | --- | --- |
| *D. yakuba* mainland | *D. yakuba* HZ | *D. santomea* | *D. teissieri* | 0.015173 | 189.739 | 0.001862 |
| *D. yakuba* mainland | *D. yakuba* other islands | *D. santomea* | *D. teissieri* | 0.030080 | 311.41 | 0.003607 |
| *D. yakuba* mainland | *D. yakuba* HZ | *D. santomea* | *D. melanogaster* | -0.019728 | -292.231 | -0.003141 |
| *D. yakuba* mainland | *D. yakuba* other islands | *D. santomea* | *D. melanogaster* | -0.023744 | -286.188 | -0.003742 |
| *D. teissieri* (not Bioko) | *D. teissieri* (Bioko) | *D. yakuba* | *D. melanogaster* | -0.030094 | -754.157 | -0.002709 |

Next, we computed Patterson’s D statistic looking for gene flow between *D. yakuba* and *D. teissieri.* We focused on *D. teissieri* from the hybrid zone on Bioko as the recipient population of introgression. We find evidence for introgression in this species pair (Table 1). The average direction of gene flow is from *tei-*into*-yak (D. yakuba* from Bioko). These results provide evidence that there has indeed been genetic exchange between the two species pairs that naturally hybridize in the *yakuba* species complex.

#### (ii) Treemix

We used the program *Treemix* to identify gene flow between species and populations within the *D. yakuba* clade. We ran *Treemix* separately for the *X* chromosome (Figures 1A, Figure S2) and the autosomes (Figures 1B, Figure S3). For the *X* chromosome, *Treemix* found 2 admixture events within *D. yakuba* and none between species. The first event goes from the lowlands of São Tomé to the African mainland (weight = 0.21) and the second from the lowlands of São Tomé to the islands of Príncipe and Bioko (weight=0.046). For the autosomes, *Treemix* found evidence of 4 migration events, one of them between populations of *D. yakuba* (weight = 0.375), the other three events between species. One of these events indicates gene flow from *D. yakuba* on the islands of Príncipe and Bioko to *D. santomea* (weight=0.104), the second from *D. santomea* to mainland *D. yakuba* (weight = 0.08), and the third from *D. teissieri* to *D. yakuba* at the hybrid zone (with *D. santomea*) on São Tomé (weight = 0.005). These results suggest that there has been introgression between the species in the *yakuba* complex. They also suggest that introgression is more likely to occur on the autosomes than on the *X* chromosome. Next, we explored the fine-scale patterns of introgression in the nuclear genomes of the three species.

**Figure 1.**
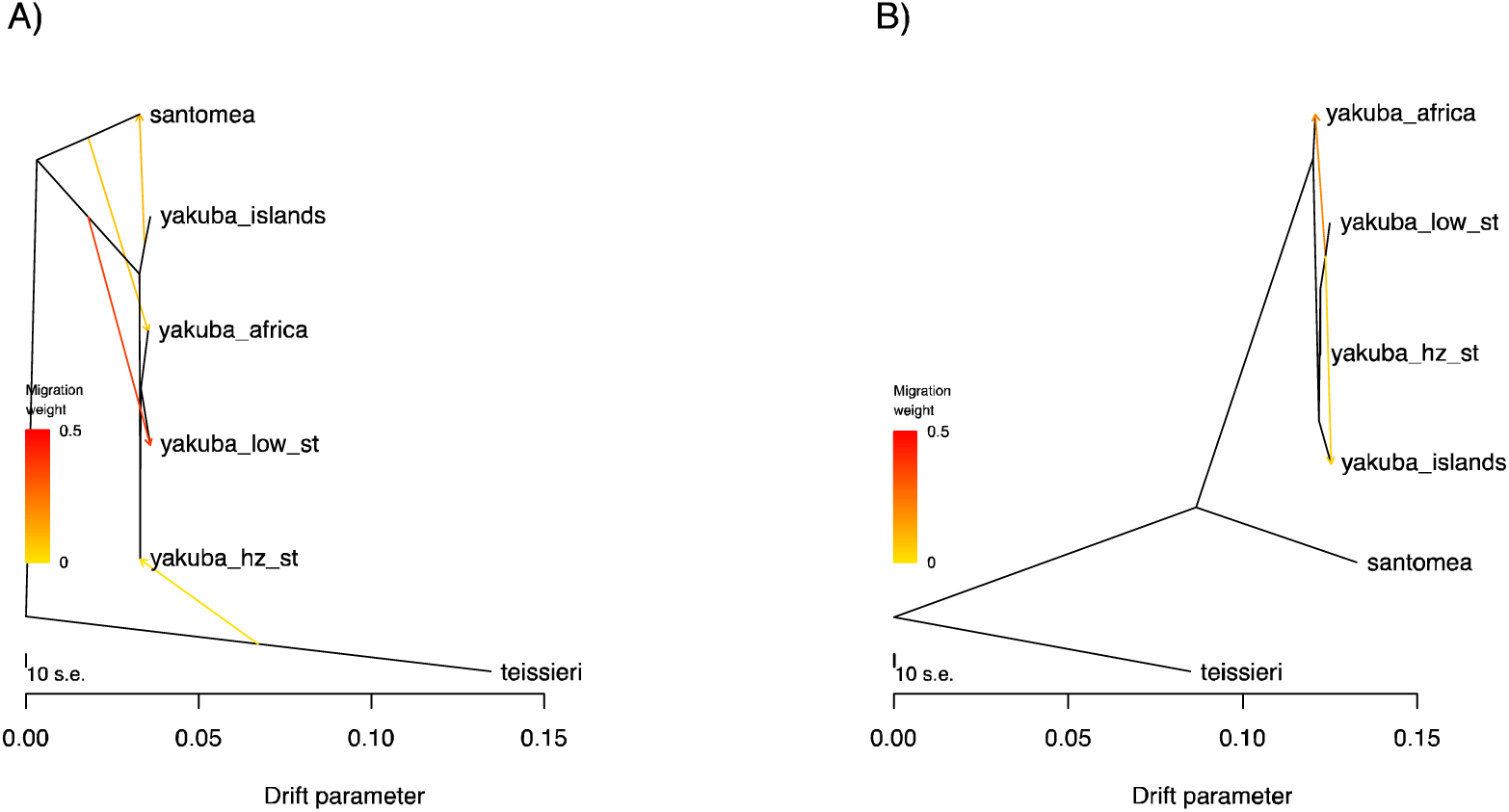
*Treemix results* for the *D. yakuba* clade indicate gene flow has occurred among species of the *yakuba* clade. *Treemix* trees with the best supported number of migration edges. *D. yakuba* has been split into four populations: “africa” (Cameroon, Kenya, Ivory Coast), “islands” (Principe and Bioko), “low_st” (lowlands of São Tomé), and “hz_st” (hybrid zone on São Tomé). **A)** Autosomal tree with 4 migration edges. **B)** *X* chromosome tree with 2 migration edges. Other demographic scenarios are shown in Figures S2 and S3.

### Linkage Disequilibrium

Linkage disequilibrium decays rapidly in *D. melanogaster.* r^2^ decays to 0.2 within 5,000bp ([35,60], but see [61,62]). Such rapid decay of LD seriously constrains the possibility of using long-range LD to detect admixture since most methods that use LD rely on identifying within population haplotype variation. The low levels of LD seen in *Drosophila* preclude identifying haplotypes thus preventing such methods from working properly. We evaluated whether similar patterns of LD decay exist in the three species of the *yakuba* species clade. We measured linkage disequilibrium (LD) for all three species in the *D. yakuba* clade using PLINK [73]. For both the *X* chromosome and the autosomes, LD declined sharply at a scale of ~300bp before leveling off (Figure S4). At a distance of 1kb, the average r^2^ for *D. yakuba* was 0.0652 for the autosomes and 0.0464 for the *X* chromosome (Figure S4A), for *D. santomea* the average r^2^ was 0.1347 for the autosomes and 0.1518 for the *X* chromosome (Figure S4B), and for *D. teissieri the* average r^2^ was 0.1517 for the autosomes and 0.134 for the *X* chromosome (Figure S4C). This fast decay indicated the need to develop a framework to detect introgressed alleles that does not rely on LD.

### Identifying introgressed tracts

#### (i) Performance of the method: simulation results

We developed a Hidden Markov Model (HMM) to identify specific introgressed regions, Int-HMM. First, we determined the sensitivity of the method by assessing whether it could detect simulated introgressions. We simulated independent introgressions with sizes ranging from 100bp up to 100kb for both directions of gene flow in admixed genomes between *D. yakuba* and *D. santomea* and between *D. yakuba* and *D. teissieri* with some introgressed regions being homozygous and others heterozygous. Then, we used Int-HMM on the simulated data. We found that Int-HMM correctly identified a majority of introgressed regions with the percentage of correctly identified introgressions increasing with the size of the introgressed region (Figure 2). Int-HMM is more reliable at identifying homozygous introgressions than heterozygous ones. For homozygous introgressions, the false negative rates were less than 10% for all introgression sizes greater than or equal to 4kb for introgressions between *D. yakuba* and *D. santomea* and 2kb for introgressions between *D. yakuba and D. teissieri.* For heterozygous introgressions, Int-HMM performed better with introgressions between *D. yakuba* and *D. teissieri where* the false negative rate was less than 10% for introgressions greater than or equal to 3kb. The rate did not drop to 10% for *D. yakuba* and *D. santomea* introgressions until the size was at least 15kb. The model likely performs less well for smaller regions due to a relative paucity of informative markers. False positive rates were negligible in all cases and were always less than 0.3% (Table S2).

**Figure 2.**
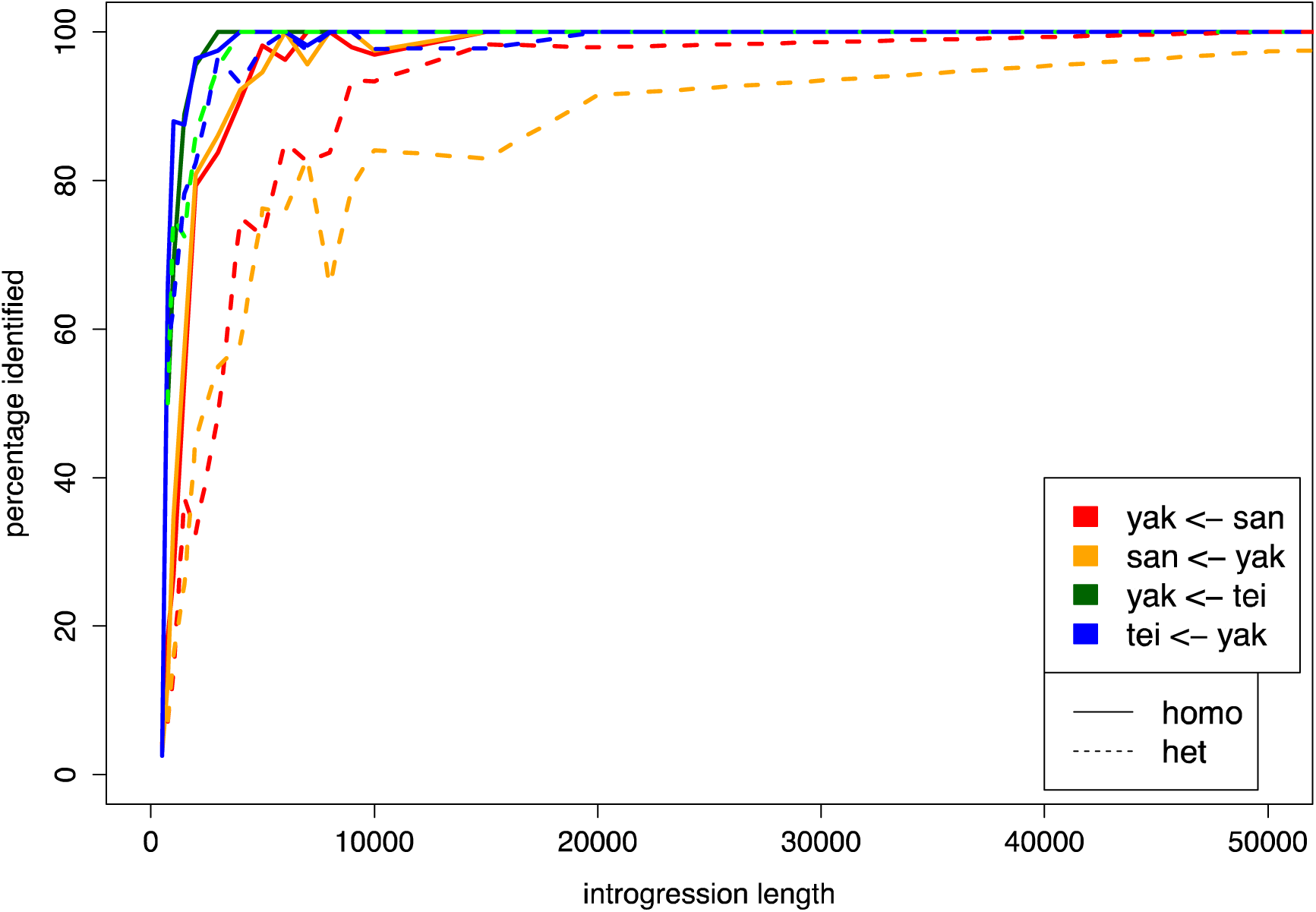
Proportion of correctly identified simulated introgressions by Int-HMM. The HMM successfully identified over 80% of introgressions longer than 10kb for all directions of introgression. It consistently performed better at identifying homozygous introgressions than heterozygous ones. Additionally, it identified higher percentages of introgressions between *D. yakuba* and *D. teissieri* than those between *D. yakuba* and *D. santomea.*

#### (ii) HMM results

We identified the specific genomic regions that had introgressed from among species in the *yakuba* species complex. We looked for introgressed regions in both directions between *D. yakuba* and *D. santomea* (*san*-into*-yak*, *yak-*into *-san*) and between *D. yakuba* and *D. teissieri* (*tei-into-yak, yak-into-tei*) using the newly developed Int-HMM. The HMM was run individually on the genomic data from each genotype call (SNP) which had between 933,776 and 951,384 markers for *san*-into-*yak,* between 907,959 and 923,227 for *yak-*into*-san,* between 1,867,399 and 1,888,413 markers for *tei-*into*-yak,* and between 2,275,453 and 2,468,955 markers for *yak-*into*-tei* (Table S1). On average the markers were separated by 127-133bp for *san-*into*-yak,* 131-133bp for *yak-*into*-san,* 64-65bp for *tei-*into*-yak,* and 49-53bp for *yak-*into*-tei.* The HMM returned a probability that each marker was either homozygous for the recipient species, heterozygous, or homozygous for the donor species; adjacent sites with identical, most-probable states were combined into tracts. We next describe the results for each species pair.

##### Introgression tracts: *D. yakuba/D. santomea*

*Drosophila yakuba* and *D. santomea* hybridize in the midlands of São Tomé and form a stable hybrid zone with the highest rate of hybridization known in *Drosophila.* We hypothesized that we would find a rate of introgression comparable with the rate of hybridization. Yet, and despite the continuous and ongoing hybridization between these two species, we found evidence that introgression at the genomic level is rare. Of the 17 *D. santomea* lines we assessed, on average 0.35% of the *D. santomea* genome was introgressed from *D. yakuba* with individual levels ranging from 0.1% (Qiuja630.39) up to 1.04% (san_Field3) (Figure 3A, Table S3). The introgressions in the different lines covered different genomic regions, and cumulatively, they spanned 3.48% of the genome. We found comparable levels of introgression from *D. santomea* into *D. yakuba* with an average of 0.22% of the *D. yakuba* genome originating from *D. santomea.* Together, the introgressions across the 56 lines covered 5.56% of the genome. The magnitude of the introgressed genetic material varied almost two orders of magnitude across lines: individual levels ranged from 0.012% (3_16) up to 1.20% (Anton_2_Principe) (Figure 3A, Table S3).

**Figure 3.**
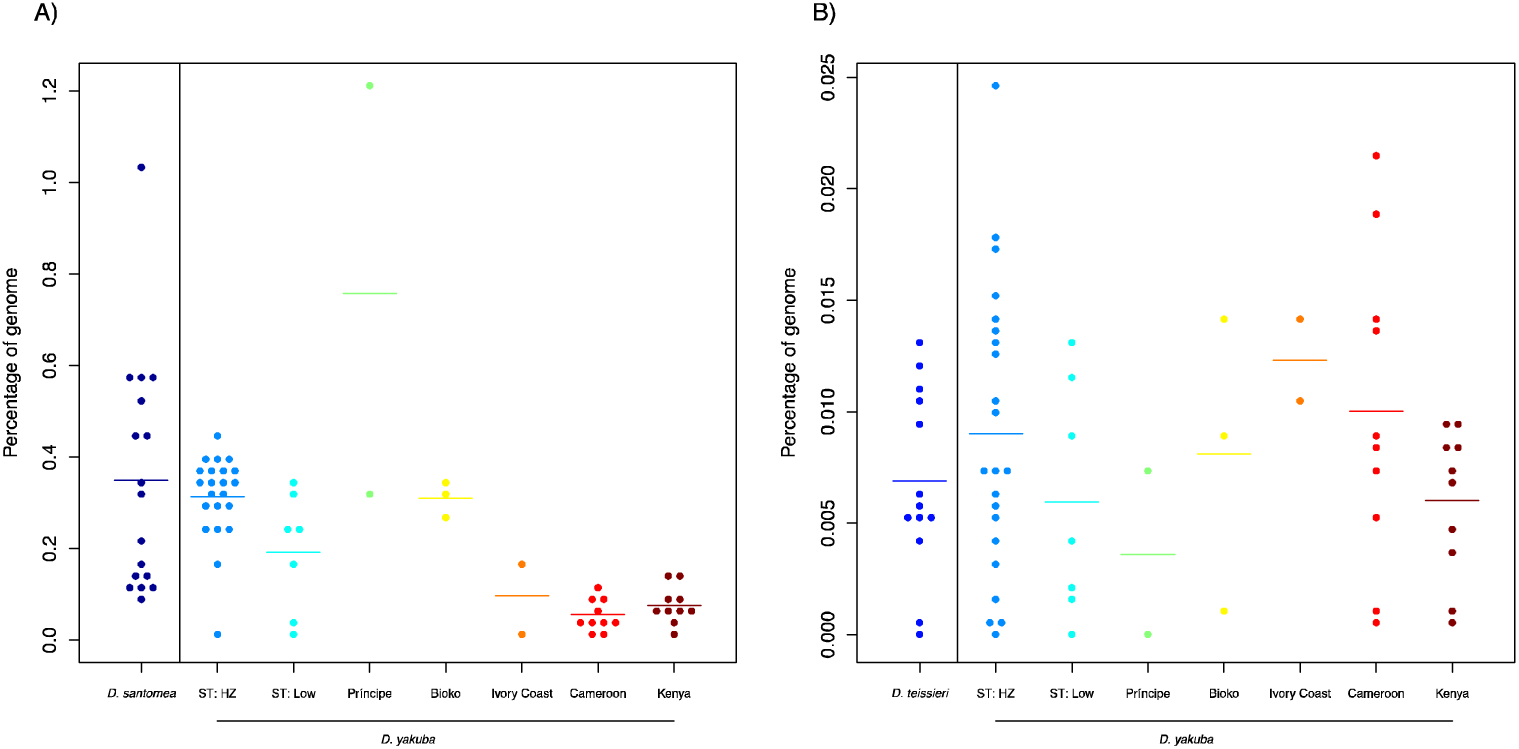
Percentage of genome introgressed between each species pair. Percentage of the genome that was introgressed for each line as determined by the cumulative length of introgression tracts identified by Int-HMM. *D. yakuba* has been divided into geographical populations where ‘ST: HZ’ refers to the São Tomé hybrid zone and ‘ST: Low’ to the lowlands of São Tomé. **A)** *yak-*into*-san* and *san-*into*-yak* introgressions. **B)** *yak-*into*-tei* and *tei*-into-*yak* introgressions.

A majority of the introgressed regions were intronic (*san-*into*-yak* 55.7%, *yak-*into*-san.* 49.3%) and intergenic (*san-*into*-yak.* 35.2%, *yak-*into*-san.* 43.1%) (Table S4). In the *san-*into*-yak* direction, the RNA coding regions (CDS, 3’ prime UTR, and 5’ prime UTR) are observed more than expected by chance. In the *yak-*into-*san*, 10kb inter and 3’ prime UTR are more likely than random to be included in introgressions. Each type of sequence had similar marker densities; thus, it is unlikely that differences in read mapping affected these results (Table S4).

Next, we compared the magnitude of introgression for the two reciprocal directions of each cross. Globally, there was significantly more introgression from *D. yakuba* into *D. santomea* (Mann-Whitney U = 301, p= 0.0228), than from *D. santomea* into *D. yakuba.* Since *D. yakuba* has a geographic range that dwarfs that of *D. santomea,* we also repeated the species comparison excluding *D. yakuba* flies from Cameroon and Kenya, collection sites completely outside of *D. santomea’*s range. When only *D. yakuba* lines from near the Gulf of Guinea were included, levels of introgression were the same in both directions (Mann-Whitney U = 288, p= 0.7413).

We next asked whether the magnitude of the *san*-into-*yak* introgression varied across *D. yakuba* lines from different locations. We found more introgression in *D. yakuba* flies collected within the hybrid zone with *D. santomea* (midlands of São Tomé; genomic average across individuals = 0.314%) than in flies collected on the island but at lower elevations outside of the hybrid zone (genomic average across individuals = 0.192%) (Mann-Whitney U = 123, p=0.018). Surprisingly, *D. yakuba* flies from the hybrid zone did not have more introgression than flies from the nearby islands of Bioko (Mann-Whitney U = 41, p=0.550) or Príncipe (Mann-Whitney U = 12, p=0.355).

We next analyzed the distribution of sizes of the haplotypes shared across the *yak/san* species boundary. The average tract size for the *D. yakuba* into *D. santomea* introgressions was 6.8kb with a maximum size of 112kb (Figure 4A). The average tract size for *D. santomea* into *D. yakuba* introgressions was 6kb with a maximum size of 959.5kb (Figure 4B). Interestingly, the 959.5KB tract was from a line from the island of Príncipe where *D. santomea* is not known to currently exist (Figure S5). The next largest tract was 120.5kb and was seen in two lines from the hybrid zone on São Tomé (Cascade_SN6_1, Montecafe_17_17).

**Figure 4.**
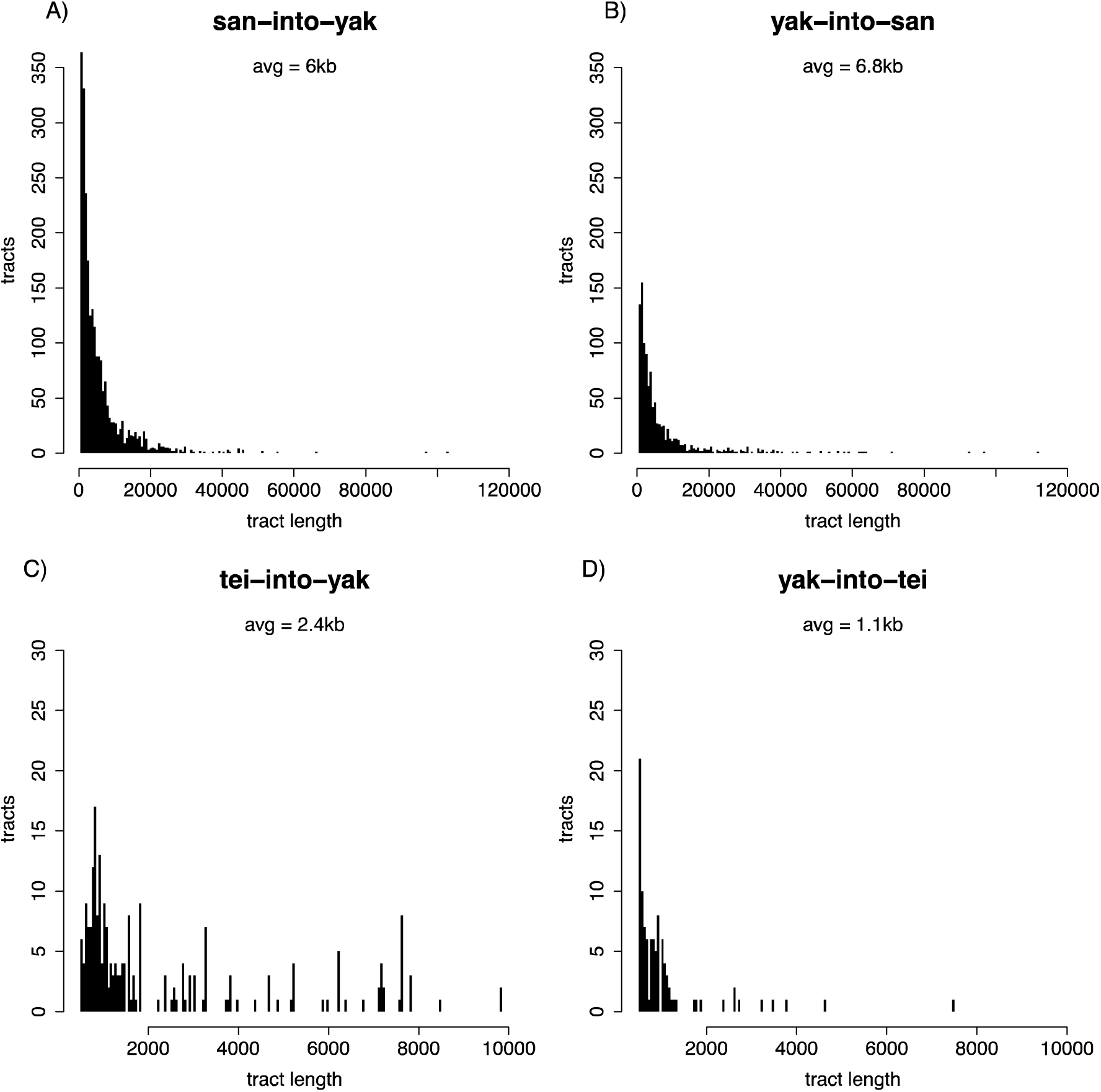
Introgression tracts are generally small. Distributions of tract sizes. Note that tracts smaller than 500bp were not included in the analysis. **A)** *san-into-yak.* The distribution has been truncated to exclude a single large 959kb tract shown in Figure S5. **B)** *san-*into*-yak.* **C)** *tei-*into*-yak.* **D)** *yak-*into*-tei.*

##### Introgression tracts: *D. yakuba/D. teissieri*

*Drosophila yakuba* and *D. teissieri* also hybridize in the highlands of Bioko in a very narrow and geographically restricted hybrid zone [54]. As expected, given their divergence and the narrow hybrid zone, introgression from *D. yakuba* into *D. teissieri* (*yak-*into-*tei*) was rare. Among the 13 *D. teissieri* lines, on average 0.0074% of the genome originated from *D. yakuba* with individual values ranging from 0% (Anton_2_Principe, Montecafe_17_17, SJ_1) to 0.0129% (Selinda) (Figure 3B, Table S3). Together the introgressions span 0.0669% of the genome. There was no difference in the amount of *yak-*into*-tei* introgression between the *D. teissieri* population on Bioko where a known hybrid zone is located and flies from outside Bioko (Mann-Whitney U=22, P = 0.826).

For the 56 *D. yakuba* lines, on average 0.0086% of the genome of the two species has crossed the species boundary. Individual percentages ranged from 0% (cascade_2_1) to 0.0244% (2_8) (Figure 3B, Table S3). Collectively, the introgressions span 0.0914% of the genome. In the *tei*-into-*yak* direction the only type of region that shows an enrichment is ‘introns’, while in the reciprocal direction, *yak-into-tei,* both intergenic and intronic regions show an enrichment. As with the *yak/san* case, all type of sequences had similar markers densities (Table S4).

We also compared the magnitude of introgression in both directions. Similar to the hybrid zone between *yakuba and santomea,* we found no asymmetry in the amount of introgression between *D. yakuba* and *D. teissieri* (Mann-Whitney U = 354.5, p= 0.5426). The *D. yakuba* lines with the highest levels of introgression from *D. teissieri* were from Cameroon and the *yak/san* hybrid zone on São Tomé (Figure 3). Whereas *D. teissieri is* also present in Cameroon, this species (or its plant host *Parinan*) has never been collected on the island of São Tomé.

Finally, we assessed the distribution of sizes of the haplotypes shared across the *yak/tei* species boundary. The average tract size for *D. yakuba* into *D. teissieri* introgressions was 1.1kb with a maximum size of 7.5kb (Figure 4C). *Drosophila teissieri* into *D. yakuba* introgressions were larger than those in the reciprocal cross with an average of 2.4kb and a maximum size of 9.8kb (Figure 4D). The amount of exchanged genetic material was larger in the latter direction of the cross (Mann-Whitney U = 17,034, P = 5.37 × 10^-13^).

##### Species pair comparisons

Since the split between *D. yakuba* and *D. santomea* occurred much more recently than that between *D. teissieri* and *D. yakuba*, fewer genetic incompatibilities will have evolved. Selection will purge alleles linked with those negatively selected alleles. Thus, we expected to find more introgression between *D. yakuba* and *D. santomea* than between *D. yakuba* and *D. teissieri.* Indeed, there was significantly more *san-* into *-yak* than *tei*-into-*yak* introgression (Mann-Whitney U = 44, P< 1 × 10^-15^) and *yak-*into*-san* than *yak-*into*-tei* introgression (Mann-Whitney U = 0, P = 6.96 × 10^-6^).

There are several similarities in the patterns of introgression in the two species pairs. First, in all four cross directions, introgressions were present at low frequencies (Figure 5). The average frequencies were 11.1% (*san-*into*-yak*), 11.8% (*yak-*into*-san*), 10.8% (*tei-*into*-yak),* and 15.1% (*yak-*into*-tei*).

**Figure 5.**
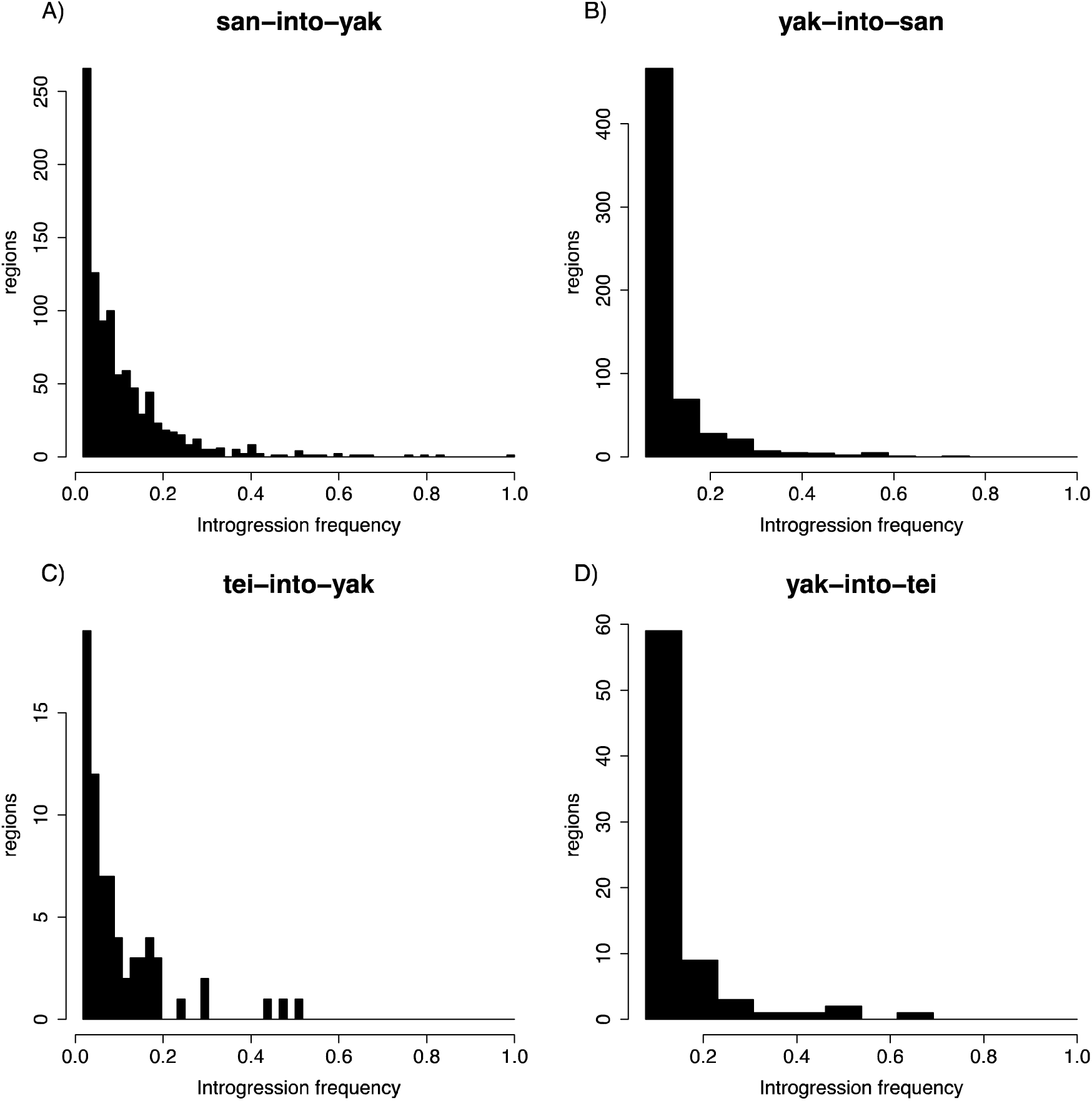
Most introgressions are present at low frequencies. Frequencies of introgressed regions defined as inclusive sets of overlapping individual introgressions. **A)** *san-*into*-yak.* **B)** *yak-*into*-san.* **C)** *tei-*into*-yak.* **D)** *yak-*into*-tei.*

Second, for both species pairs, introgression tracts were not uniformly distributed across the genome (Figure 6). We observed less introgression on the *X* chromosome than on autosomes in all four directions (permutation tests; *san*-into-*yak.* P < 0.0001, *yak-into-san.* P < 0.0001, *tei-*into*-yak* P < 0.0001, *yak-*into*-ter.* P = 0.0198; Figure S6). Intriguingly, a region at the start of the *X* chromosome where we did not find any *san*-into-*yak* introgression and limited *yak-*into*-san* introgression corresponds with a QTL implicated in hybrid male sterility between the two species [48]. Finally, we not only found less *X*-linked introgression, but introgressed tracts were also shorter on the *X* chromosome than on the autosomes: 3.16kb versus 6.05 kb (*san-*into*-yak*), 3.34kb versus 6.9kb (*yak-*into*-san*), and 0.87kb versus 1.08kb (*yak-*into*-tei*). Notably, we did not find any *X*-linked introgressions for *tei-*into*-yak.*

**Figure 6.**
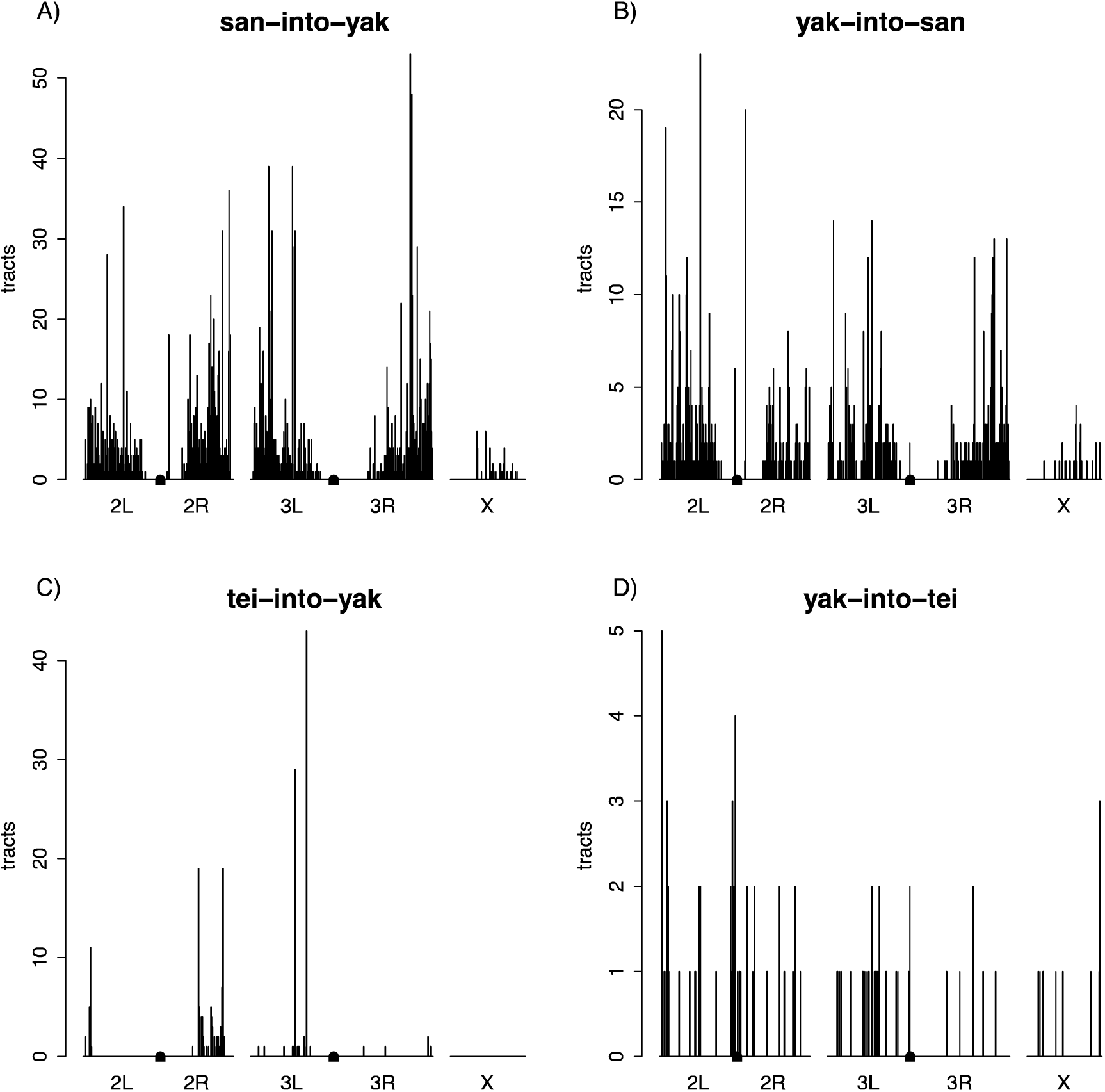
Genomic distributions of introgression tracts. Centromeres are denoted by rectangles in the center of chromosomes 2 and 3. **A)** *san-*into*-yak.* **B)** *yak-*into*-san.* **C)** *tei-*into*-yak* **D)** *yak-*into*-tei.*

### Dating introgression

The percentage of the genome containing introgressions and the size distribution of introgression tracts within a population contain information on the timing and rates of historic introgression. The size of introgressions we observed are surprisingly small given the stable nature of the two hybrid zones, and the observation of hybrid individuals in nature. These pattern suggest that, despite low levels of hybridization, introgression is old because recombination has broken down introgressed regions over time. To obtain a rough estimate of the age of introgression, we used the program SELAM [74]. Modeling all of the potential demographic and introgression histories would be beyond the scope of this paper. Instead we modeled the simplest hybridization with introgression scenario: a single generation pulse of introgression (i.e., hybrids are formed only once). We recorded the size of the resulting introgression tracts from 50 individuals for 10,000 generations under four different models (i.e., magnitude of the hybridization event). We ran five independent simulations each for initial migration rates m = 0.0001, 0.001, 0.01, and 0.1. We found that the percentage of the genome containing introgressed tracts declined to levels observed between *D. yakuba* and *D. santomea* within 100-300 generations for m = 0.0001, 100-200 generations for m = 0.001, 200 generations for m = 0.01, and 200 generations for m = 0.1 (Figure S7). Simulated percentages fell to levels seen between *D. yakuba* and *D. teissieri* within 1,600 to 8,100 generations for m = 0.0001, 5,800-6,900 generations for m = 0.001, 6,800-7,800 generations for m = 0.01, and 7,800-8,000 generations for m = 0.1 (Figure S7). The average length of introgressed tracts shrunk to levels seen between *D. yakuba* and *D. santomea* within 1,700-6,200 generations with two runs never decreasing as much for m = 0.0001, 7,900-9,800 generations for m = 0.001, 10,000 generations with four runs never decreasing as much for m = 0.01, and no runs decreasing as much for m = 0.1 (Figure S8). The average length decreased to levels seen between *D. yakuba* and *D. teissieri* within 2,700 generations with four runs never decreasing as much for m = 0.0001 and no runs decreasing as much for m = 0.001, 0.01, or 0.1 (Figure S8). The small average length of observed tracts, therefore, suggests that introgression may be old and the original rate of gene flow (m; assuming a single pulse of introgression) was low.

### Ancestral variation

Shared genetic variation between species could result from introgression but may also represent genetic variation present prior to speciation that is still segregating in both species (i.e., incomplete lineage sorting; [75,76]). To assess how likely this scenario was, we looked at the expected number of generations after speciation before ancestral variation was lost and the expected size distribution of ancestral haplotypes. The number of generations before a neutral allele segregating in the ancestral population is lost is 39,800 generations for N_e_=10^4^ and 3,979,933 generations for N_e_=10^6^ (Figures S10A-B, S11A-B). (This of course does not account for trans-specific balancing selection.) We estimated the divergence time between *D. yakuba* and *D. santomea* to be 1 million years (MY) and between *D. yakuba and D. teissieri* to be 2.6MY. We then estimated the number of generations since *D. yakuba* and *D. santomea* diverged to be 26.1 × 10^6^, 17.4 × 10^6^ and 13.0 × 10^6^ generations respectively for generation lengths of 14, 21, and 28 days and 67.8 × 10^6^, 45.2 × 10^6^, and 33.9 × 10^6^ generations respectively for *D. yakuba* and *D. teissieri.* All of the estimates are much older than the ~4 million generations that a SNP is expected to remain polymorphic (Figures S10A-B, S11A-B). It is, therefore, unlikely that the regions of shared ancestry represent ancestral polymorphism and are much more likely to represent introgressed regions. Furthermore, the tracts we identify need to contain at least 10 putatively introgressed SNPs, and the probability of independently observing so many in a row is small. These results strongly argue against ancestral polymorphism occurring in any of the two species pairs.

However, an introgressed fragment could be an ancestral haplotype block that is still segregating in only one of the species. If this is the case, recombination will break down ancestral haplotypes over time. We next looked at the expected distribution of fragment lengths that would still be segregating and are derived from the ancestral species. Between *D. yakuba* and *D. santomea* the 99^th^ quantiles for expected fragment lengths assuming a divergence time of 1 MY and generation lengths of 14, 21, and 28 days were 12bp, 19bp, and 24bp respectively (Figure S9C-E). Assuming a divergence time of 2.6 MY (as estimated above), the respective 99^th^ quantiles for *D. yakuba* and *D. teissieri* were 7bp, 9bp, and 10bp (Figure S10). All of these expected lengths are much smaller than our cutoff of 500bp and observed means of 6.0kb for san-into-yak, 6.8kb for *yak-*into*-san*, 2.4kb for *tei-*into*-yak,* and 1.1kb for *yak-*into*-tei.* Collectively given the small expected fragment sizes and large number of generations since ancestral polymorphism would be expected to have been lost from the recipient species make it unlikely that the introgression tracts we found are actually ancestral variation.

### Adaptive introgression

Finally, we explored the possibility of introgressions that had become fixed in the recipient population. We looked for introgressions that were fixed in a population within the hybrid zone but not present in allopatric populations. We found no evidence of *san*-into-*yak* introgressions that have completely swept locally to fixation within the hybrid zone on São Tomé. We did identify three regions that had introgressions present in the majority of individuals from the hybrid zone (i.e., with frequencies greater than or equal to 50% Figures S11-S13). The first one, 2R: 19,918,908-19,927,758, is 8.9kb long, is at 50-54.6% frequency, and contains four genes (eEF5, *RpL12, CG13563* and the promoter and 5’ region of *ppk29;* Figure S11). The second introgression, 3L: 6,225,896-6,257,088, is 31.2kb and is at 50% frequency. It contains three genes: two genes with no orthologs in *D. melanogaster* (*FBgn0276401*, and *FBgn0276736)* and *Sif*(Figure S12). The last of the three introgressions at high frequency in the hybrid zone, 3L: 12,187,525-12,209,675, is 22.2kb and is at 59.1-68.2% frequency (depending on the introgression block); it includes three genes: *CG9760*, *Rh7*, and the 3’ portion of *Neurexin IV* (Figure S13). Their multiple breakpoints suggest these introgressions have been segregating within *D. yakuba* long enough for multiple independent recombination events to act.

We did not do a similar analysis at the *D. yakuba/D. teissieri* hybrid zone because we only had three *D. yakuba* individuals for that population.

## DISCUSSION

We found evidence for low levels of introgression between the three species in the *D. yakuba* clade which contains the two only known stable hybrid zones in the *Drosophila* genus. We hypothesized that given ongoing hybridization, we would find high levels of genetic exchange between the two species pairs. Yet, in both hybrid zones, the introgressed regions of the genome are small and generally present at low frequencies. Given the divergence time between the two species and low levels of linkage disequilibrium for the three species (Figure S4), the blocks of shared ancestry are unlikely to represent incomplete lineage sorting and instead reflect introgression. Given their small sizes, low frequencies, and non-consistent enrichment for a type of sequence, it is likely that a majority of the introgressed regions are selectively neutral. Since the results for the two pairs of species differ quantitatively and qualitatively, we discuss them separately.

### yak/san

Int-HMM detected low levels of introgression between *D. yakuba* and *D. santomea,* average levels of introgression are around 0.4% and never exceed 1.2%. Introgressed fragments are generally small, with average sizes of 6.8kb for *yak-*into-*san* and 6kb for san-into-*yak* suggesting recombination has reduced their size over multiple generations and implying that the introgressions are not recent.

Introgression must have occurred through hybrid females (who are fertile), as hybrid males are sterile. Introgression also must have originated in the hybrid zone in an area of secondary contact, likely the midlands of São Tomé, and subsequently spread into other areas.

Notably, *san*-into-*yak* introgressed tracts are not limited to São Tomé. We also found introgressed *D. santomea* alleles in *D. yakuba* lines from other islands in the Gulf of Guinea that are far from the hybrid zone on São Tomé (over 150km). There are two possible explanations for this distribution. First, gene flow within *D. yakuba* between islands in the Gulf of Guinea might be common allowing introgressions to easily spread throughout the archipelago. However, there is some evidence of genotypic and phenotypic differentiation between different *D. yakuba* populations [44]. The second possibility is that *D. santomea* is not endemic to São Tomé but is (or once was) present on other islands in the Gulf of Guinea. Sampling on the islands of Príncipe [39] and Bioko [41,54,77], has not yielded *D. santomea* collections. It is worth noting however, that these collections only inform the current distribution of *D. santomea* and not its historical range. Regardless of the explanation, introgression between these two species pairs is limited and likely to be ancient.

### yak/tei

Also using Int-HMM, we found evidence for introgression between *D. yakuba* and *D. teissieri,* two highly divergent species (Ks~11%). Average levels of introgression are around 0.005% and never exceed 0.025% (i.e., much lower than between *D. yakuba* and *D. santomea*). Most introgressions between these species are small and have low allelic frequencies. The average tract size for *D. yakuba* into *D. teissieri* introgressions was also smaller than between *D. yakuba and D. santomea* (1.1kb and 2.4kb depending on the direction of the introgression). Introgression between these species is asymmetric with higher rates from *D. yakuba* into *D. teissieri than* in the reciprocal direction. The reasons behind this asymmetry are unclear but do not stem from differences in the magnitude of reproductive isolation between the two directions of the cross [57]. Notably, *D. teissieri* flies from the hybrid zone on Bioko had some of the highest levels of introgression of all *D. teissieri* lines. A similar pattern did not hold for *D. yakuba,* as multiple populations of *D. yakuba* show similar levels of introgression. We observed a similar pattern for the *yak/san* pair: *D. yakuba* does not show differences in the magnitude of introgression at different locations.

*Drosophila yakuba* and *D. teissieri* coexist over large swaths of the African continent [53], and thus it is unclear–yet likely–whether other hybrid zones exist. These results are not explained by different rates of migration between *D. yakuba* and *D. teissieri* as they tend to move similar distances [54].

Moreover, we find that introgression from *D. teissieri* is present in all lines of *D. yakuba,* including those from the island of São Tomé. These results might indicate that the colonization of *D. yakuba* to São Tomé occurred after hybridization between *D. yakuba* and *D. teissieri and* the genomes of the colonizing *D. yakuba* flies already contained introgressions from *D. teissieri.* Currently there are no *D. teissieri* on São Tomé and there is no record of *Parinari* (the main substrate of *D. teissieri)* on this island either. We cannot infer the ancestral range of *D. teissieri with* certainty, but it seems unlikely that this species was present on this island. Notably, *tei-into-yak* introgressions do not overlap with the *san*-into-*yak* introgressions we observed. This rules out the possibility that putative *tei*-into-*yak* introgressions in the hybrid zone actually are from *D. santomea.*

### General patterns from both species pairs

Introgression in both species pairs shows that despite strong reproductive isolating barriers, genetic exchange mediated through hybridization is possible in *Drosophila.* In total, over 15 barriers to gene flow have been reported between *D. yakuba, D. santomea,* and *D. teissieri* [46,50,57,78,79]. Females invariably prefer males from the same species, and interspecific matings are rare [46] but see [80]. Interactions between gametes from the three different species can also go awry precluding fertilization [57,77,81]. Hybrid individuals may also be inviable or show behavioral defects. Hybrid males from all interspecific crosses are sterile [38,57]. This wide variety of phenotypes is expected to reduce the amount of introgression between species pairs.

The introgressions we found in all four directions appear to be primarily selectively neutral (but see below). They were generally small, being on average only a few kilobases long, indicating that they had been present in the recipient species for many generations, and recombination had ample time to reduce their size. If these introgressions were ubiquitously beneficial, they would have been swept to fixation across the full geographic range or at least in local populations. We only found a handful of cases where the frequency of introgressed segments exceeded 50%, and most of the introgressions we did find were at low frequencies as expected under neutral drift.

We also addressed whether introgressions were uniformly distributed across the genome. Theoretical models [69] have argued that in species with a hemizygous sex, sex chromosomes should have lower rates of introgression than autosomes. In *Drosophila,* the *X* chromosome plays a large role in reproductive isolation [82-83]. The hemizygosity of the *X* chromosomes means that recessive alleles that are deleterious in an admixed genomic background will manifest their deleterious phenotype in males (whereas an autosomal allele would manifest such a phenotype only when homozygous) [82-86]. Reduced introgression on the *X* chromosome has been found in the *Drosophila simulans* clade [8], in hominids [28,87], and in other mammals [88-90]. Our results support this hypothesis. We saw less introgression on the *X* chromosome in agreement with the large *X* effect [91-93]. Such an effect has also been observed in the *yakuba* species complex [48,94].

Our SELAM results suggest that the percentage of the genome containing introgression can decline quickly after a single generation of introgression reaching the 0.35% seen between *D. santomea* and *D. yakuba* within 100 to 200 generations. This would imply that the introgression was relatively recent. However, the small average introgression sizes that we observe would suggest otherwise. The average tract lengths from the SELAM simulations indicate that thousands of generations are necessary for the average tract size to reach the 6.8kb we see for *yak-*into*-san.* We recognize that a single generation of introgression may not properly model the introgression history within the *D. yakuba* clade, but it provides a rough approximation. Existing models for estimating the magnitude and timing of admixture based on tract sizes do not perform well for old admixture events involving small tracts and when recombination has occurred between admixed fragments [30,95-97]. Our simulations also assume that introgressed alleles are selectively neutral, which is unlikely to be true between such highly diverged species, but modeling the genomic distributions of hybrid incompatibilities and their interactions is beyond the scope of this study.

### Hybridization vs introgression

Our results pose an apparent contradiction. We studied the only two stable hybrid zones known to date in *Drosophila.* Additionally, there seems to have been a recent event of mitochondrial homogenization in these species that can only be explained through hybridization [37,39,56,58]. Yet, we find little introgression between hybridizing species in both cases. How to reconcile the continuous and relatively high level of hybridization with the small amount of observed genomic introgression that seems to be old? *Drosophila yakuba* and *D. teissieri* hybridize in the island of Bioko but the hybrid zone they form is extremely narrow indicating strong selection against the hybrids. Field and laboratory experiments revealed the potential source of this selection: *D. yakuba* prefers open habitats while *D. teissieri* prefers dense forests. Congruently, *D. yakuba* is able to tolerate desiccating conditions, while *D. teissieri* is not well suited for this type of stress. F1 hybrids between these two species show a deleterious combination of traits; while they prefer open habitats like *D. yakuba,* they cannot tolerate osmotic stress. This maladaptive combination of traits might preclude the possibility of these hybrids passing genes to the next generation. Indeed, while hybrids may be sampled on Bioko, no advanced-generation hybrid genotypes have been found [54].

The case of *D. yakuba* and *D. santomea* is more puzzling because the number of hybrids produced in their hybrid zone is much higher [44]. One possible scenario is that there is also strong selection against the hybrids and they simply are not able to reproduce. At least one line of evidence indicates this is the case. Hybrid males in the *yak/san* hybrid zone from one of the directions of the cross migrate towards the top of Pico de São Tomé largely outside of the geographic range of the two parental species. These males are sterile, but hybrid females, which are fertile, might show similar defects. There is evidence that hybrids from both sexes show behavioral defects [98]. The reason for this aberrant migration is unknown, but is likely to be caused by similar behavioral defects.

A second factor that might have diminished the possibility of contemporary gene exchange between these two species is the evolution of postmating prezygotic isolation by reinforcing selection [77]. *Drosophila yakuba* females from the hybrid zone show stronger gametic isolation towards *D. santomea* than females from other regions which might contribute to the reduction in the production of hybrids. Notably reinforced reproductive isolation evolves in just a few generations of experimental sympatry [77,99] and can evolve even in the face of gene flow [100]. Such strengthened reproductive isolation might explain the levels of introgressions we observe in the *yak/san* hybrid zone: a combination of stronger prezygotic isolation (evolved via reinforcement) and strong selection against F1 hybrids, would lead to high rates of hybridization and little introgression. The observed levels of introgression might be a relic of even higher levels of hybridization before reinforced gametic isolation was in place.

### Adaptive introgression

The vast majority of introgressions were at low frequency, but we tested whether any of the alleles identified in our screen showed evidence of adaptive introgression. We find potential evidence for three alleles that have increased in frequency locally (i.e., in the São Tomé hybrid zone; [101]) after crossing the species boundary from *D. santomea* or from *D. teissieri* into *D. yakuba.* It is worth noting that given their size and the rather large number of breakpoints, these introgressions are unlikely to have entered *D. yakuba* in the recent past.

We found three *san*-into-*yak* introgressions that increased to high frequency in the hybrid zone. The first one, 2R: 19,918,908-19,927,758, contains four genes: *eIF5, RpL2, CG13563,* and the 5’ portion of *ppk29.* The most intriguing of these candidates is *ppk29* because the gene is involved in intraspecific male-male aggression in *D. melanogaster* [102], and larval social behavior also in *D. melanogaster* [103]. *ppk29* is also necessary for promoting courtship to females [104] and inhibit courtship towards males [104].

The second *san*-into-*yak* introgression, 3L: 6,225,896-6,257,088, contains a portion of the intron of *Sif* and two genes with no known orthologs in *D. melanogaster* ( *GE28246, GE28581). Sif* is differentially expressed after light stimulation, and functional analyses in *D. melanogaster* show a strong effect of the gene on the regulation of circadian rhythm [105]. Surprisingly, knockdowns of *Sif* in projection neurons result in changes in odor-guided behavior: mutants are more attracted to fermenting fruit [106,107]. Other effects of the gene show that it is implicated in resistance to fungal pathogens [108].

The final *san*-into-*yak* introgression at high frequency in the hybrid zone, 3L:12,187,525-12,209,675, contains three genes: *Nrx-IV*, *CG9760*, *and Rh7. Nrx-IV* human orthologs (*CHRNA5*, *CHRNA* 7) have been implicated in alcohol dependence and natural intronic polymorphism segregating within *D. melanogaster* has been associated with resistance to alcohol [109]. It has also been associated with resistance to fungal pathogens [108]. *Rh7 is* a rhodopsin that has been implicated in fly vision and regulation of circadian rhythm and light perception [110].

These three introgressions contain genes that could potentially be involved in adaptation, but we cannot yet claim that these alleles are adaptively introgressed. More generally, we do not yet know whether any of these genes leads to interspecific trait differences. Only careful physiological and functional study of potentially adaptive phenotypes in the three pure species and the admixed individuals will reveal to what extent these introgressed regions are truly adaptive.

### Caveats

Our approach is not devoid of caveats. First, we sequenced individuals from isofemale lines. These lines are derived from a single inseminated female and over time their progeny will lose heterozygosity quickly [111,112]. This means that our assessment of gene exchange might be warped by this inbreeding step. On one hand, inbreeding leads to homozygote flies and deleterious introgressions will be more likely to be lost from the sample. On the other hand, if inbred flies are introgression carriers and homozygous, we will be able to detect introgression in a more reliable manner. A systematic sequencing of flies directly collected from the field will reveal whether the use of isofemale lines does indeed mislead the quantification of introgression.

Second, all our analyses were done using a *D. yakuba* reference genome. The greater divergence between *D. yakuba* and *D. teissieri* may also result in less ability to map *D. teissieri* reads in less conserved regions such as intergenic sequence thus causing us to miss introgressions.

Third, beneficial alleles would likely go to fixation quickly and would be undetectable by our approach since both species would have the same allele. Additionally, such adaptive introgressions that have swept to fixation could cause our method to misidentify the direction of introgression. We find evidence for three potential cases of adaptive introgressions (not fixed but at high frequency in the hybrid zone) but we do not believe that such instances are common. Most genes are unlikely to be adaptive in a new genomic environment [113-115]. Since linkage disequilibrium declines precipitously on the order of a few hundred base pairs in the *Drosophila* species we are working with and the minimum size for introgression tracts we are reporting is 500bp, misidentified adaptive introgressions should be very rare in our dataset. A demographic assessment of the timing and likely evolutionary history of these introgressions might help resolve the issue.

Fourth, we selected markers that were fixed in the donor species with an allele frequency difference between species greater than 0.3. This cutoff was chosen because the closer the allele frequency difference is to zero, the less information the marker contains. However, in practice this means that we were unable to detect introgressions that had increased in the recipient species to frequencies greater than 0.7. Given the distribution of allele frequencies we observed, it seems unlikely that there are many introgressions at such high frequencies, but we would be unable to detect those that existed. Given the small differences between species, such introgressions could be difficult to detect for any method, particularly one based on allele frequencies.

Our approach is also unable to detect regions of the genome with bidirectional introgression. However, given the low levels of introgression we observe (< 1%) and the small sizes of introgression tracts, such overlaps are expected to be rare. A final, and related potential caveat would be that introgressions in *D. yakuba* were attributed separately to both *D. santomea* and *D. teissieri.* However, there is a little overlap between *san*- into*-yak* and *tei-*into*-yak* introgressions with only two lines (1_5 and 1_7) each having the same overlap which spans just 2,439bp.

### Conclusions

Hybridization is common across the tree of life. Hundreds of hybrid zones have been described over the last 150 years [116-119] but until recently identifying the segments of the genome that had crossed species boundaries was all but impossible. Genome sequencing has been able to identify multiple cases of recent admixture and introgression [19,120-123]. Large pieces of the genome in modern humans originated from other hominids [29,31,115,120,124-126]. Hybridization in plants is rampant and has had deep implications in their diversification [127-130]. Systematic surveys in birds also have provided evidence that hybridization and introgression might be frequent but not ubiquitous processes ([131-133] reviewed in [134,135]). Overall, there is strong evidence that hybridization is common across animals [13,136,137], and there are clues that introgression might not be rare [19,138]. Significant progress has been made to detect introgression when migration is recent [24,139]. Ancient introgression remains a largely underexplored question because identifying small introgressions is challenging (but see [28,33,87]). We provide a general method to detect introgression that does not depend on having phased data or on identifying pure individuals beforehand. Additionally, our method reliably identifies introgressions even when introgression is rare. We have mapped such introgressions between two pairs of species in the *Drosophila yakuba* clade and found minimal genomic introgression despite the existence of stable hybrid zones and ongoing hybridization. Our results indicate that hybridization does not necessarily imply gene flow between species. The two species pairs in the *yakuba* clade likely represent the later stages of the speciation process and similar mapping efforts are necessary in species pairs that are less diverged to better understand how divergence time affects rates of hybridization and subsequent genomic introgression.

## METHODS

### Genome sequencing

#### Fly Collection

*Drosophila* lines were collected in the islands of São Tomé and Bioko. To collect flies, we set up banana traps in plastic bottles hanging from trees. Flies were aspirated from the traps without anesthesia using a putter [140,141]. Flies were then sorted by sex and species. Males were kept in RNAlater; females were individually placed in 10mL plastic vials with instant potato food (Carolina Biologicals, Burlington, NC). Propionic acid and a pupation substrate (Kimwipes Delicate Tasks, Irving TX) were added to each vial. We collected the progeny from each female and established isofemale lines [140]. All collected stocks and populations were reared on standard cornmeal/Karo/agar medium at 24°C under a 12 h light/dark cycle. The taxonomical identification was confirmed by performing crosses with tester stocks (*D. santomea:* sanSYN2005*; D.yakuba:* Täi18; *D. teissieri:* Selinda). Other additional lines were donated by J.A. Coyne and are listed in Table S1. Figure S14 indicates the number of fly lines used in this study from each geographic location.

#### DNA extraction

DNA was extracted from single female flies using the QIAamp DNA Micro Kit (Qiagen, Chatsworth, CA, USA) kit. We followed the manufacturer’s instruction using cut pipette tips to avoid shearing the DNA. This protocol yields on average ~40ng (range: 23ng-50ng) of DNA per fly per extraction.

#### Library Construction

For short read sequencing, we constructed libraries following two methods. 54 libraries were built using the TrueSeq Kappa protocol (University of North Carolina, Chapel Hill). For these libraries, ~10 ug of DNA was sonicated with a Covaris S220 to a mean fragment size of 160 bp (range = 120–200 bp) with the program: 10% duty cycle; intensity 5; 100 cycles per burst; 6 cycles of 60 seconds in frequency sweeping mode. The other 12 libraries were built using Nextera kits at the sequencing facility of the University of Illinois, Urbana-Champaign. For these libraries, DNA was fragmented using Nextera kits which uses proprietary transposases to fragment DNA. Libraries were built following standard protocols [72].

#### Sequencing

We sequenced all libraries on Illumina HiSeq 2000 machines with v3.0 chemistry following the manufacturer’s instructions. Table S1 indicates the sequencing type (single-end or paired-end), and coverage for each library. Libraries were pooled prior to sequencing and 6 libraries were sequenced per lane. To assess the quality of the individual reads, the initial data was analyzed using the HiSeq Control Software 2.0.5 in combination with RTA 1.17.20.0 (real time analysis) performed the initial image analysis and base calling. Run statistics for each FASTQ file was generated with CASAVA-1.8.2. Resulting reads ranged from 100bp or 150bp and the target average coverage for each line was 30X. The coverages for each line are shown in Table S1. We obtained *D. yakuba* sequences for 20 previously sequenced lines (10 from Cameroon and 10 from Kenya) from [142] (Table S1).

#### Read mapping and variant calling

Reads were mapped to the *D. yakuba* genome version 1.04 [143] using bwa version 0.7.12 [144]. Bam files were merged using Samtools version 0.1.19 [145]. Indels were identified and reads were locally remapped in the merged bam files using the GATK version 3.2-2 RealignerTargetCreator and IndelRealigner functions [146,147]. SNP genotyping was done using GATK UnifiedGenotyper with the parameter het = 0.01. The following filters were applied to the resulting vcf file: QD = 2.0, FS_filter = 60.0, MQ_filter = 30.0, MQ_Rank_Sum_filter = -12.5, and Read_Pos_Rank_Sum_filter = -8.0. Sites were excluded if the coverage was less than 5 or greater than the 99^th^ quantile of the genomic coverage distribution for the given line or if the SNP failed to pass one of the GATK filters.

#### PCA

We used Principal Component Analysis (PCA) to assess the partition of genetic variation within the *yakuba* species complex. PCA transforms a set of possibly correlated variables into a reduced set of orthogonal variables. Sampled individuals are then projected in a two dimensional space where the axes are the new uncorrelated variables, or principal components. We used the R package *adegenet* [148] to run separate PCA analyses for the *X*chromosome and autosomes and plotted the first five principal components. For all PCA, we calculate the amount of variance explained by each principal component.

#### ABBA – BABA tests

To calculate interspecific gene flow, we first calculated historical levels of gene flow between different species pairs in the *yakuba* clade with the ABBA-BABA/D statistic [31,33,71,149] using a perl script. The ABBA-BABA test compares patterns of ancestral (A) and derived (B) alleles between four taxa. In the absence of gene flow, one expects to find equal numbers of sites for each pattern. However, gene flow from the third to the second population can lead to an excess of the ABBA pattern with respect to the BABA pattern, which is what the D statistic tests for. We used allele frequencies within the specified populations (i.e., putative recipient, putative donor, outgroup) as the ABBA and BABA counts following [33,71]. We assessed the significance of ABBA-BABA test statistics using the commonly employed method of weighted block jackknifing with 100kb windows [150]. Briefly, this systematically removes consecutive non-overlapping portions of the genome (100kb blocks in this case) and re-estimates the statistic of interest to generate a confidence interval around it. We also estimated the proportion of the genome that was introgressed with the f_d_ statistic [71]. f_d_ compares the observed difference between the ABBA and BABA counts to the expected difference when the entire genome is introgressed.

##### Treemix

We used *TreeMix* [32] to investigate the relationship between species and to look for evidence of historic gene flow. *TreeMix* estimates the most likely evolutionary history in terms of splits and mixtures of a group of populations by estimating levels of genetic drift. The analysis is done in two steps. First, it estimates the relationships between sampled populations and estimates the most likely maximum likelihood phylogeny. Second, it compares the covariance structure modeled by this dendrogram to the observed genetic covariance between populations. The user then specifies the number of admixed events. If a pair of populations is more closely related than expected by the strictly bifurcating tree, then maximum likelihood comparisons will suggest an admixture event in the history of those populations. We ran *Treemix* separately for the *X* chromosome and the autosomes. The program was run with 6 populations. We assigned only one population for *D. santomea* due to its limited range and one population for *D. teissieri* since we only had 13 lines even though they originated from multiple geographic locations. *Drosophila yakuba* was partitioned into four populations: an ‘islands’ population that included lines from Príncipe and Bioko, an ‘africa’ population containing the mainland African lines from the Ivory Coast, Cameroon, and Kenya, a ‘low_st’ population for the lowlands of São Tomé, and a ‘hz_st’ population for lines from the hybrid zone on São Tomé. We ran Treemix for each dataset with m=0 through 5 migration edges and determined the most likely number of migration events (n) by doing a log likelihood test comparing the runs with m = n and m = n - 1. The most likely value of m was the largest value of n before the test was no longer significant at a 0.05 level.

### Hidden Markov model

#### Selecting markers for the hidden Markov model

*Treemix* and the ABBA-BABA D statistic can be used to assess whether genetic exchange has occurred between species (and populations), but they do not identify specific introgressed regions of the genome. We identified introgressed regions in all individuals from all three species using a hidden Markov model. The hidden Markov model determined the most likely genotype (the hidden state) for each SNP we used as a genomic marker. When looking for introgression from one species into another, we would ideally have allopatric and sympatric populations of the recipient species. Fixed differences between a putative donor species and an allopatric population of the recipient species are informative markers that can be used to help identify introgression. However, allopatric populations do not exist for all of the species pairs in the *yakuba* clade. The ranges of *D. yakuba* and *D. teissieri* overlap extensively, and no *D. santomea* flies live more than a few miles from the hybrid zone with *D. yakuba.* We were, therefore, unable to identify markers that were definitively associated with the recipient or donor species. Instead, we selected SNPs to be markers where the donor species was monomorphic and the allele frequency differences between the two species was greater than or equal to 30%. 30% was chosen because the smaller the allele frequency difference between species, the less informative an individual site is for identifying introgression and the noisier the data becomes as neutral mutations that are segregating at a low frequency in the recipient species are also included. Furthermore, we required that every individual in the donor species and at least one individual in the recipient species had a called genotype. We also excluded sites where more than 80% of the individuals with a genotype call from GATK were heterozygous as the high frequency of heterozygotes likely indicated mapping error. For the *D. santomea* into *D. yakuba* analysis, we excluded four lines from *the D. santomea* donor population that were more similar to *D. yakuba* for both PC1 and PC2 for the autosomal PCA analysis: sanST07, BS14, C550_39, and san_Field3. They were excluded since they were expected to have higher levels of introgression which would reduce the number of markers because of the requirement that the donor species be monomorphic.

#### Transition Probabilities

Transition probabilities determine how likely the HMM is to move between the hidden states. The transition probabilities use two starting probabilities, a for transitions between non-error states and *a_e_* for transitions between error states. Separate transition probabilities and starting probabilities are calculated for each marker and depend on the distance to the next marker. We modeled the starting probabilities as Poisson variables with the parameter equal to the per site recombination rate, c, times the distance between the two sites. The parameter for *a_e_* also used a multiplier m. The multiplier ensured that it was somewhat easier to stay in an error state. For the *D. santomea* and *D. yakuba* introgression analysis, we used c = l0^−9^ and m = 25,000. Base transition probabilities for non-error (*a*) and error sites (*a_e_*) were based on the distance between the two neighboring markers (whose positions are denoted as x_i_ and x_i-1_):

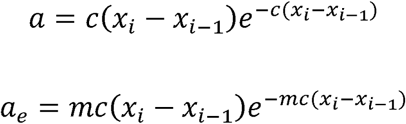

The transition probability matrix was constructed so the model was more likely to transition from a non-error state to the same non-error state. Transitioning from a non-error state to the corresponding error state was impossible (e.g. from homo_r to homo_r_e_). The probability of transitioning from an error state to another error state was small to ensure the model would quickly leave the error states. The transition probability matrix represents the probability of transferring from the state denoted by the row to that of the column.

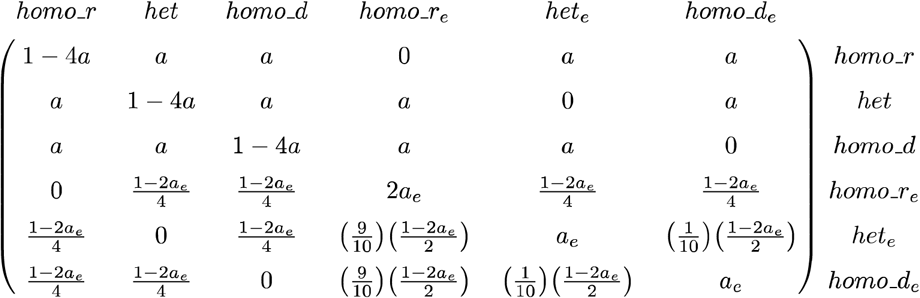

#### Emission Probabilities

The HMM only used biallelic sites, and the two alleles are expressed as *a* and *b.* Let k represent the number of copies of the a allele at a given site, and the probability of seeing k copies without sequencing error is:

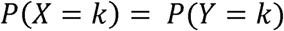

Where Y is a random variable denoting the number of *a* alleles present in the DNA fragments for that site chosen for sequencing. These reads are then sampled from to determine which reads are subjected to sequencing error. Define two random variables A and B as respectively the number of a and b alleles resulting from sequencing error. For a total coverage of n, P(X = k) can be written as:

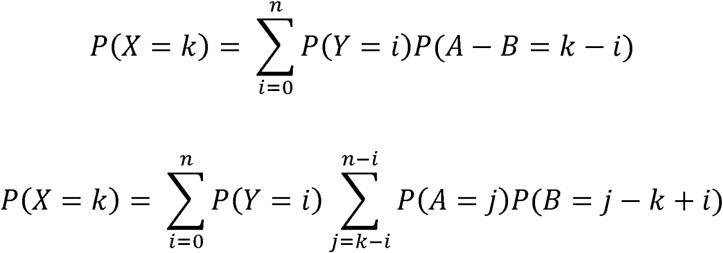

Using binomial probabilities for Y, A, and B, equation (3) can be expanded to:

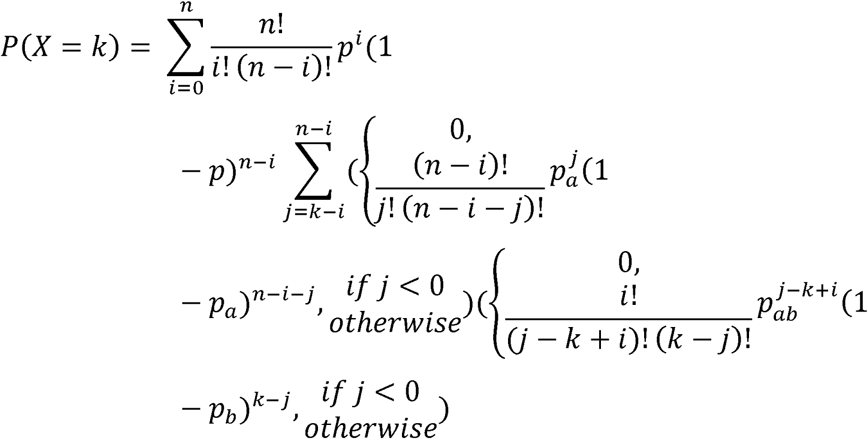

where p is the probability of sampling an *a* allele from the sequenced DNA fragments, pa is the per base probability of obtaining an a allele via sequencing, and likewise, p_b_ is the sequencing error probability for a b allele. Simplifying further yields:

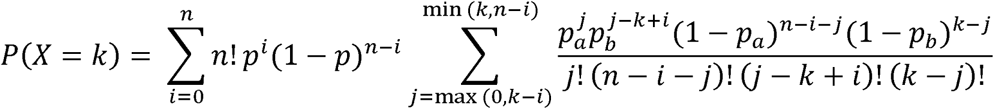

Assuming all alleles are equally likely through sequencing error, we define p_ab_ = p_a_ = p_b_, and equation (5) simplifies further to yield the per base emission probability:

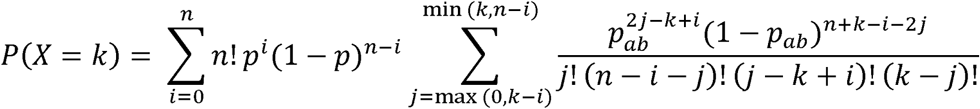

#### Identifying introgression tracts

The HMM determined the most probable genotype for each marker in each individual. We defined tracts as contiguous markers with the same genotype, and a series of seven filters were then applied to the tracks in the order listed below. In the descriptions that follow, “het” refers to a heterozygous tract, “homo_d” to a tract that is homozygous for donor species alleles, “homo_r” to a track that is homozygous for recipient species alleles, and an introgression tract can be either a het or homo_d tract. Introgression SNPs are defined as those within the tract where the HMM probability for an introgression state (het or homo_d) was ≥ 50%. In subsequent analyses we treated homozygous and heterozygous introgression tracts equally because the sequenced lines were isofemale and because the filtering rules combined adjacent homozygous and heterozygous tracts. We applied the following filters:

1. Merge het and homo_d tracts – Adjacent het and homo_d tracts represented a single introgression that the HMM assigned to multiple genotypes. In such cases, all adjacent het and homo_d tracts were combined into a single tract with the genotype determined by the genotype that had the most introgression SNPs. Ties were assigned to het.
2. Remove small het and homo_d tracts in high error regions – Some genomic regions were characterized by multiple rapid transitions between states. These regions could result from mapping error, incorrectly assembled genomic regions, and or ancient introgressions that had been greatly reduced by recombination. Most of the tracts in such regions were small and ended up in error states. When a het or homo_d tract was found in the middle of one of these regions, we deemed it best to treat it as an error state, het and homo_d tracts were assigned to their corresponding error state if they had less than 15 introgression snps and the number of their introgression snps divided by the sum of SNPs from all adjacent contiguous blocks of error tracts was less than 3.
3. Remove error blocks – homo_r tracks were sometimes broken up by either a single error tract or short blocks of error tracts. Such cases likely resulted from mapping error, incorrectly assembled regions of the genome, or new mutations in the recipient species. In such cases, the contiguous blocks of error tracts bounded on both sides by homo_r tracts were reassigned to homo_r.
4. Merge small error tracts – Similarly to filter 3, introgression tracks could also be broken up by error tracts. Contiguous blocks of error tracts bounded on both sides by introgression tracts were all combined and assigned whichever of the het or homo_d tracts had the most introgression SNPs. Ties were assigned to het.
5. Convert error tracts to homo_r – After the first four filters were applied, any remaining error tracts were changed to homo_r.
6. Remove small homo_r tracts – After the error tracts were removed, some of the larger introgression tracts were broken up by small homo_r tracts. Such cases could result from mapping error, incorrectly assembled genomic regions, or newly arisen mutants in the recipient species. Since most of the intervening homo_r tracts were small and on the order of a single SNP or less than 100bp, they were unlikely to represent multiple crossover events. We, therefore, combined the introgression and homo_r tracts. homo_r tracts with less than 5 total SNPs bounded on both sides by introgression tracts with at least 10 total SNPs each were all combined into a single tract. The genotype was determined by whichever of the het or homo_d tracts had the most total SNPs. Ties were assigned to het.
7. Merge het and homo_d tracts – The first filter was run a second time.

Figure S15A contains a graphical representative of the process of a representative region for a *san*-into-*yak* introgression for the line SãoTomé_city_14_26. The panels show the markers, the allele coverages at those markers, the genotype probabilities returned by the HMM, and the unfiltered and filtered tracts. Figure S15B shows the tracts for the same region for all 56 *D. yakuba* lines. Software and documentation for Int-HMM are available at https://github.com/dturissini/Int-HMM.

#### Simulating introgressions

We ran the HMM on simulated introgressions to test its accuracy. The introgressions were simulated with a perl script that processed the vcf file containing the genotyping results for the all three species. Introgressions were simulated with sizes ranging from 100b to 100kb with random distances between them uniformly distributed between 25kb and 75kb. Each introgressed region had a 50% chance of being either heterozygous or homozygous. Alleles at each site were determined by sampling from the alleles present in the donor species within introgressed regions and from the recipient species elsewhere with probabilities determined by population level allele frequencies. Per site coverages from sequencing data vary, and we modeled this by randomly sampling the coverage at each site from a uniform distribution with values ranging from 10 to 25. At heterozygous sites, the relative allele coverages were determined by binomial sampling. We then created markers for individuals and ran the HMM analysis for a single individual for three introgression scenarios: *D. yakuba* into *D. santomea, D. santomea* into *D. yakuba, D.yakuba* into *D. teissieri,* and *D. teissieri* into *D. yakuba.*

#### Comparing introgressions between the *X* chromosome and autosomes

*Drosophila* males are heterogametic, and the *X* chromosome is commonly involved in hybrid breakdown [69,93,151]. Introgressions may be more easily purged by selection when *X* linked rather than autosomal. We determined if introgressions on the *X* chromosome were underrepresented with respect to the autosomes by comparing their cumulative introgression lengths using randomization tests. In a purely neutral scenario, introgressions will be uniformly distributed across chromosomes. Since the *X*-chromosome encompasses 18% of the *Drosophila* assembled genome, any significant downward deviations from this number might indicate selection against introgression in the *X*-chromosome. For each of the four introgression directions (two reciprocal directions in two species pairs), we compared the observed proportion of introgressed sequence on the *X* to a distribution of proportions obtained by reshuffling the introgressed tracts randomly through the genome from all individuals 10,000 times without replacement. Each iteration of the resampling calculated the percentage of introgressed material that was on the *X*-chromosome (given this random-neutral assortment) given that each fragment had a 18.41% chance of being assigned to *X* chromosome. P values for the hypothesis that introgression tracts were underrepresented on *X*-chromosomes were obtained by dividing the number of resampled proportions that were lower than the observed value by 10,000.

#### Expected patterns from ancestral variation

Distinguishing introgression from ancestral polymorphism is crucial to understand the causes of shared genetic variation. We assessed whether our purported introgressions could instead be the result of ancestral variation that was still segregating in the recipient species using two approaches. First, we calculated the expected time for an allele segregating in the ancestral population to be either fixed [152]:

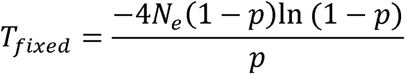

or lost [152]:

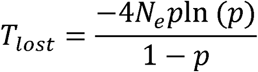

Where *p* is the initial allele frequency, and *N_e_* is the ancestral effective population size. We used three values of *N_e_:* 10^4^,10^5^, and 10^6^.

We also looked into the expected lengths of ancestral haplotype blocks still segregating in the recipient species. We generated a distribution of expected fragment lengths by sampling from the expected distribution of one-sided distances to a recombination event using the simplified probability density function for equation (3) from [153]. We sampled from the distribution twice and added the two lengths to obtain the expected fragment length. We assumed a 50Mb chromosome, a divergence time of one million years, and effective population sizes (Ne) of 10^4^ and 10^6^. We also used generation times of 14, 21, and 28 days to cover a range of lengths around the estimated generation length of 24 days for *Drosophila melanogaster* [36]. This procedure was repeated one million times to generate a distribution of expected fragment lengths. Genetic map lengths were converted to base pairs by dividing by a per base recombination rate of r = 1.2 × 10^-8^ for *D. melanogaster* [154].

#### SELAM simulations

We obtained rough estimates of the age of introgression by comparing the length of our observed introgressions to those generated by simulations from the program SELAM [74]. SELAM is a forward time simulation program capable of modeling admixture at a genomic scale with recombination. We ran SELAM assuming census population sizes of 10,000 for both species. All sites were assumed to be neutral, and the simulations had three chromosomes with lengths of 1, 1, and 0.75 morgans to represent the *D. yakuba* chromosomes 2, 3, and *X* respectively. In the absence of a recombination map for *D. yakuba,* recombination rates were assumed to be uniform across each chromosome. We modeled a single generation pulse of introgression and recorded the introgression tracks from 50 sampled individuals every 100 generations for 10,000 generations. We ran the program five times each for four initial migration rates of 0.1, 0.01, 0.001, and 0.0001.

#### Linkage Disequilibrium

We calculated r^2^ as a metric of linkage disequilibrium (LD). This measurement had two goals. First, we calculated the amount of LD to verify if we could use published methods to detect introgression that explicitly relies on this measurement (e.g., [26]). A fast decay of LD, precludes the possibility of using these methods because even when LD based methods can detect admixture LD, these methods often rely on calculating background LD as well. Second, we used it to confirm that the introgressed tracts we identified were not ancestral haplotypes that were still present in both the donor and recipient species. Low levels of LD argue against this possibility. LD was measured for each of the three species using PLINK [73]. LD was measured separately for the *X* chromosome and the autosomes. We only used SNPs where the per site coverage was between 5× and the 99^th^ quantile of the genomic distribution of coverages for a given individual. Also, at least half of the individuals needed to have had a called genotype in the VCF file produced by GATK. PLINK was run with the following parameters: plink --noweb - --r2 -maf .05 --ld-window 999999 --ld-window-kb 25 --ld-window-r2 0.

#### Measuring proportions of F1 hybrids in the field

The proportion of males collected in the field that are F1 hybrids is a proxy for the current rate of hybridization. For *D. yakuba* and *D. santomea,* we used estimates of F1 hybridization from [44,45,49]. We also added estimates of F1 hybridization from a new field collection in 2016. We used estimates for F1 hybridization between *D. yakuba* and *D. teissieri* from 2009 and 2013 as reported in [54].

#### Data availability

Fastq files are available at SRA (Accession number: TBD). All analytical code has been deposited at Dryad (Accession number: TBD). Code for Int-HMM is available at https://github.com/dturissini/Int-HMM.

#### Ethics statement

*Drosophila* flies were collected in São Tomé é Principe with the permission of the Direccão Geral do Ambiente do São Tomé é Principe. Permits to collect in Bioko were issued by the Universidad Nacional de Guinea Ecuatorial (UNGE). Live flies imported to USA as stated in the USDA permit: P526P-15-02964.

## ACKNOWLEDGEMENTS

We would like to thank members of the Jones, Burch, Vision, and Matute labs for helpful feedback and UNC for startup funding. A. Comeault, B. Cooper, C.D. Jones, K.L. Gordon, R. Marquez, C. Martin and M. Turelli gave us useful comments. We do not have any conflicts of interest.

## SUPPLEMENTARY FIGURES

**Figure S1.**
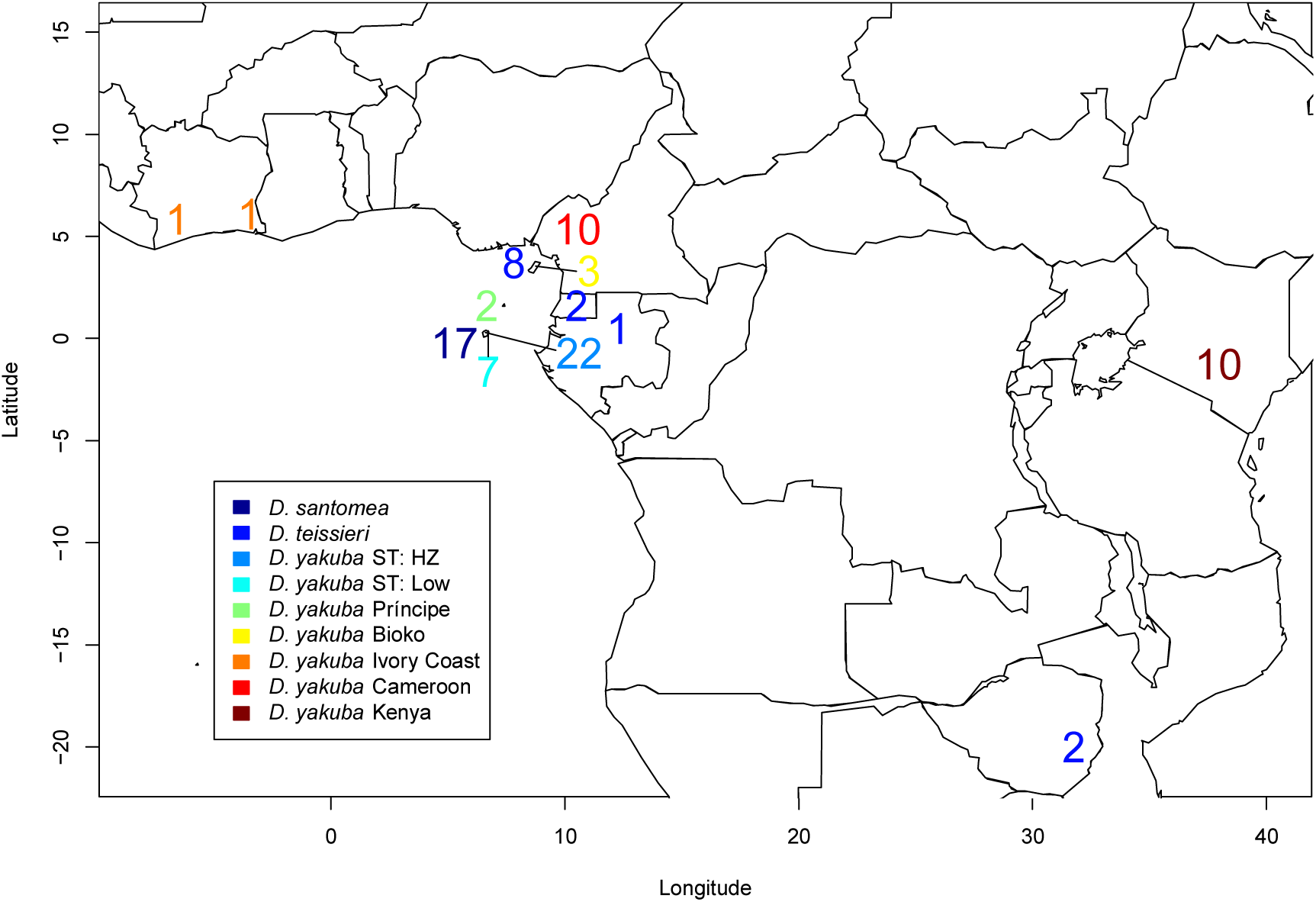
Principle Component Analysis (PCA) for the *D. yakuba clade*. Principle component results for PC1 and PC2 for the *D. yakuba* clade. **A)** Autosomes. **B)** *X* chromosome.

**Figure S2.**
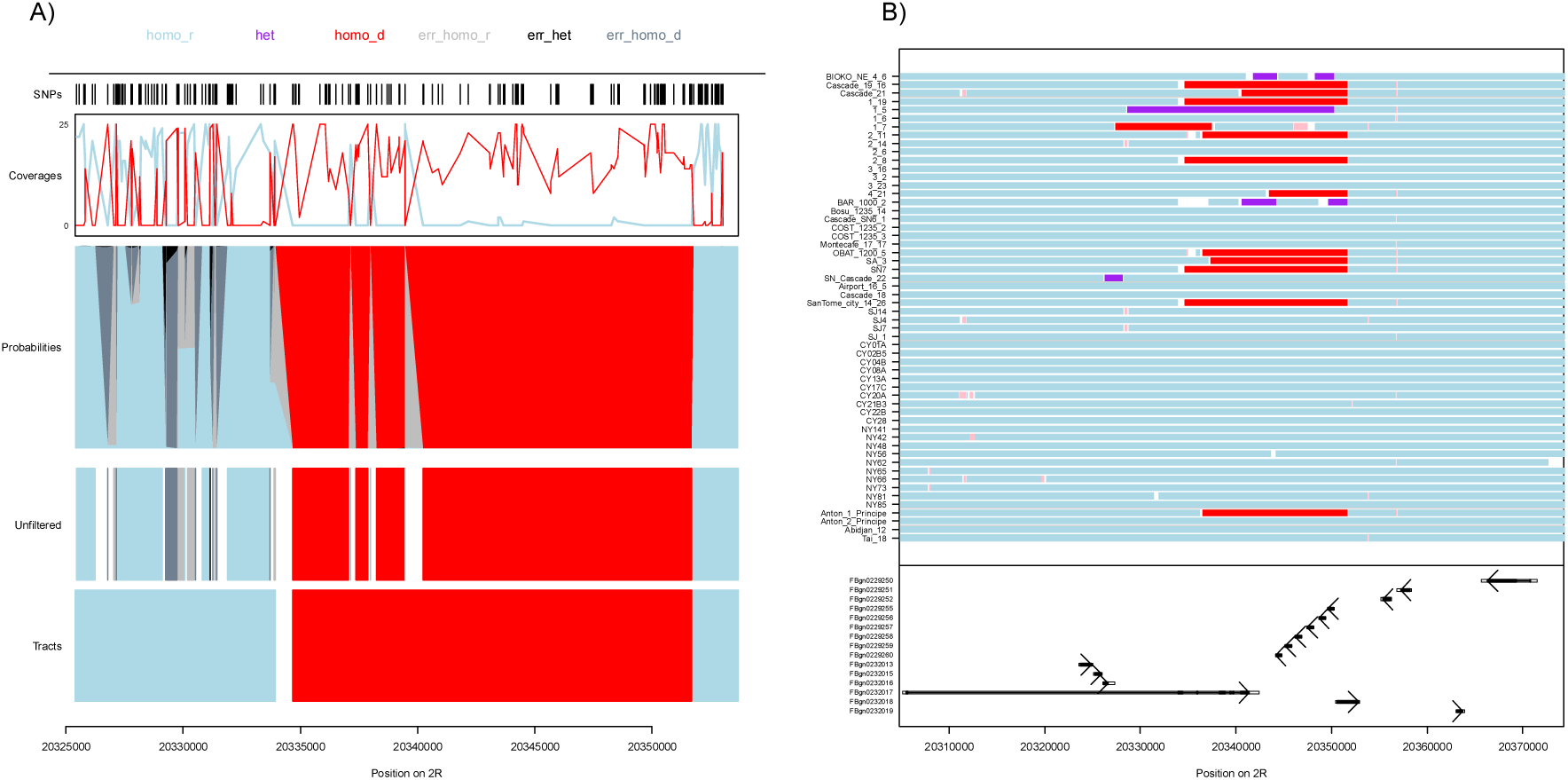
*Treemixresrxlts* for the *X* chromosome, *X* chromosome *Treemix* trees with 0 to 3 migration edges (the most likely value of m=4, Figure 1A). *Drosophila yakuba* was split into four populations: “africa” (Cameroon, Kenya, Ivory Coast), “islands” (Príncipe and Bioko), “low_st” (lowlands of São Tomé), and “hz_st” (hybrid zone on São Tomé). The P value was calculated for a tree with m migration edges by taking a log-likelihood ratio test using the likelihoods for the threes with m and m-1 migration edges. **A)** m=0. **B)** m=1. **C)** m=2. **D)** m=3.

**Figure S3.**
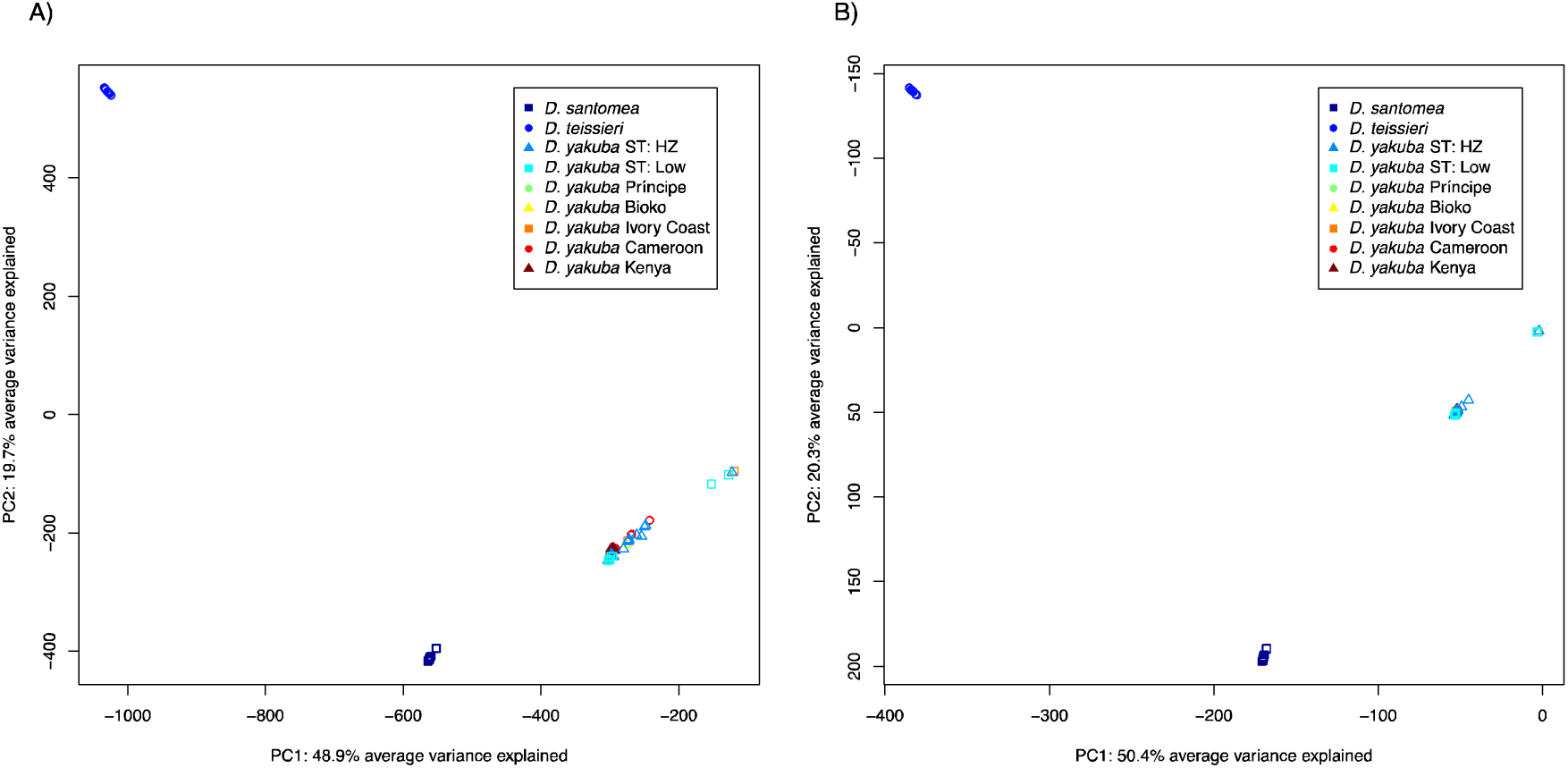
Treemix results for autosomes. Autosomal Treemix trees with 0 to 3 migration edges (the most likely value of m=4, Figure 1B). *Drosophila yakuba* was split into four populations: “africa” (Cameroon, Kenya, Ivory Coast), “islands” (Príncipe and Bioko), “low_st” (lowlands of São Tomé), and “hz_st” (hybrid zone on São Tomé). The P value was calculated for a tree with m migration edges by taking a log-likelihood ratio test using the likelihoods for the threes with m and m-1 migration edges. **A)** m=0. **B)** m=1. **C)** m=2. **D)** m=3.

**Figure S4.**
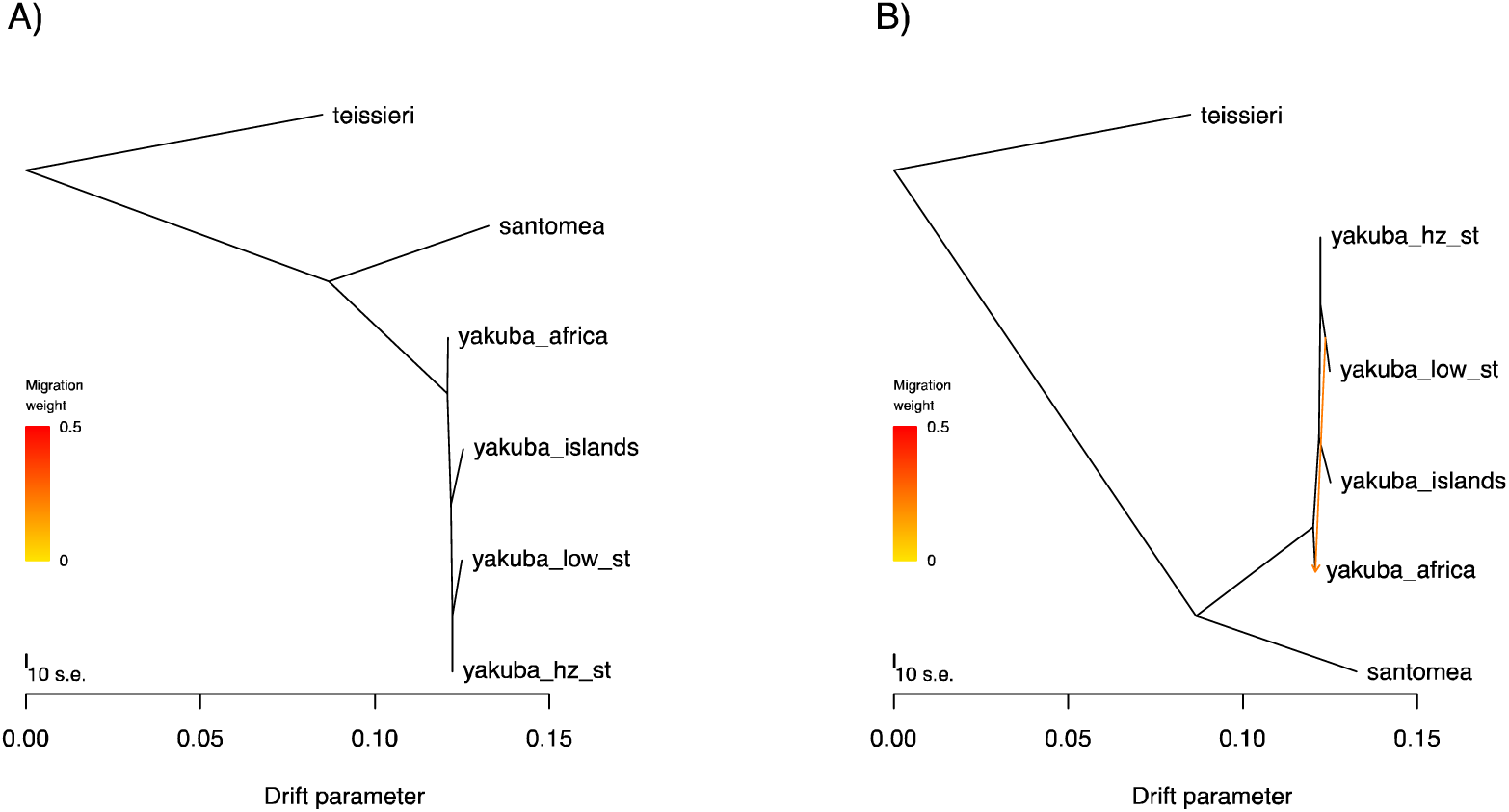
Linkage disequilibrium decays on the order of a few hundred base pairs in all three species in the *D. yakuba* clade. Average LD as measured by r^2^ between pairs of SNPs with distances binned every 10bp. r^2^ was averaged separately for the autosomes (red) and *X*chromosome (blue). **A)** *D. yakuba.* **B)** *D. santomea.* **C)** *D. teissieri.*

**Figure S5.**
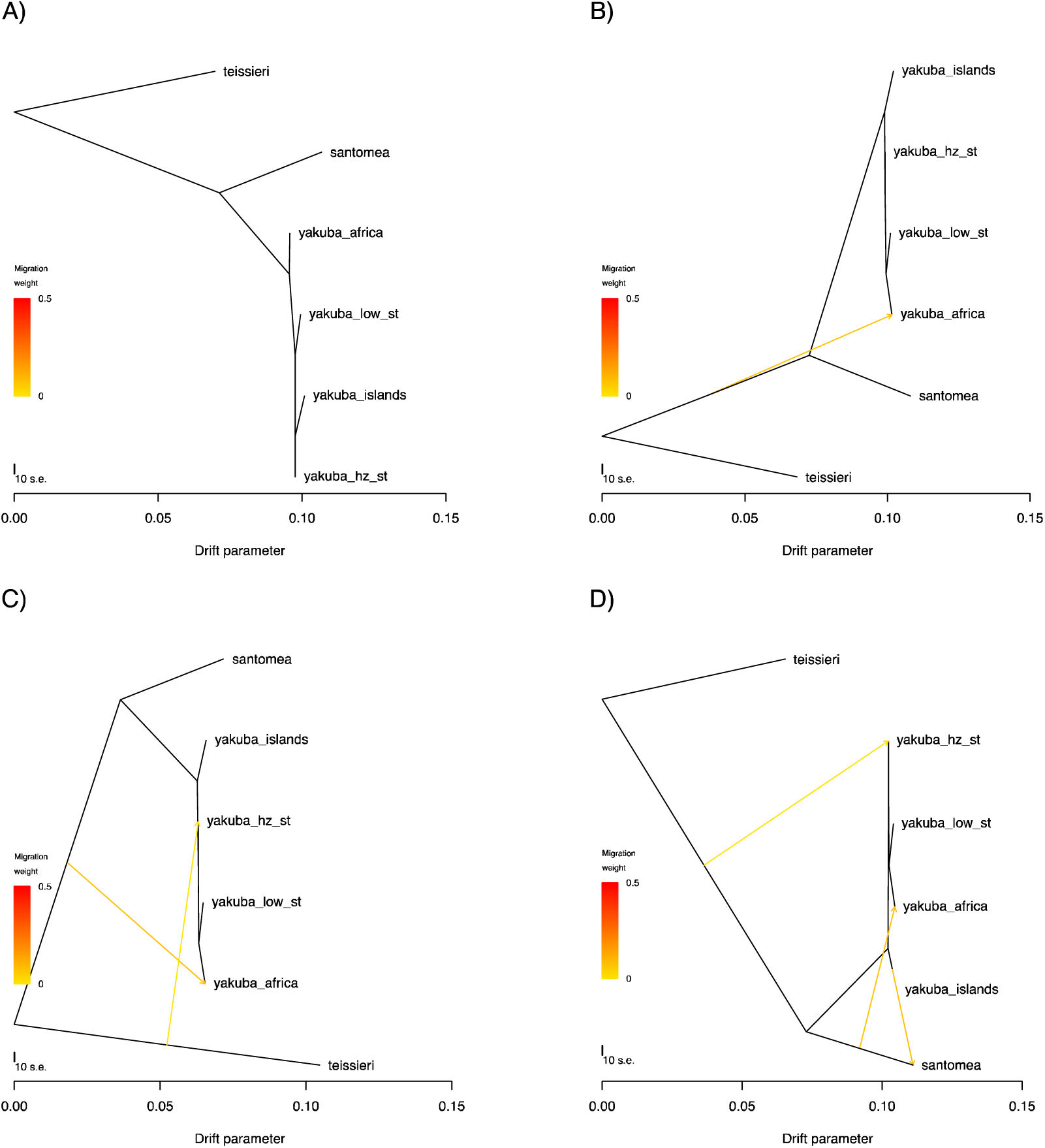
Large introgression in the line Anton_2_Principe. 959kb introgression from *D. santomea* into the *D. yakuba* line Anton_2_Principe collected from the island of Príncipe that is significantly larger than the second biggest introgression (120kb). Red denotes homozygous *D. santomea* tracts, light blue tracts are homozygous *D. yakuba,* and purple tracts are heterozygous (inferred using Int-HMM). The lower panel shows genes in the genomic region on 3R with the arrows denoting the direction of transcription.

**Figure S6.**
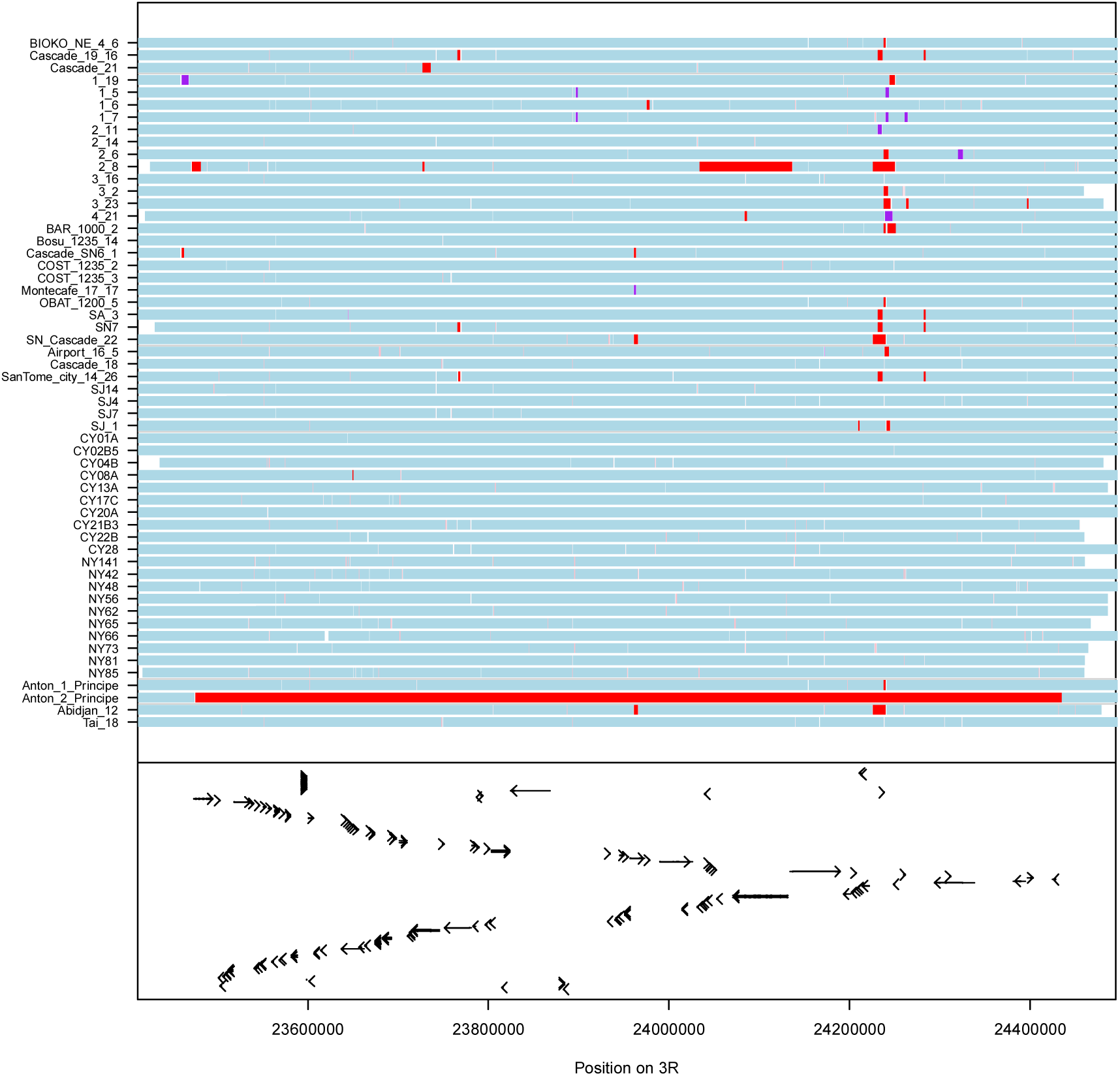
Less introgression on the *X* chromosome than on the autosomes for all directions of gene flow. Percentage of introgressed sequence on the *X* chromosome for 10,000 iterations of resampling without replacement in a neutral scenario where introgressions are uniformly distributed across the genome. P values were obtained by dividing the number of resampled proportions that were lower than the observed value by 10,000. The red line indicates the observed percentage of introgressed sequence on the *X* chromosome, and the blue line is the average percentage from the 10,000 resampling iterations. **A)** *san-*into*-yak.* **B)** *yak-*into*-san.* **C)** *tei-*into*-yak* **D)** *yak-*into*-tei.*

**Figure S7.**
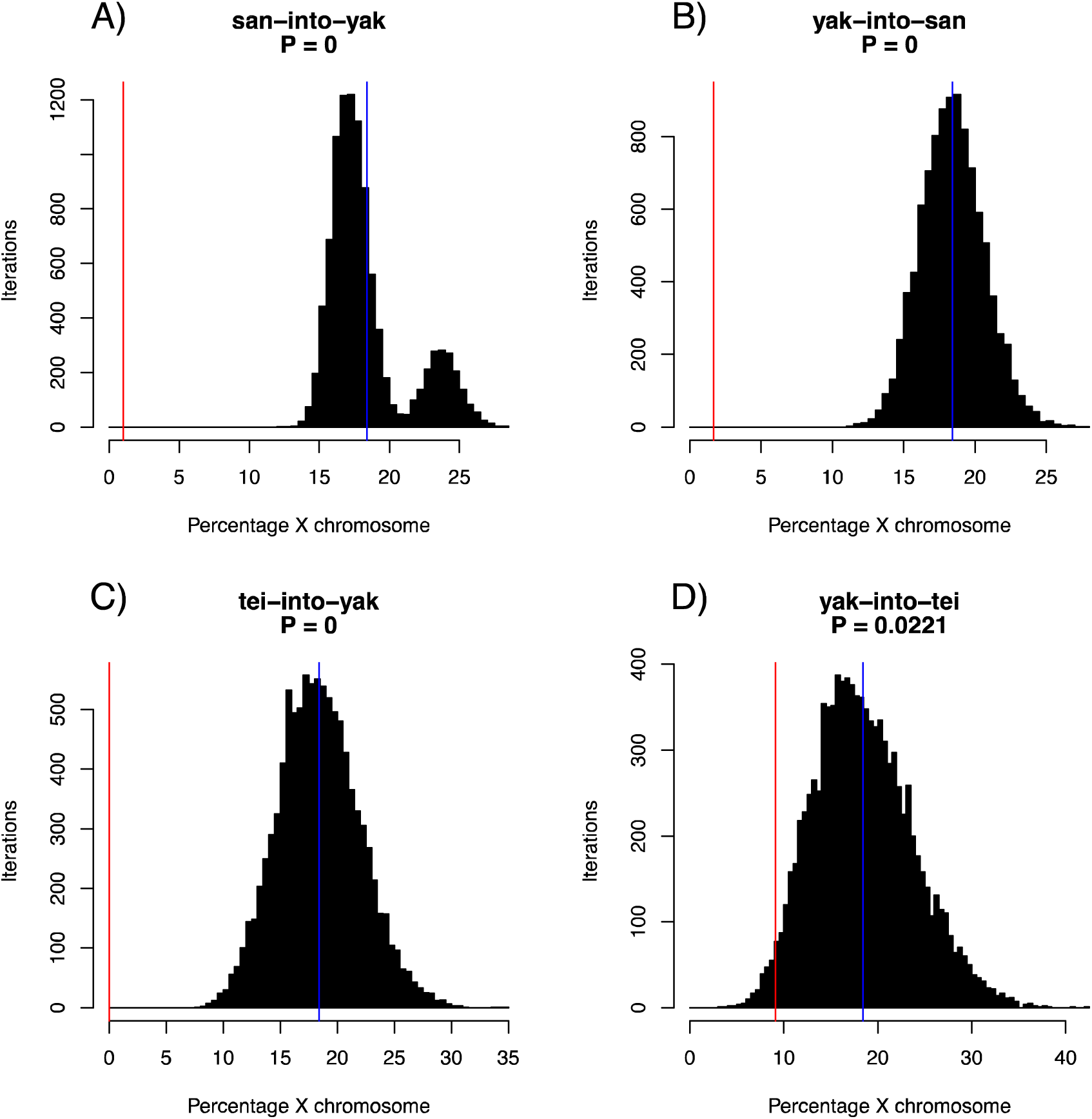
Percentage of the genome containing introgressed tracts following a single generation of introgression. Results for five independent SELAM runs with a population size of 10,000 following a single generation of admixture. The horizontal red line represents the observed value for introgression between *D. yakuba* and *D. santomea* (0.35%), and the horizontal blue line denotes the observed value for introgression between *D. yakuba* and *D. teissieri* (0.01%). **A)** m=0.0001. **B)** m=0.001. **C)** m=0.01. **D)** m=0.1.

**Figure S8.**
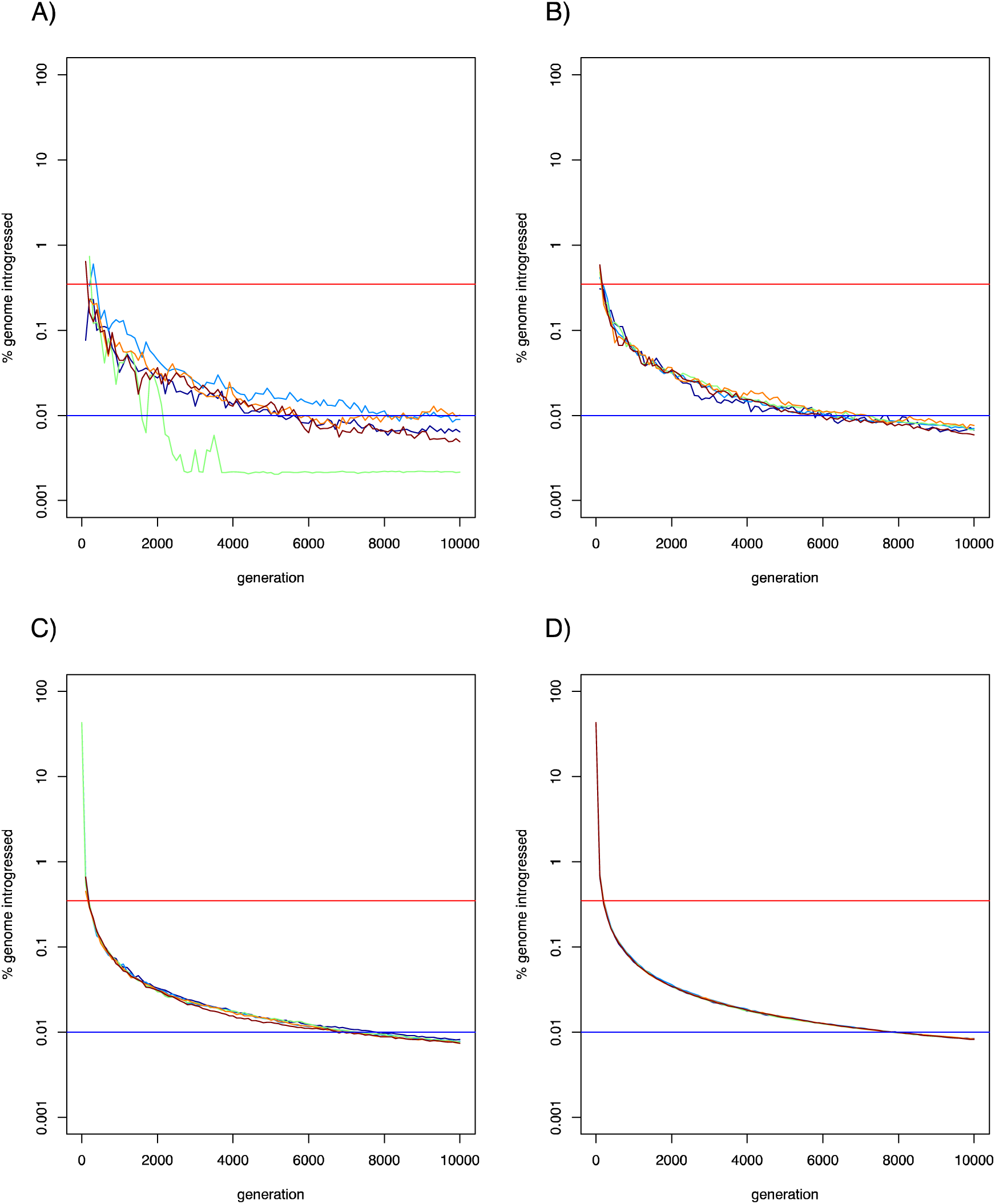
Expected average length of introgression tracts following a single generation of introgression. Results for five independent SELAM runs with a population size of 10,000 following a single generation of admixture. The horizontal red line represents the observed value for introgression between *D. yakuba* and *D. santomea* (6.8kb), and the horizontal blue line denotes the observed value for introgression between *D. yakuba* and *D. teissieri* (2.4kb). **A)** m=0.0001. **B)** m=0.001. **C)** m=0.01. **D)** m=0.1.

**Figure S9.**
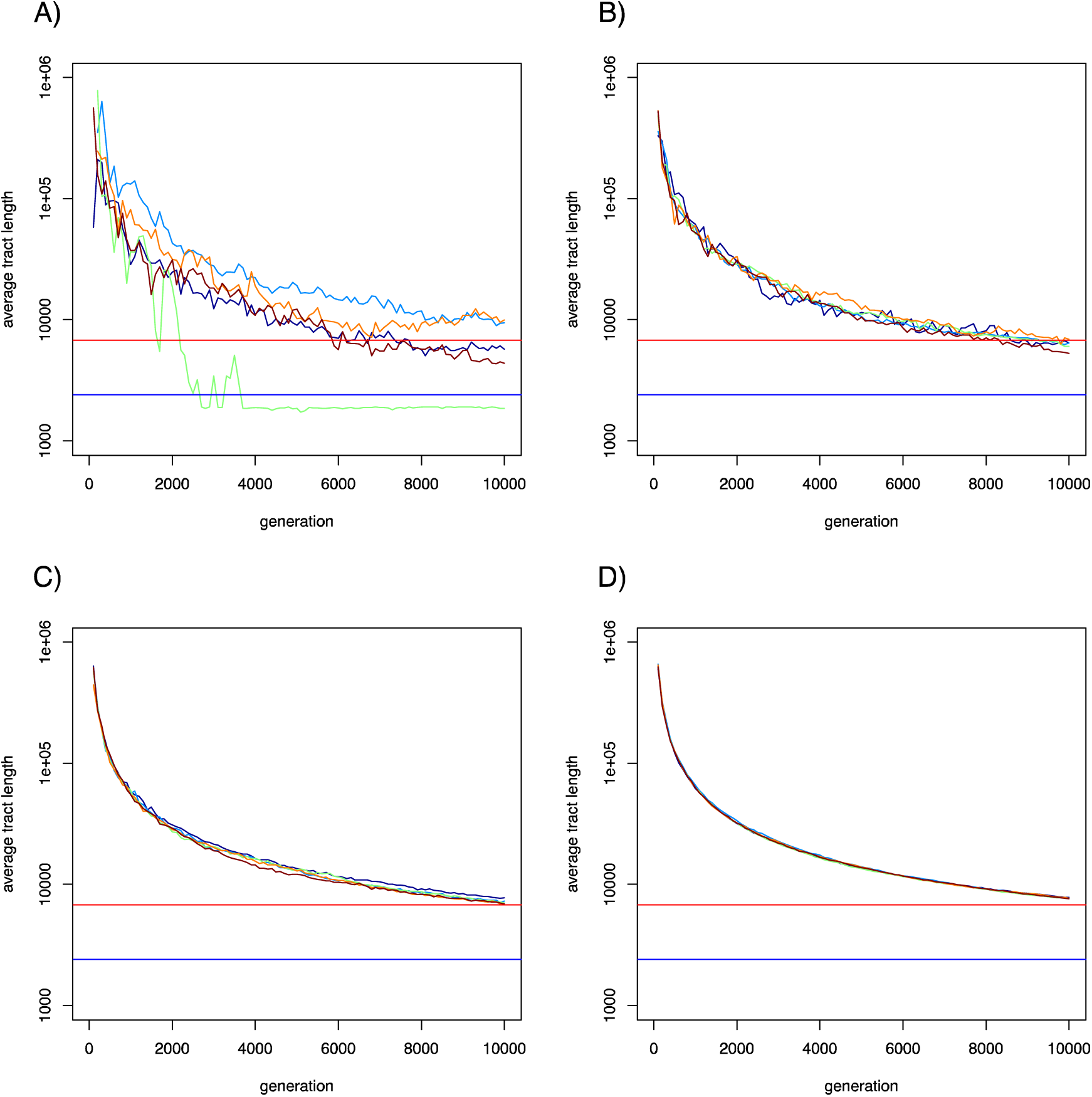
Identified introgressions between *D. yakuba* and *D. santomea* are unlikely to represent ancestral variation that is still segregating in the recipient species. Panels A) through F) show the expected lengths of ancestral haplotype fragments that would still be segregating in the recipient species. The red line and text indicate the 99^th^ quantile of the distribution of fragment sizes. Distributions were calculated assuming different values for the effective population size and generation length. Panels G) and H) show the expected number of generations that an allele at a given frequency p would take to either be fixed (black line) or lost (red line) from the population. Horizontal lines denote the number of populations since the two species diverged assuming generation lengths of 14 days (green line), 21 days (blue line), and 28 days (purple line). **A)** N_e_ = 10^4^ and generation length = 14 days. **B)** N_e_ = 10^4^ and generation length = 21 days. **C)** N_e_ = 10^4^ and generation length = 28 days. **D)** N_e_ = 10^6^ and generation length = 14 days. **E)** N_e_ = 10^6^ and generation length = 21 days. **F)** N_e_ = 10^6^ and generation length = 28 days. **G)** N_e_ = 10^4^. **H)** N_e_ = 10^6^.

**Figure S10.**
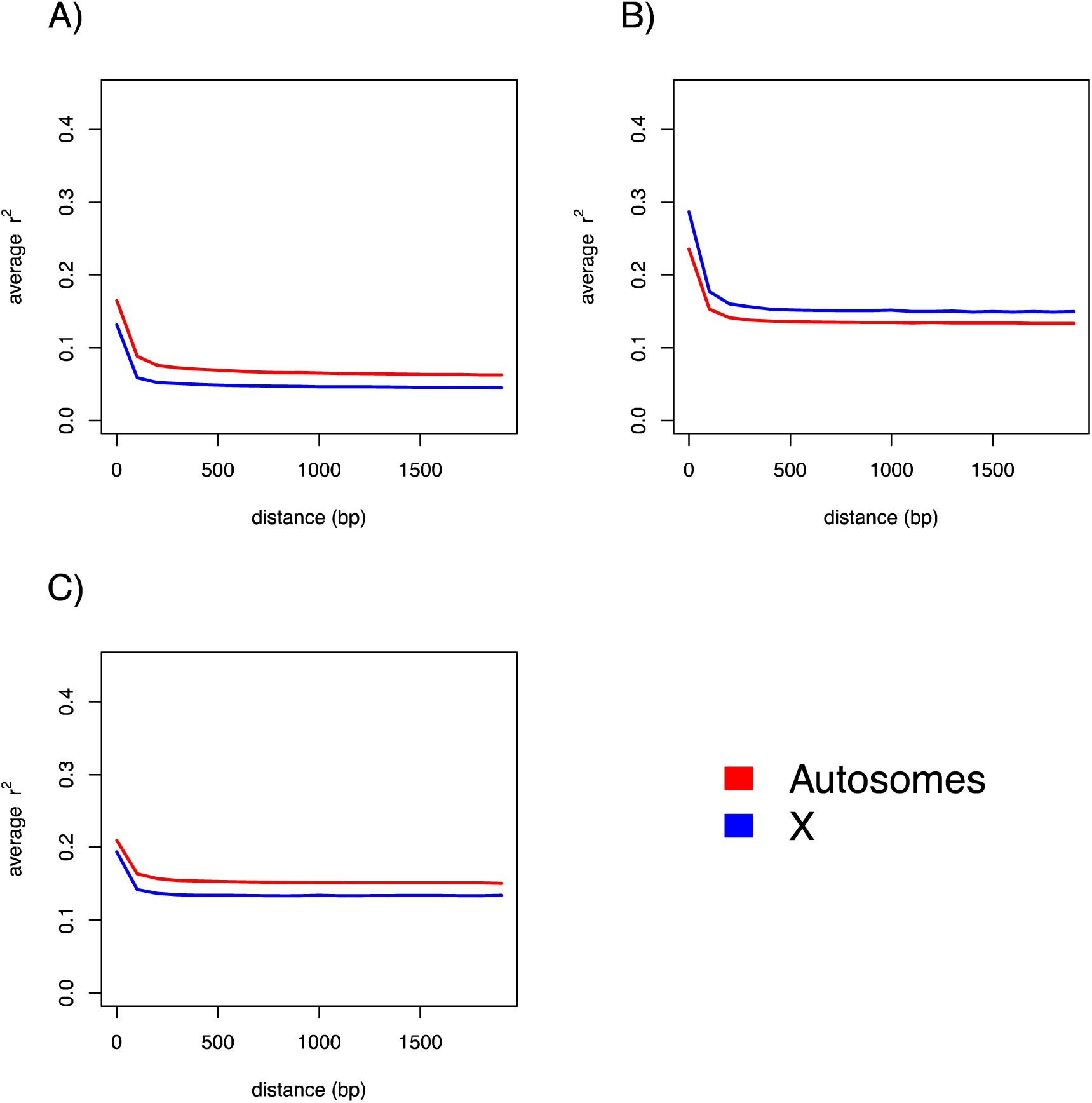
Identified introgressions between *D. yakuba* and *D. teissieri* are unlikely to represent ancestral variation that is still segregating in the recipient species. Panels A) through F) show the expected lengths of ancestral haplotype fragments that would still be segregating in the recipient species. The red line and text indicate the 99^th^ quantile of the distribution of fragment sizes. Distributions were calculated assuming different values for the effective population size and generation length. Panels G) and H) show the expected number of generations that an allele at a given frequency p would take to either be fixed (black line) or lost (red line) from the population. Horizontal lines denote the number of populations since the two species diverged assuming generation lengths of 14 days (green line), 21 days (blue line), and 28 days (purple line). **A)** N_e_ = 10^4^ and generation length = 14 days. **B)** N_e_ = 10^4^ and generation length = 21 days. **C)** N_e_ = 10^4^ and generation length = 28 days. **D)** N_e_ = 10^6^ and generation length = 14 days. **E)** N_e_ = 10^6^ and generation length = 21 days. **F)** N_e_ = 10^6^ and generation length = 28 days. **G)** N_e_ = 10^4^. **H)** N_e_ = 10^6^.

**Figure S11.**
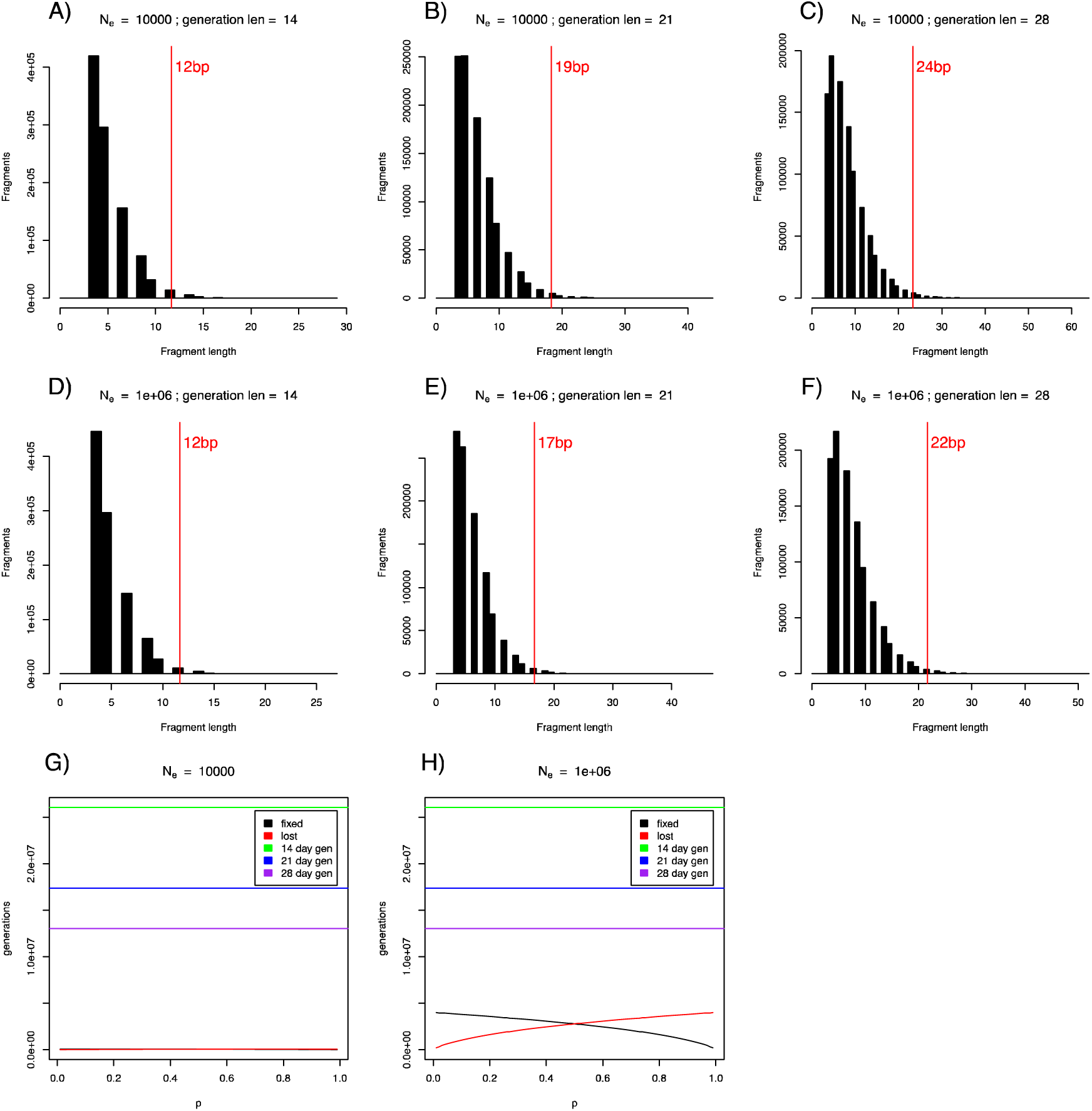
Region on chromosome 2R (2R: 19,918,908-19,927,758) with the *san-* into*-yak* introgressed frequency ≥ 50% in the hybrid zone on São Tomé. Tracts for all 56 *D. yakuba* lines. Red bars indicate homozygous *D. santomea* tracts, purple bars are heterozygous tracts, light bars are homozygous *D. yakuba tracts,* and light pink tracts indicate homozygous donor tracts that were not considered as introgression tracts because they were either less than 500bp, had less than SNPs with the donor allele, or contained more than 30% repetitive sequence. The region of interest is highlighted by a grey rectangle, and lines from the hybrid zone have blue names. The bottom of the plot contains rectangles indicating annotated genes with an arrow indicating the direction of transcription and solid black rectangles denoting coding sequence.

**Figure S12.**
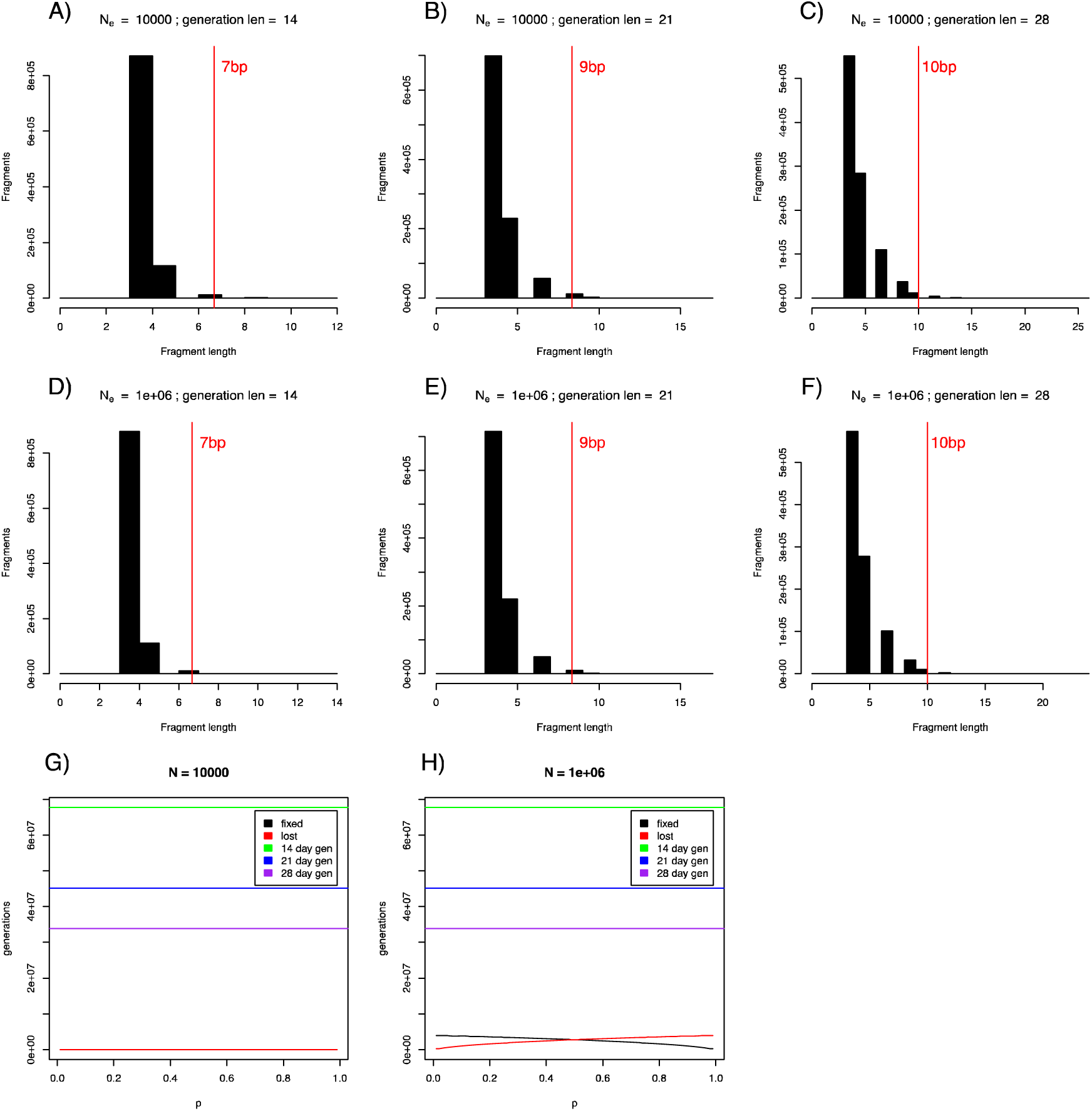
Region on chromosome 3L (3L: 6,225,896-6,257,088) with the *san*-into-yak introgressed frequency ≥ 50% in the hybrid zone on São Tomé. Tracts for all 56 *D. yakuba* lines. Red bars indicate homozygous *D. santomea* tracts, purple bars are heterozygous tracts, light bars are homozygous *D. yakuba* tracts, and light pink tracts indicate homozygous donor tracts that were not considered as introgression tracts because they were either less than 500bp, had less than SNPs with the donor allele, or contained more than 30% repetitive sequence. The region of interest is highlighted by a grey rectangle, and lines from the hybrid zone have blue names. The bottom of the plot contains rectangles indicating annotated genes with an arrow indicating the direction of transcription and solid black rectangles denoting coding sequence.

**Figure S13.**
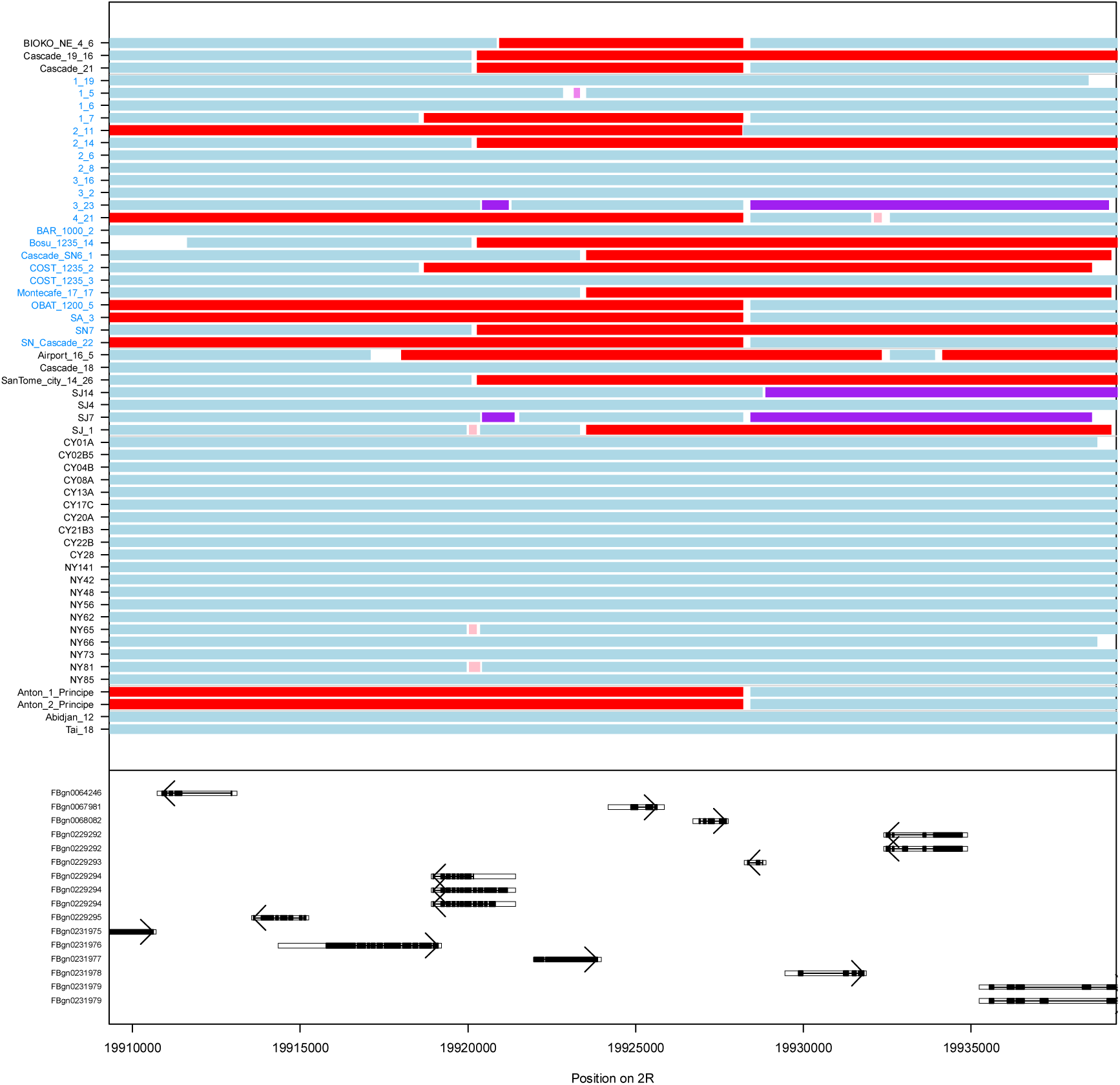
Region on chromosome 3L (3L: 12,187,525-12,209,675) with the *san-* into*-yak* introgressed frequency ≥ 50% in the hybrid zone on São Tomé. Tracts for all 56 *D. yakuba* lines. Red bars indicate homozygous *D. santomea* tracts, purple bars are heterozygous tracts, light bars are homozygous *D. yakuba tracts,* and light pink tracts indicate homozygous donor tracts that were not considered as introgression tracts because they were either less than 500bp, had less than SNPs with the donor allele, or contained more than 30% repetitive sequence. The region of interest is highlighted by a grey rectangle, and lines from the hybrid zone have blue names. The bottom of the plot contains rectangles indicating annotated genes with an arrow indicating the direction of transcription and solid black rectangles denoting coding sequence.

**Figure S14.**
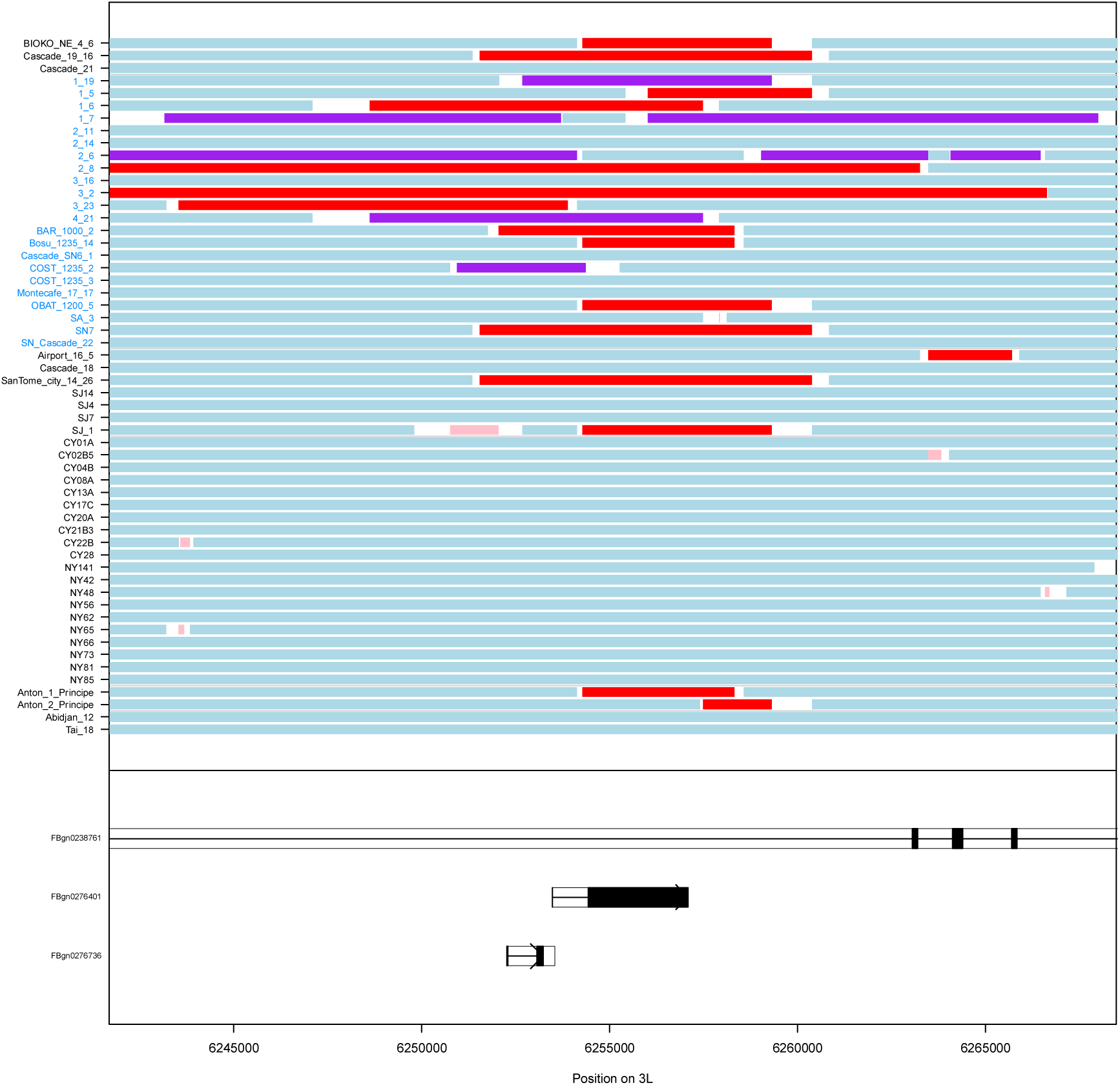
Collection map. Map indicating the number of fly lines used in this study that were collected from each geographic location. ‘ST:HZ’ is the hybrid zone on Säo Tomé, and ‘ST: Low’ are the lowland areas on the island of São Tomé.

**Figure S15.**
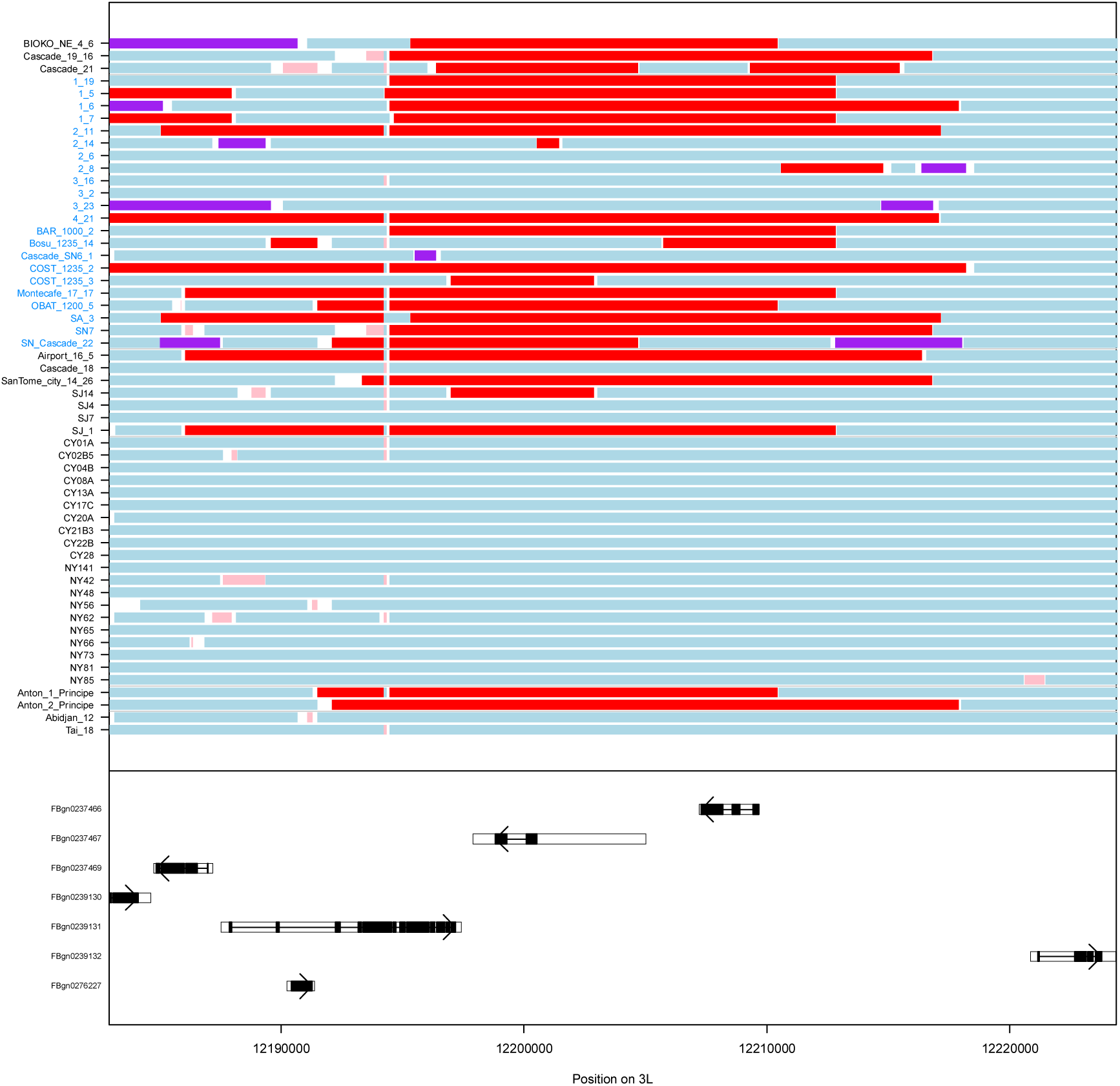
Example introgressions. Example introgressions from *D. santomea* into *D. yakuba* for a region on chromosome arm 2R. **A)** Introgression from *D. santomea* into the *D. yakuba* line SanTome_city_14_26. ‘SNPs’ represent the markers for this line. ‘Coverages’ show the number of reads with either the donor (*D. santomea,* red) or recipient allele (*D. yakuba*; light blue) at each site. Coverages greater than 25x were downscaled to integer values between 0 and 25x. ‘Probabilities’ are the probabilities returned by Int-HMM for all six states at each site (light blue: homozygous recipient, purple: heterozygous, red: homozygous donor, light grey: homozygous recipient error state, black heterozygous error state, and dark grey homozygous donor error state). ‘Unfiltered’ represent the raw tracks obtained by grouping contiguous blocks of SNPs with the same most probable state. ‘Tracts’ are the filtered tracts. **B)** The same region from A) but showing the filtered tracts for all 56 *D. yakuba* lines. Light pink tracts indicate homozygous donor tracts that were not considered as introgression tracts because they were either less than 500bp, had less than SNPs with the donor allele, or contained more than 30% repetitive sequence. The bottom of the plot contains rectangles indicating annotated genes with an arrow indicating the direction of transcription and solid black rectangles denoting coding sequence.

## SUPPLEMENTARY TABLES

**Table SI.**
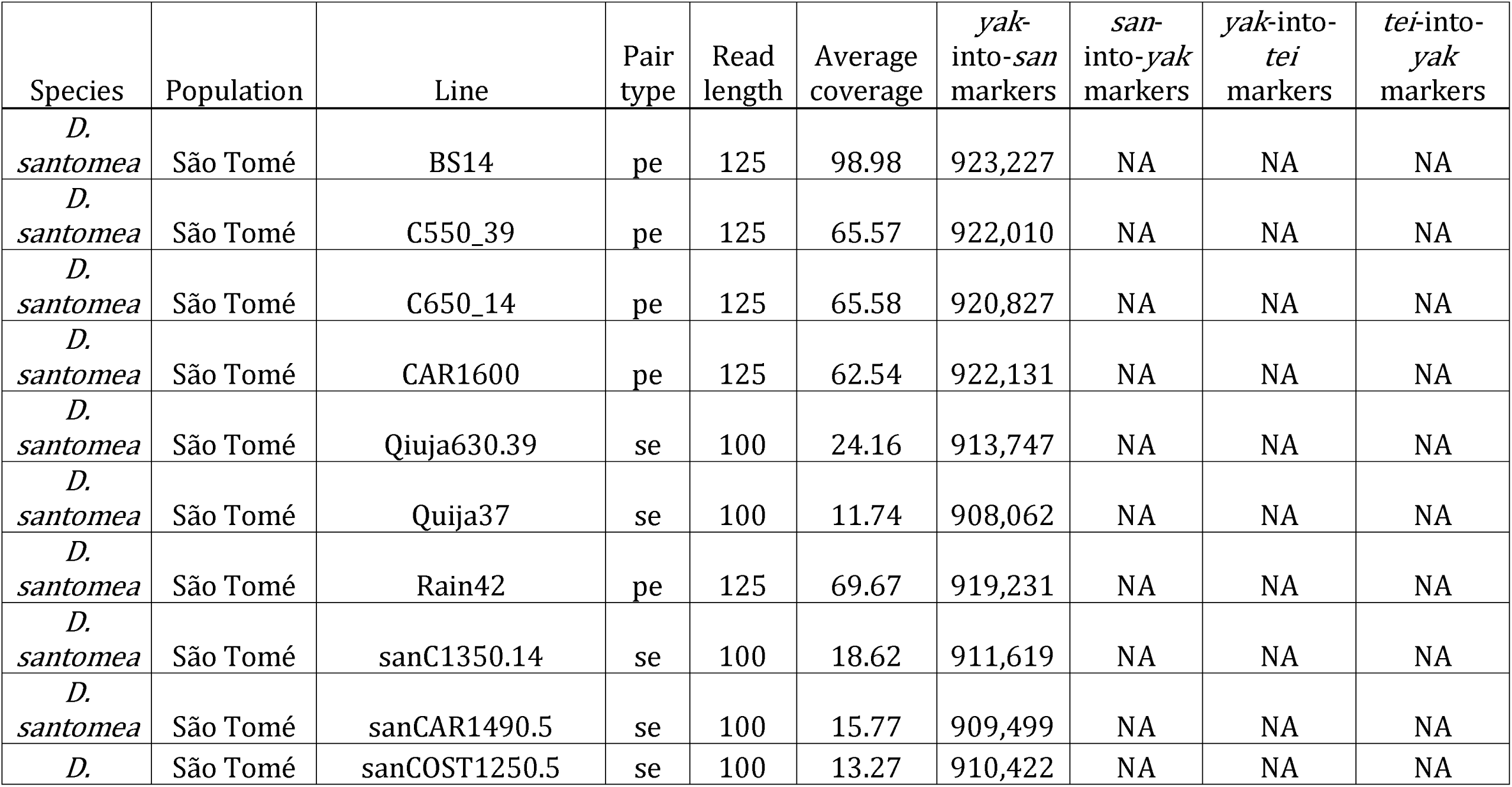
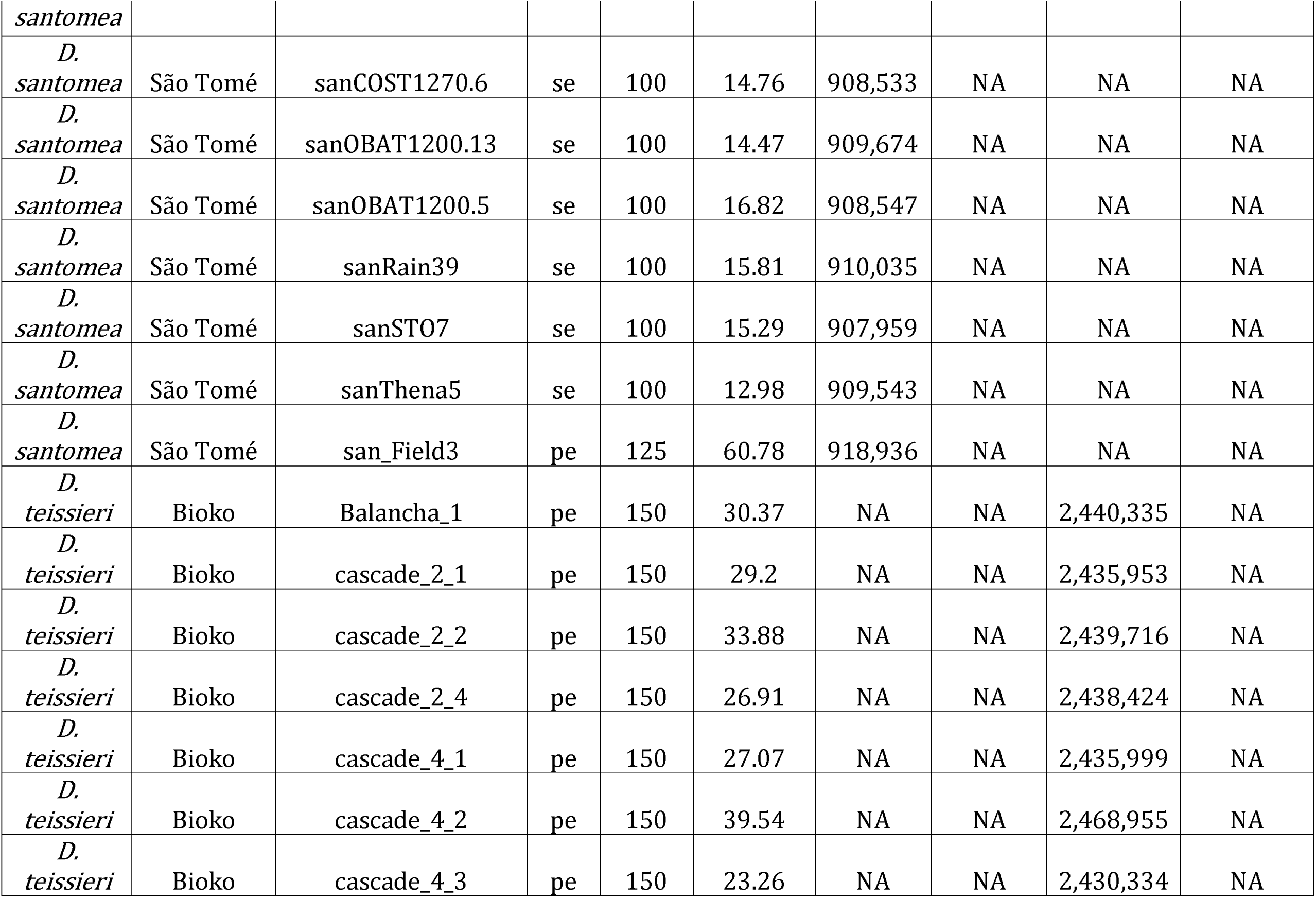
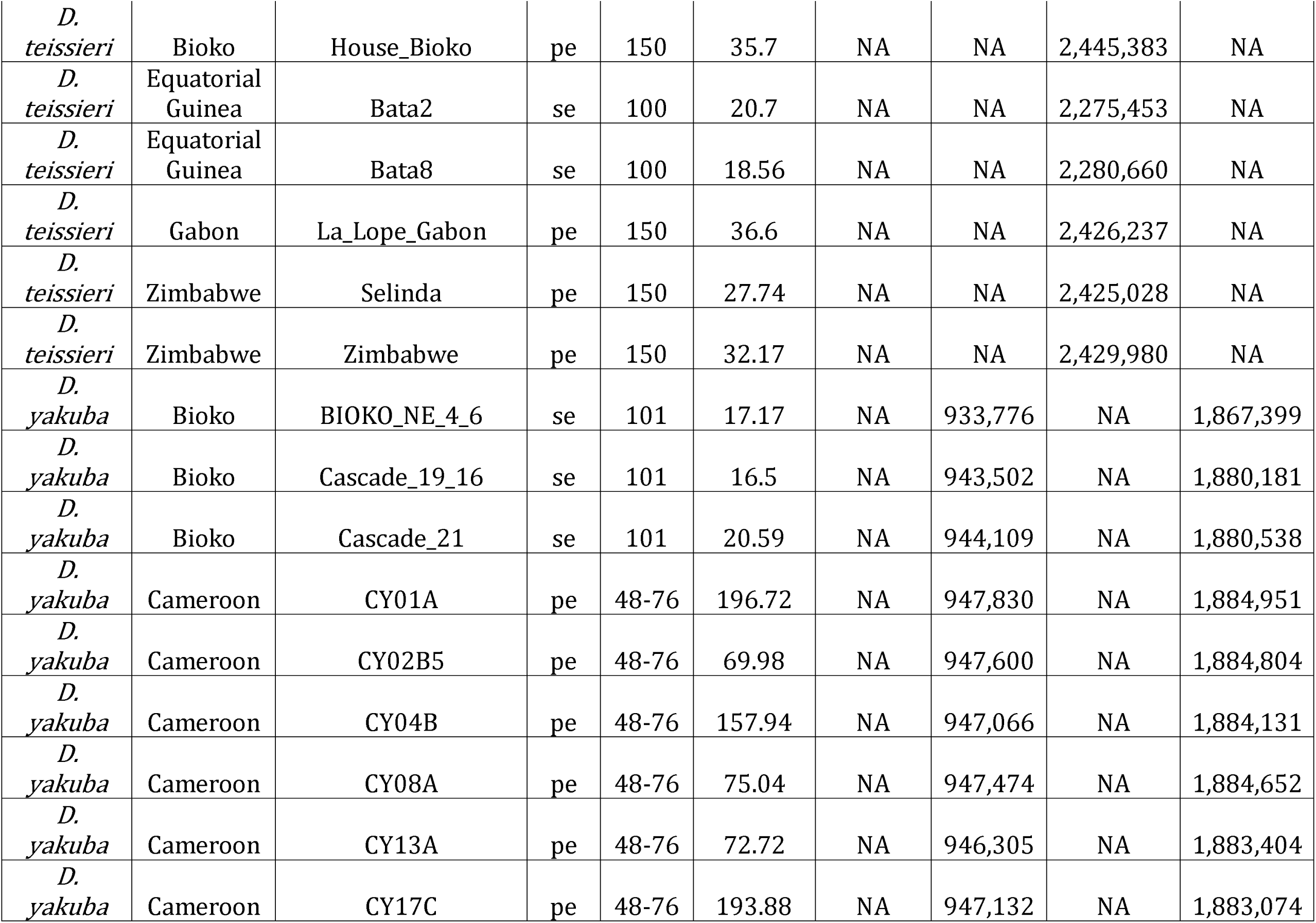
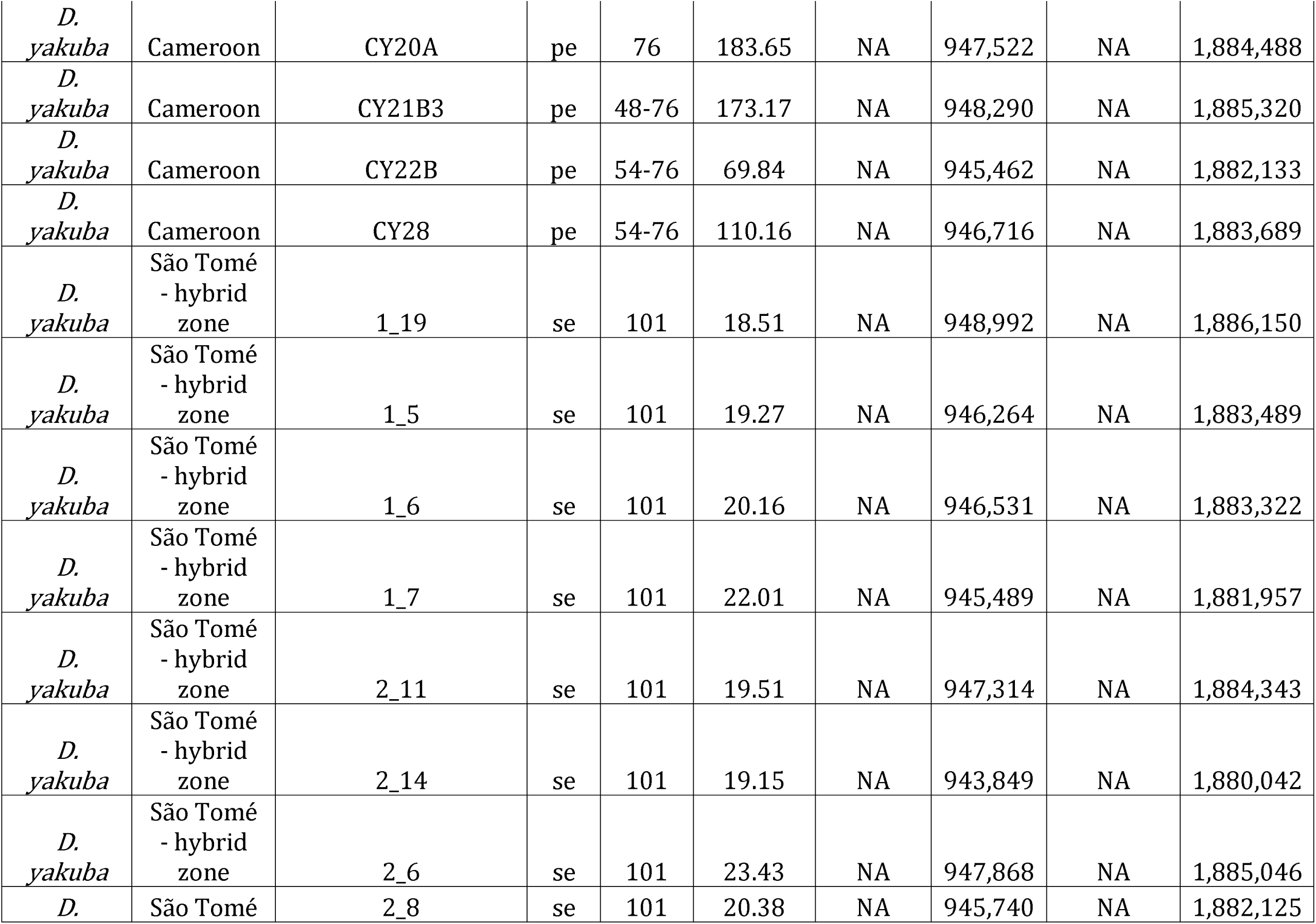
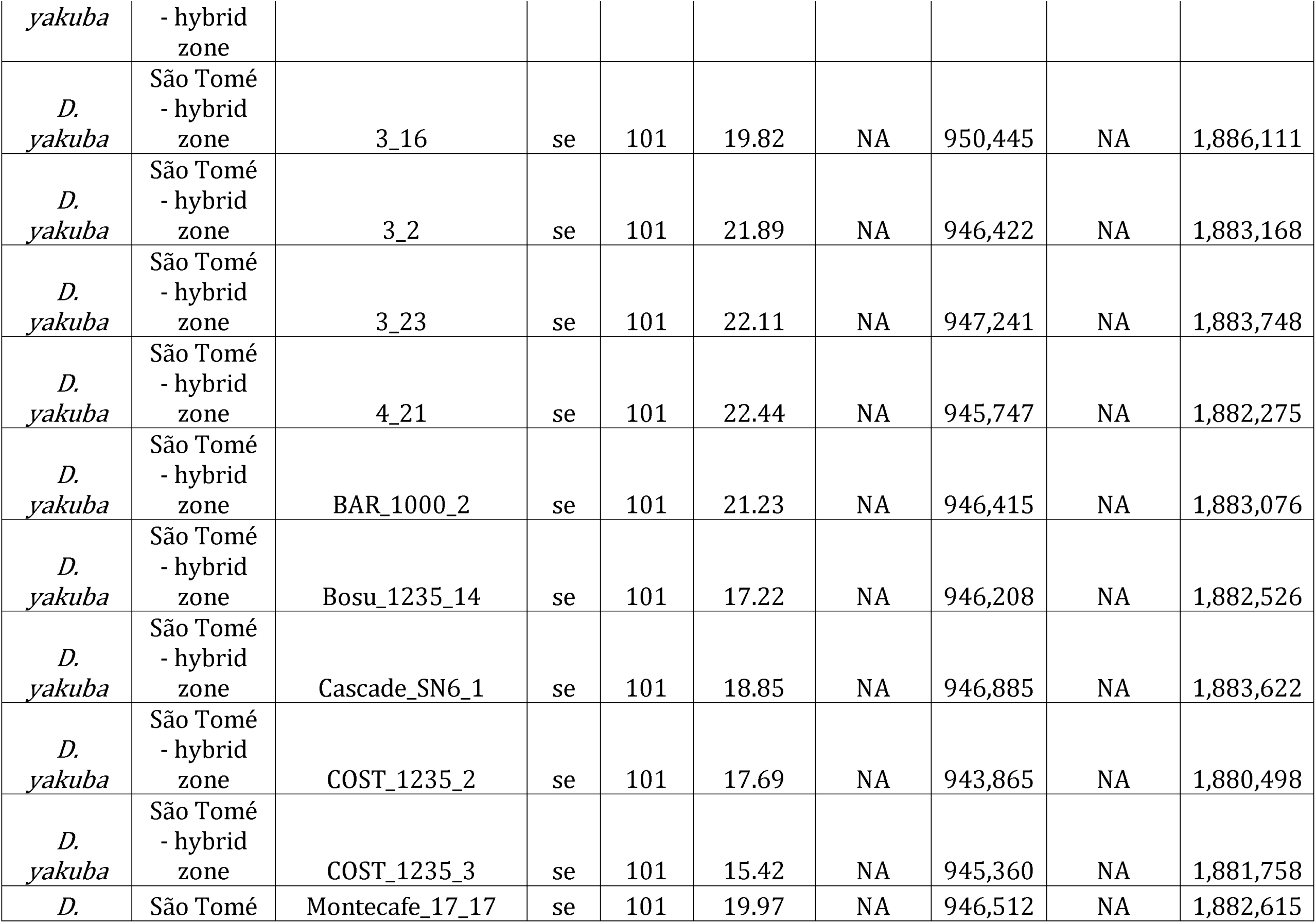
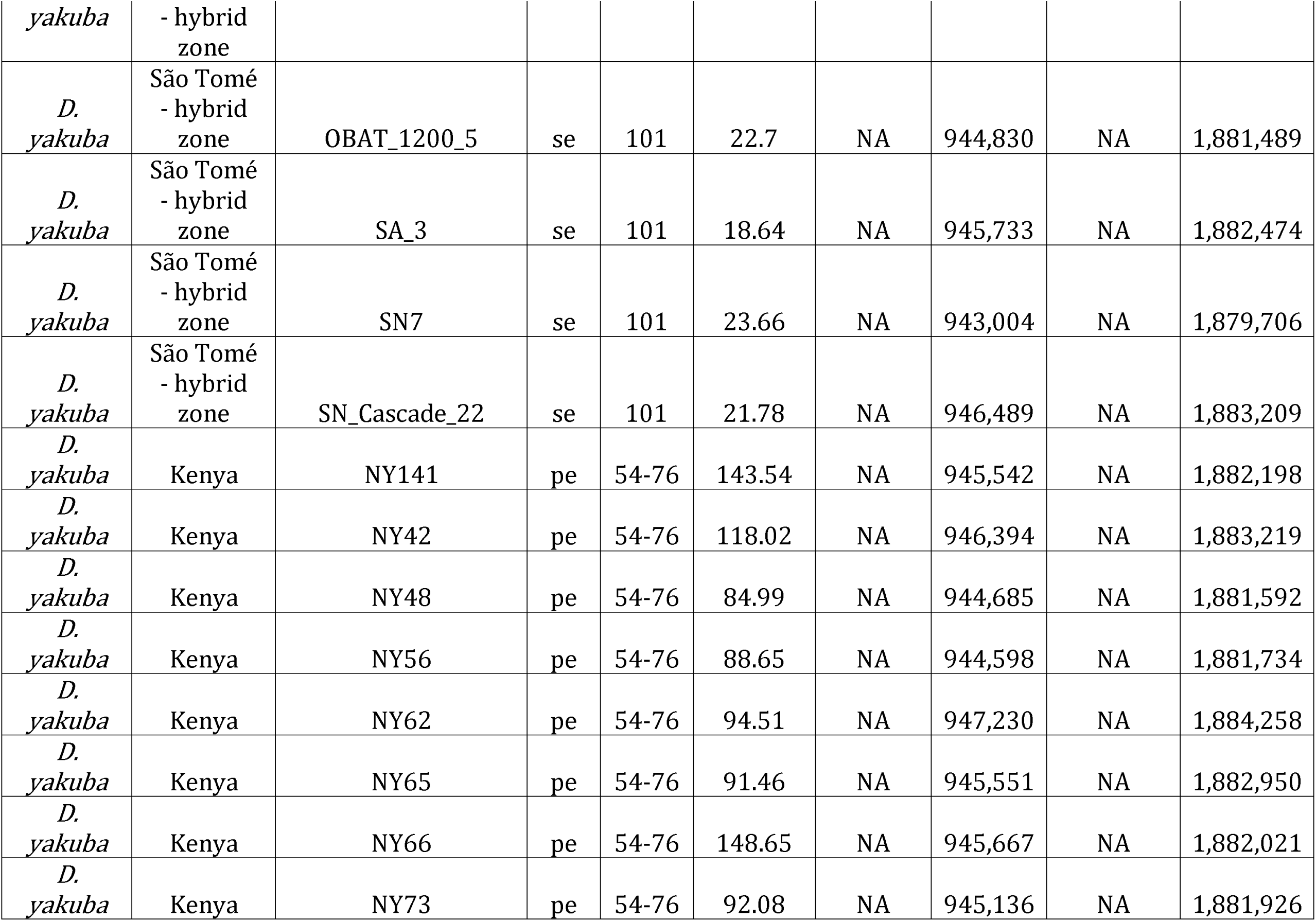
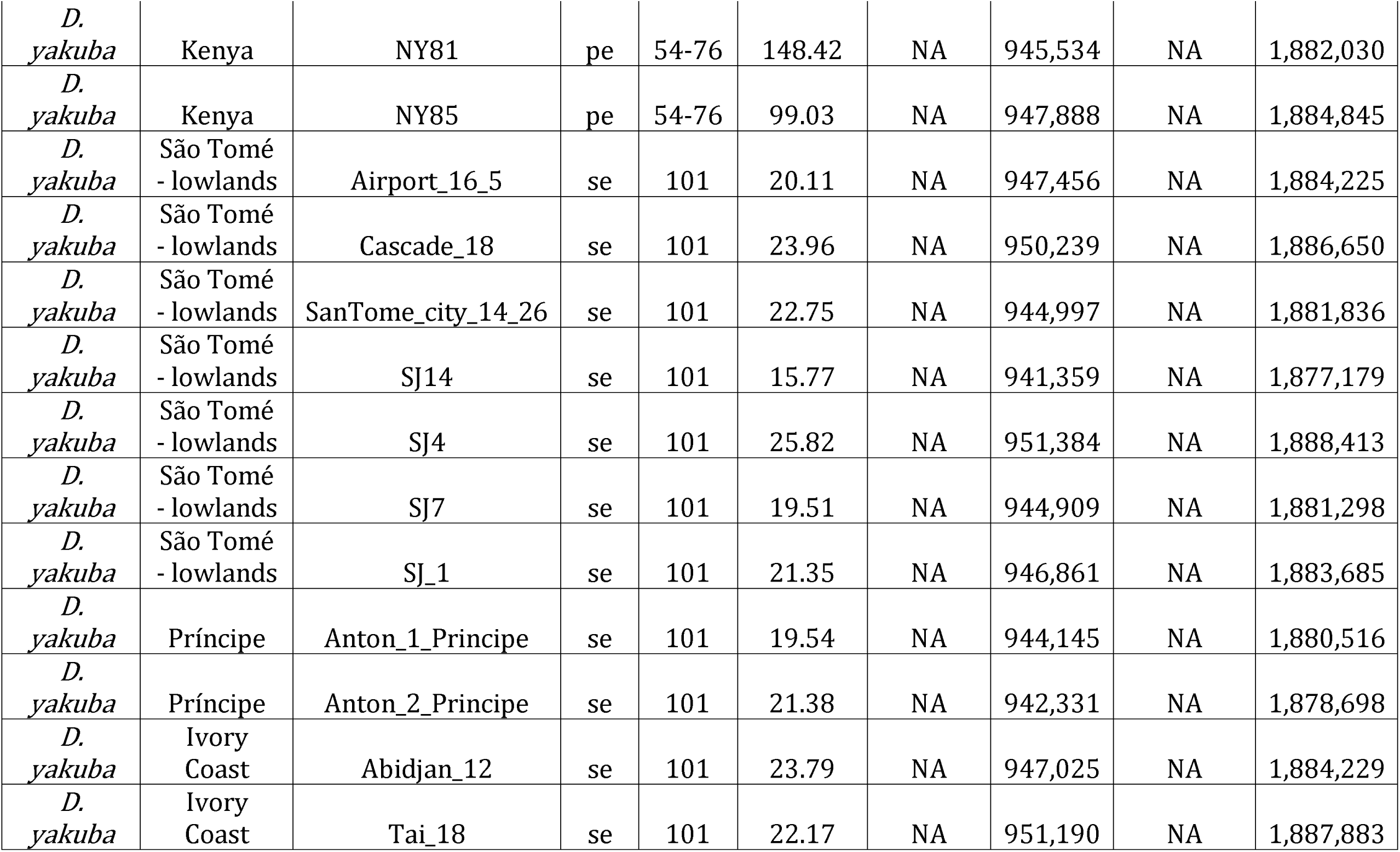
Fly lines used in this study. Lines used in the study, their geographic origin, and the length and paired status (se = single end, pe = paired end) of Illumina sequencing reads. Average coverage is the average number of reads mapped overlapping a given site in the genome. The markers columns denote the number of markers used in the HMM when identifying a given direction of introgression.

**Table S2.** False positive tracts from identified by Int-HMM from the simulated data. Counts denote the number of tracts, percentages refer to amount of genomic sequence covered by those tracts, and cum lengths are the combined tract lengths.

| direction | homozygous<br>count | homozygous<br>percentage | homozygous<br>cum length | heterozygous<br>count | heterozygous<br>percentage | heterozygous<br>cumulative<br>length |
| --- | --- | --- | --- | --- | --- | --- |
| *yak*-into-<br>*san* | 2 | 0.177 | 2087bp | 0 | 0 | 0bp |
| *san*-into-<br>*yak* | 3 | 0.253 | 5652bp | 0 | 0 | 0bp |
| *yak*-into-<br>*tei* | 1 | 0.0702 | 800bp | 0 | 0 | 0bp |
| *tei*-into-<br>*yak* | 0 | 0 | 0bp | 0 | 0 | 0bp |

**Table S3.** Percentage of genome that was introgressed. Percentage of the genome that was introgressed for each line as determined by the cumulative length of introgression tracts identified by the HMM.

| Species | Population | Line | *yak*-into-<br>*san* | *san*-into-<br>*yak* | *yak*-into-<br>*tei* | *tei*-into-<br>*yak* |
| --- | --- | --- | --- | --- | --- | --- |
| *D. santomea* | São Tomé | Quija630.39 | 0.1006 | NA | NA | NA |
| *D. santomea* | São Tomé | Quija37 | 0.4365 | NA | NA | NA |
| *D. santomea* | São Tomé | sanC1350.14 | 0.1221 | NA | NA | NA |
| *D. santomea* | São Tomé | sanCAR1490.5 | 0.5787 | NA | NA | NA |
| *D. santomea* | São Tomé | sanCOST1250.5 | 0.1037 | NA | NA | NA |
| *D. santomea* | São Tomé | sanCOST1270.6 | 0.1698 | NA | NA | NA |
| *D. santomea* | São Tomé | sanOBAT1200.13 | 0.3148 | NA | NA | NA |
| *D. santomea* | São Tomé | sanOBAT1200.5 | 0.1028 | NA | NA | NA |
| *D. santomea* | São Tomé | sanRain39 | 0.1501 | NA | NA | NA |
| *D. santomea* | São Tomé | sanST07 | 0.5866 | NA | NA | NA |
| *D. santomea* | São Tomé | sanThena5 | 0.34 | NA | NA | NA |
| *D. santomea* | São Tomé | Rain42 | 0.5293 | NA | NA | NA |
| *D.* | São Tomé | BS14 | 0.4534 | NA | NA | NA |
| *santomea* |  |  |  |  |  |  |
| *D. santomea* | São Tomé | C650_14 | 0.1293 | NA | NA | NA |
| *D. santomea* | São Tomé | C550_39 | 0.5837 | NA | NA | NA |
| *D. santomea* | São Tomé | san_Field3 | 1.0401 | NA | NA | NA |
| *D. santomea* | São Tomé | CAR1600 | 0.212 | NA | NA | NA |
| *D. teissieri* | Bioko | Balancha_1 | NA | NA | 0.0053 | NA |
| *D. teissieri* | Bioko | cascade_4_3 | NA | NA | 0.0112 | NA |
| *D. teissieri* | Bioko | House_Bioko | NA | NA | 0.0123 | NA |
| *D. teissieri* | Bioko | cascade_4_2 | NA | NA | 0.0107 | NA |
| *D. teissieri* | Bioko | cascade_4_1 | NA | NA | 0.0053 | NA |
| *D. teissieri* | Bioko | cascade_2_4 | NA | NA | 0.0093 | NA |
| *D. teissieri* | Bioko | cascade_2_2 | NA | NA | 0.0052 | NA |
| *D. teissieri* | Bioko | cascade_2_1 | NA | NA | 0 | NA |
| *D. teissieri* | Equatorial Guinea | Bata8 | NA | NA | 0.0008 | NA |
| *D. teissieri* | Equatorial Guinea | Bata2 | NA | NA | 0.004 | NA |
| *D. teissieri* | Gabon | La_Lope_Gabon | NA | NA | 0.0065 | NA |
| *D. teissieri* | Zimbabwe | Zimbabwe | NA | NA | 0.006 | NA |
| *D. teissieri* | Zimbabwe | Selinda | NA | NA | 0.0129 | NA |
| *D. yakuba* | Bioko | BIOKO_NE_4_6 | NA | 0.2751 | NA | 0.0088 |
| *D. yakuba* | Bioko | Cascade_19_16 | NA | 0.3349 | NA | 0.0144 |
| *D. yakuba* | Bioko | Cascade_21 | NA | 0.321 | NA | 0.0011 |
| *D. yakuba* | Cameroon | CY28 | NA | 0.1097 | NA | 0.0089 |
| *D. yakuba* | Cameroon | CY22B | NA | 0.0688 | NA | 0.0007 |
| *D. yakuba* | Cameroon | CY21B3 | NA | 0.0146 | NA | 0.0054 |
| *D. yakuba* | Cameroon | CY20A | NA | 0.0919 | NA | 0.0215 |
| *D. yakuba* | Cameroon | CY17C | NA | 0.0168 | NA | 0.001 |
| *D. yakuba* | Cameroon | CY13A | NA | 0.0922 | NA | 0.0073 |
| *D. yakuba* | Cameroon | CY08A | NA | 0.031 | NA | 0.0137 |
| *D. yakuba* | Cameroon | CY04B | NA | 0.0492 | NA | 0.0144 |
| *D. yakuba* | Cameroon | CY01A | NA | 0.0483 | NA | 0.019 |
| *D. yakuba* | Cameroon | CY02B5 | NA | 0.0342 | NA | 0.0084 |
| *D. yakuba* | São Tomé - hybrid zone | Cascade_SN6_1 | NA | 0.3078 | NA | 0.0032 |
| *D. yakuba* | São Tomé - hybrid zone | SN7 | NA | 0.3645 | NA | 0.0144 |
| *D. yakuba* | São Tomé - hybrid zone | SN_Cascade_22 | NA | 0.3039 | NA | 0.0175 |
| *D. yakuba* | São Tomé - hybrid zone | 1_19 | NA | 0.3896 | NA | 0.0065 |
| *D. yakuba* | São Tomé - hybrid zone | 1_5 | NA | 0.3047 | NA | 0.0138 |
| *D. yakuba* | São Tomé - hybrid zone | 1_6 | NA | 0.339 | NA | 0.0129 |
| *D. yakuba* | São Tomé - hybrid zone | 1_7 | NA | 0.3717 | NA | 0.0124 |
| *D. yakuba* | São Tomé - hybrid zone | 2_11 | NA | 0.2351 | NA | 0.0107 |
| *D. yakuba* | São Tomé - hybrid zone | 2_14 | NA | 0.2357 | NA | 0.0007 |
| *D. yakuba* | São Tomé - hybrid zone | 2_6 | NA | 0.3771 | NA | 0.006 |
| *D. yakuba* | São Tomé - hybrid zone | 2_8 | NA | 0.357 | NA | 0.0244 |
| *D. yakuba* | São Tomé - hybrid zone | 3_16 | NA | 0.012 | NA | 0.0176 |
| *D. yakuba* | São Tomé - hybrid zone | 3_2 | NA | 0.4387 | NA | 0.0101 |
| *D. yakuba* | São Tomé - hybrid zone | 3_23 | NA | 0.4047 | NA | 0.0073 |
| *D. yakuba* | São Tomé - hybrid zone | 4_21 | NA | 0.3901 | NA | 0.0153 |
| *D. yakuba* | São Tomé - hybrid zone | COST_1235_2 | NA | 0.304 | NA | 0.0014 |
| *D. yakuba* | São Tomé - hybrid zone | COST_1235_3 | NA | 0.1677 | NA | 0.004 |
| *D. yakuba* | São Tomé - hybrid zone | Montecafe_17_17 | NA | 0.3496 | NA | 0 |
| *D. yakuba* | São Tomé - hybrid zone | BAR_1000_2 | NA | 0.3118 | NA | 0.0072 |
| *D. yakuba* | São Tomé - hybrid zone | Bosu_1235_14 | NA | 0.3575 | NA | 0.0008 |
| *D. yakuba* | São Tomé - hybrid zone | OBAT_1200_5 | NA | 0.3394 | NA | 0.0072 |
| *D. yakuba* | São Tomé - hybrid zone | SA_3 | NA | 0.2419 | NA | 0.0053 |
| *D. yakuba* | Kenya | NY81 | NA | 0.1487 | NA | 0.0084 |
| *D. yakuba* | Kenya | NY85 | NA | 0.0211 | NA | 0.0008 |
| *D. yakuba* | Kenya | NY66 | NA | 0.1322 | NA | 0.0049 |
| *D. yakuba* | Kenya | NY65 | NA | 0.0813 | NA | 0.0036 |
| *D. yakuba* | Kenya | NY62 | NA | 0.0667 | NA | 0.0069 |
| *D. yakuba* | Kenya | NY42 | NA | 0.0821 | NA | 0.0097 |
| *D. yakuba* | Kenya | NY48 | NA | 0.0608 | NA | 0.0092 |
| *D. yakuba* | Kenya | NY73 | NA | 0.0581 | NA | 0.0074 |
| *D. yakuba* | Kenya | NY56 | NA | 0.0739 | NA | 0.001 |
| *D. yakuba* | Kenya | NY141 | NA | 0.0301 | NA | 0.0084 |
| *D. yakuba* | São Tomé - lowlands | Airport_16_5 | NA | 0.2525 | NA | 0.0021 |
| *D. yakuba* | São Tomé - lowlands | Cascade_18 | NA | 0.013 | NA | 0.0114 |
| *D. yakuba* | São Tomé - lowlands | SanTome_city_14_26 | NA | 0.3421 | NA | 0.0133 |
| *D. yakuba* | São Tomé - lowlands | SJ14 | NA | 0.171 | NA | 0.0044 |
| *D. yakuba* | São Tomé - lowlands | SJ4 | NA | 0.0264 | NA | 0.0087 |
| *D. yakuba* | São Tomé - lowlands | SJ7 | NA | 0.2344 | NA | 0.0017 |
| *D. yakuba* | São Tomé - lowlands | SJ_1 | NA | 0.3074 | NA | 0 |
| *D. yakuba* | Príncipe | Anton_1_Principe | NA | 0.3142 | NA | 0.0072 |
| *D. yakuba* | Príncipe | Anton_2_Principe | NA | 1.2012 | NA | 0 |
| *D. yakuba* | Ivory Coast | Tai_18 | NA | 0.0182 | NA | 0.0103 |
| *D. yakuba* | Ivory Coast | Abidjan_12 | NA | 0.1756 | NA | 0.0143 |

**Table S4.** Distribution of markers used by Int-HMM is similar across sequence types in all the four introgressions directions.

| Direction | type | markers | total | Markers<br>per kb |
| --- | --- | --- | --- | --- |
| *yak*-into-*san* | 10kb inter | 166,112 | 22,780,104 | 7.292 |
| *yak*-into-*san* | 2kb upstream inter | 92,911 | 10,100,186 | 9.1989 |
| *yak*-into-*san* | 3'prime UTR | 31,596 | 4,121,972 | 7.6653 |
| *yak*-into-*san* | 5'prime UTR | 27,077 | 3,346,831 | 8.0903 |
| *yak*-into-*san* | CDS | 145,211 | 22,059,257 | 6.5828 |
| *yak*-into-*san* | exon | 2,375 | 317,599 | 7.478 |
| *yak*-into-*san* | intergenic | 50,263 | 8,754,040 | 5.7417 |
| *yak*-into-*san* | intron | 415,743 | 48,178,319 | 8.6293 |
| *san*-into-*yak* | 10kb inter | 166,866 | 22,780,104 | 7.3251 |
| *san*-into-*yak* | 2kb upstream inter | 98,371 | 10,100,186 | 9.7395 |
| *san*-into-*yak* | 3'prime UTR | 30,881 | 4,121,972 | 7.4918 |
| *san*-into-*yak* | 5'prime UTR | 26,267 | 3,346,831 | 7.8483 |
| *san*-into-*yak* | CDS | 156,748 | 22,059,257 | 7.1058 |
| *san*-into-*yak* | exon | 2,207 | 317,599 | 6.949 |
| *san*-into-*yak* | intergenic | 49,851 | 8,754,040 | 5.6946 |
| *san*-into-*yak* | intron | 421,513 | 48,178,319 | 8.749 |
| *yak*-into-*tei* | 10kb inter | 435,107 | 22,780,104 | 19.1003 |
| *yak*-into-*tei* | 2kb upstream inter | 237,045 | 10,100,186 | 23.4694 |
| *yak*-into-*tei* | 3'prime UTR | 83,077 | 4,121,972 | 20.1547 |
| *yak*-into-*tei* | 5'prime UTR | 74,830 | 3,346,831 | 22.3585 |
| *yak*-into-*tei* | CDS | 475,558 | 22,059,257 | 21.5582 |
| *yak*-into-*tei* | exon | 6,035 | 317,599 | 19.0019 |
| *yak*-into-*tei* | intergenic | 131,295 | 8,754,040 | 14.9982 |
| *yak*-into-*tei* | intron | 1,105,338 | 48,178,319 | 22.9426 |
| *tei*-into-*yak* | 10kb inter | 307,221 | 22,780,104 | 13.4864 |
| *tei*-into-*yak* | 2kb upstream inter | 168,122 | 10,100,186 | 16.6454 |
| *tei*-into-*yak* | 3'prime UTR | 67,187 | 4,121,972 | 16.2997 |
| *tei*-into-*yak* | 5'prime UTR | 61,183 | 3,346,831 | 18.2809 |
| *tei*-into-*yak* | CDS | 401,401 | 22,059,257 | 18.1965 |
| *tei*-into-*yak* | exon | 4,589 | 317,599 | 14.449 |
| *tei*-into-*yak* | intergenic | 91,980 | 8,754,040 | 10.5071 |
| *tei*-into-*yak* | intron | 788,765 | 48,178,319 | 16.3718 |

**Table S5.** Sequence types containing introgressions. The genome was partitioned by sequence type with each region being assigned to a single sequence type with the following hierarchy: CDS (coding sequence), exon, 5prime UTR, 3prime UTR, intron, 2kb upstream inter (intergenic sequence 2kb upstream of a gene), lOkb inter (intergenic sequence within lOkb of a gene), and intergenic (intergenic sequence more than lOkb from a gene). ‘Introgressed percentage’ is the percentage of introgressions overlapping a given sequence type for that direction, ‘Genomic percentage’ is the percentage of the genome represented by a given sequence type, and ‘Enrichment’ = (Introgressed percentage) / (Genomic percentage). P-values were calculated with permutation tests as described in the Methods.

| Direction | Sequence type | Length (kb) | Introgressed percentage | Genomic percentage | Enrichment | P-value |
| --- | --- | --- | --- | --- | --- | --- |
| *yak*-into-*san* | 10kb inter | 553.5 | 23.5 | 19.0 | 1.23 | 0.0020 |
| *yak*-into-*san* | 2kb upstream inter | 130.1 | 5.5 | 8.5 | 0.65 | 0.1980 |
| *yak*-into-*san* | 3prime UTR | 23.4 | 1.0 | 3.5 | 0.29 | 0.0010 |
| *yak*-into-*san* | 5prime UTR | 6.4 | 0.3 | 2.8 | 0.10 | 0.3020 |
| *yak*-into-*san* | CDS | 148.8 | 6.3 | 18.4 | 0.34 | 0.2960 |
| *yak*-into-*san* | exon | 0.2 | 0.0 | 0.3 | 0.03 | 0.2310 |
| *yak*-into-*san* | intergenic | 333.2 | 14.1 | 7.4 | 1.92 | 0.7320 |
| *yak*-into-*san* | intron | 1160.3 | 49.3 | 40.2 | 1.23 | 0.9910 |
| *san*-into-*yak* | 10kb inter | 611.8 | 18.2 | 19.0 | 0.96 | 0.8110 |
| *san*-into-*yak* | 2kb upstream inter | 166.8 | 5.0 | 8.5 | 0.59 | 0.7940 |
| *san*-into-*yak* | 3prime UTR | 33.1 | 1.0 | 3.5 | 0.28 | 0.0360 |
| *san*-into-*yak* | 5prime UTR | 18.7 | 0.6 | 2.8 | 0.20 | 0.0050 |
| *san*-into-*yak* | CDS | 252.5 | 7.5 | 18.4 | 0.41 | 0.0440 |
| *san*-into-*yak* | exon | 2.1 | 0.1 | 0.3 | 0.23 | 0.0120 |
| *san*-into-*yak* | intergenic | 403.4 | 12.0 | 7.4 | 1.63 | 0.9550 |
| *san*-into-*yak* | intron | 1869.2 | 55.7 | 40.2 | 1.39 | 0.1090 |
| *yak*-into-*tei* | 10kb inter | 18.0 | 31.0 | 19.0 | 1.63 | 0.5630 |
| *yak*-into-*tei* | 2kb upstream inter | 10.3 | 17.7 | 8.5 | 2.10 | 0.9510 |
| *yak*-into-*tei* | 3prime UTR | 0.3 | 0.5 | 3.5 | 0.15 | 0.1570 |
| *yak*-into-*tei* | 5prime UTR | 0.3 | 0.5 | 2.8 | 0.18 | 0.7430 |
| *yak*-into-*tei* | CDS | 9.7 | 16.6 | 18.4 | 0.90 | 1.0000 |
| *yak*-into-*tei* | exon | 0.0 | 0.0 | 0.3 | 0.00 | 0.0000 |
| *yak*-into-*tei* | intergenic | 4.1 | 7.0 | 7.4 | 0.94 | 0.0470 |
| *yak*-into-*tei* | intron | 15.6 | 26.7 | 40.2 | 0.66 | 0.0000 |
| *tei*-into-*yak* | 10kb inter | 14.5 | 22.5 | 19.0 | 1.18 | 0.8800 |
| *tei*-into-*yak* | 2kb upstream inter | 8.1 | 12.6 | 8.5 | 1.49 | 0.9940 |
| *tei*-into-*yak* | 3prime UTR | 0.6 | 0.9 | 3.5 | 0.25 | 0.0510 |
| *tei*-into-*yak* | 5prime UTR | 1.0 | 1.6 | 2.8 | 0.56 | 0.1960 |
| *tei*-into-*yak* | CDS | 18.5 | 28.6 | 18.4 | 1.55 | 0.8940 |
| *tei*-into-*yak* | exon | 0.0 | 0.0 | 0.3 | 0.00 | 0.0000 |
| *tei*-into-*yak* | intergenic | 2.3 | 3.6 | 7.4 | 0.49 | 0.6500 |
| *tei*-into-*yak* | intron | 19.5 | 30.3 | 40.2 | 0.75 | 0.0020 |

## REFERENCES

1. Rieseberg LH, Linder CR, Seiler GJ. Chromosomal and genic barriers to introgression in *Helianthus*. Genetics. 1995; 141:1163-71.

2. Wu C-I. The genic view of the process of speciation. J Evol Biol. 2001; 14: 851–865.

3. Nosil P. Speciation with gene flow could be common. Mol Ecol. 2008;17: 2103–2106.

4. Schumer M, Cui R, Rosenthal GG, Andolfatto P. Reproductive isolation of hybrid populations driven by genetic incompatibilities. PLoS Genet. Public Library of Science; 2015; 11: el005041.

5. Kulathinal RJ, Stevison LS, Noor MAF. The genomics of speciation in *Drosophila:* diversity, divergence, and introgression estimated using low-coverage genome sequencing. PLoS Genet. Public Library of Science; 2009; 5: el000550.

6. *Heliconius* Genome (THG) Consortium. Butterfly genome reveals promiscuous exchange of mimicry adaptations among species. Nature. 2012;487:94–98.

7. Martin SH, Dasmahapatra KK, Nadeau NJ, Salazar C, Walters JR, Simpson F, et al. Genome-wide evidence for speciation with gene flow in *Heliconius* butterflies. Genome Res. 2013;23:1817–1828.

8. Garrigan D, Kingan SB, Geneva AJ, Andolfatto P, Clark AG, Thornton KR, et al. Genome sequencing reveals complex speciation in the *Drosophila simulans* clade. Genome Res. 2012;22:1499–1511.

9. Schumer M, Cui R, Powell DL, Dresner R, Rosenthal GG, Andolfatto P. High-resolution mapping reveals hundreds of genetic incompatibilities in hybridizing fish species. Elife. 2014;3: e02535.

10. Fraïsse C, Belkhir K, Welch JJ, Bierne N. Local interspecies introgression is the main cause of extreme levels of intraspecific differentiation in mussels. Mol Ecol. 2016;25: 269–286.

11. Lindtke D, Gonzalez-Martinez SC, Macaya-Sanz D, Lexer C. Admixture mapping of quantitative traits in *Populus* hybrid zones: power and limitations. Heredity (Edinb). 2013;111: 474–485.

12. Harrison RG, Larson EL. Heterogeneous genome divergence, differential introgression, and the origin and structure of hybrid zones. Mol Ecol. 2016;25:2454–2466.

13. Abbott RJ, Barton NH, Good JM. Genomics of hybridization and its evolutionary consequences. Mol Ecol. 2016;25: 2325–2332.

14. Coyne JA, Orr HA. Speciation. Sinauer Associates Incorporated; 2004.

15. Dobzhansky T. Genetics and the Origin of Species (Classics of Modern Evolution Series). 1937. doi:10.1234/12345678

16. Mayr E. Animal species and evolution. Cambridge, MA and London, England: Harvard University Press; 1963.

17. Arnold ML. Natural Hybridization and Evolution. Oxford University Press. 1997.

18. Arnold ML. Evolution through genetic exchange. Oxford University Press. 2006.

19. Hedrick PW. Adaptive introgression in animals: examples and comparison to new mutation and standing variation as sources of adaptive variation. Mol Ecol. 2013;22:4606–4618.

20. Arnold ML, Martin NH. Adaptation by introgression. Journal of Biology 2009 6:4. BioMed Central; 2009;8: 82.

21. Fontaine MC, Pease JB, Steele A, Waterhouse RM, Neafsey DE, Sharakhov IV, et al. Extensive introgression in a malaria vector species complex revealed by phylogenomics. Science. 2015;347: 1258524-1258524.

22. Zhang W, Dasmahapatra KK, Mallet J, Moreira GRP, Kronforst MR. Genome-wide introgression among distantly related *HeJiconius butterfty* species. Genome Biol. BioMed Central; 2016;17: 25. doi:10.1186/s13059-016-0889-0

23. Baack EJ, Rieseberg LH. A genomic view of introgression and hybrid speciation. Curr Opin Genet Dev. 2007;17: 513–518.

24. Rosenzweig BK, Pease JB, Besansky NJ, Hahn MW. Powerful methods for detecting introgressed regions from population genomic data. Mol Ecol. 2016;25:2387–2397.

25. Price AL, Tandon A, Patterson N, Barnes KC, Rafaels N, Ruczinski I, et al. Sensitive detection of chromosomal segments of distinct ancestry in admixed populations. PLoS Genet. 2009;5: el000519.

26. Guan Y. Detecting Structure of Haplotypes and Local Ancestry. Genetics. Genetics; 2014; 196: 625–642.

27. Lawson DJ, Hellenthal G, Myers S, Falush D. Inference of population structure using dense haplotype data. PLoS Genet. Public Library of Science; 2012;8: el002453.

28. Sankararaman S, Mallick S, Dannemann M, Prüfer K, Kelso J, Pääbo S, et al. The genomic landscape of Neanderthal ancestry in present-day humans. Nature. 2014;507: 354-357.

29. Vernot B, Tucci S, Kelso J, Schraiber JG, Wolf AB, Gittelman RM, et al. Excavating Neandertal and Denisovan DNA from the genomes of Melanesian individuals. Science. 2016;352: 235-239.

30. Loh P-R, Lipson M, Patterson N, Moorjani P, Pickrell JK, Reich D, et al. Inferring admixture histories of human populations using linkage disequilibrium. Genetics. 2013;193:1233–1254.

31. Green RE, Krause J, Briggs AW, Maricic T, Stenzel U, Kircher M, et al. A Draft Sequence of the Neandertal Genome. Science. 2010;328: 710–722.

32. Pickrell JK, Pritchard JK. inference of population splits and mixtures from genome-wide allele frequency data. PLoS Genet. Public Library of Science; 2012;8: el002967.

33. Durand EY, Patterson N, Reich D, Slatkin M. Testing for ancient admixture between closely related populations. Mol Biol Evol. 2011; 28: 2239–2252

34. Matute DR, Ayroles JF. Hybridization occurs between *Drosophila simulaos* and *D. sechelliain* the Seychelles archipelago. J Evol Biol. 2014;27:1057–1068.

35. Pool JE, Corbett-Detig RB, Sugino RP, Stevens KA, Cárdeno CM, Crepeau MW, et al. Population genomics of sub-saharan *Drosophila melanogaster.* African diversity and non-African admixture. PLoS Genet. Public Library of Science; 2012;8: el003080.

36. Pool JE. The mosaic ancestry of the *Drosophila* genetic reference panel and the *D. melanogaster* reference genome reveals a network of epistatic fitness interactions. Mol Biol Evol. 2015; 32: 3236-3251.

37. Bachtrog D, Thornton K, Clark A, Andolfatto P. Extensive introgression of mitochondrial DNA relative to nuclear genes in the *Drosophila yakuba* species group. Evolution. 2006;60: 292-302.

38. Lachaise D, Harry M, Solignac M, Lemeunier F, Bénassi V, Cariou ML. Evolutionary novelties in islands: *Drosophila santomea,* a new *melanogaster* sister species from São Tomé. Proceedings of the Royal Society of London B: Biological Sciences. The Royal Society; 2000;267:1487-1495.

39. Llopart A, Lachaise D, Coyne JA. Multilocus analysis of introgression between two sympatric sister species of *Drosophila: Drosophilayakuba* and *D. santomea.* Genetics. Genetics; 2005;171:197–210.

40. Yassin A, Debat V, Bastide H, Gidaszewski N, David JR, Pool JE. Recurrent specialization on a toxic fruit in an island *Drosophila* population. Proc Natl Acad Sei USA. 2016;113: 4771–4776.

41. Comeault AA, Serrato Capuchina A, Turissini DA, McLaughlin PJ, David JR, Matute DR. A nonrandom subset of olfactory genes is associated with host preference in the fruit fly *Drosophila orena*. Evolution Letters. 2017;63:16.

42. Lachaise D, Cariou M-L, David JR, Lemeunier F, Tsacas L, Ashburner M. Historical biogeography of the *Drosophila melanogaster* species subgroup. Evolutionary Biology. Boston, MA: Springer US; 1988. pp. 159–225.

43. Lachaise D, Lemeunier F. Clinal variations in male genitalia in Drosophila teissieri Tsacas. The American Naturalist. 1981 Apr 1;117(4):600–608.

44. Comeault AA, Venkat A, Matute DR. Correlated evolution of male and female reproductive traits drive a cascading effect of reinforcement in *Drosophila yakuba*. Proc Biol Sci. 2016;283: 20160730.

45. Cariou ML, Silvain JF, Daubin V, Da Lage JL, Lachaise D. Divergence between *Drosophila santomea and* allopatric or sympatric populations of *D. yakuba* using paralogous amylase genes and migration scenarios along the Cameroon volcanic line. Mol Ecol. 2001; 10: 649–660.

46. Coyne JA, Kim SY, Chang AS, Lachaise D, Elwyn S. Sexual isolation between two sibling species with overlapping ranges: *Drosophila santomea* and *Drosophila yakuba*. Evolution. 2002;56: 2424–2434.

47. Coyne JA, Elwyn S, Kim SY, Llopart A. Genetic studies of two sister species in the *Drosophila melanogaster* subgroup, *D. yakuba* and *D. santomea*. Genetical research. 2004; 84:11–26.

48. Moehring AJ, Llopart A, Elwyn S, Coyne JA, Mackay TFC. The genetic basis of postzygotic reproductive isolation between *Drosophila santomea* and *D. yakuba* due to hybrid male sterility. Genetics. Genetics; 2006;173: 225–233.

49. Llopart A, Lachaise D, Coyne JA. An anomalous hybrid zone in *Drosophila*. Evolution. 2005; 59:2602–2607.

50. Cooper BS, Ginsberg PS, Turelli M, Matute DR. *Wolbachia* in the *Drosophila yakuba* complex: pervasive frequency variation and weak cytoplasmic incompatibility, but no apparent effect on reproductive isolation. Genetics. Genetics; 2017;205: 333–351.

51. Llopart A, Elwyn S, Lachaise D, Coyne JA. Genetics of a difference in pigmentation between *Drosophila yakuba* and *Drosophila santomea*. Evolution. 2002;56(11):2262-2277.

52. Carbone MA, Llopart A, deAngelis M, Coyne JA, Mackay TFC. Quantitative trait loci affecting the difference in pigmentation between *Drosophila yakubaand D. santomea*. Genetics; 2005;171: 211–225.

53. Cobb M, Huet M, Lachaise D, Veuille M. Fragmented forests, evolving flies: molecular variation in African populations of *Drosophila teissieri*. Mol Ecol. 2000;9:1591–1597.

54. Cooper BS, Sedghifar A, Nash WT, Comeault AA, Matute DR. A maladaptive combination of traits contributes to the maintenance of a stable hybrid zone between two divergent species of *Drosophila.* bioRxiv. Cold Spring Harbor Labs Journals; 2017;: 138388. doi:10.1101/138388

55. Lemeunier F, Ashburner M. Relationships within the *melanogaster* Species subgroup of the genus *Drosophila (Sophophorà).* II. Phylogenetic relationships between six species based upon polytene chromosome banding sequences. Proceedings of the Royal Society of London B: Biological Sciences; 1976; 193: 275–294.

56. Monnerot M, Solignac M, Wolstenholme DR. Discrepancy in divergence of the mitochondrial and nuclear genomes of *Drosophila teissieri and Drosophila yakuba*. Journal of Molecular Evolution. 1990; 30:500-508.

57. Turissini DA, Liu G, David JR, Matute DR. The evolution of reproductive isolation in the *Drosophila yakuba* complex of species. J Evol Biol. 2015;28: 557-575.

58. Llopart A, Herrig D, Brud E, Stecklein Z. Sequential adaptive introgression of the mitochondrial genome in *Drosophila yakuba* and *Drosophila santomea*. Mol Ecol. 2014;23:1124–1136.

59. Beck EA, Thompson AC, Sharbrough J, Brud E, Llopart A. Gene flow between *Drosophila yakuba* and *Drosophila santomea* in subunit V of cytochrome c oxidase: A potential case of cytonuclear cointrogression. Evolution. 2015;69: 1973–1986.

60. Langley CH, Stevens K, Cardeno C, Lee YCG, Schrider DR, Pool JE, et al. Genomic variation in natural populations of *Drosophila melanogaster*. Genetics; 2012;192: 533–598

61. Garud NR, Petrov DA. Elevation of linkage disequilibrium above neutral expectations in ancestral and derived populations of *Drosophila melanogaster*. Genetics; 2016;203: 863-880

62. Garud NR, Messer PW, Buzbas EO, Petrov DA. Recent selective sweeps in North American *Drosophila melanogaster* show signatures of soft sweeps. PLoS Genet. Public Library of Science; 2015;11: el005004.

63. Llopart A, Elwyn S, Lachaise D, Coyne JA. Genetics of a difference in pigmentation between *Drosophil ayakuba* and Drosophila santomea. 2002: 56:2262-2277.

64. Begun DJ, Holloway AK, Stevens K, Hillier LW, Poh Y-P, Hahn MW, et al. Population Genomics: Whole-genome analysis of polymorphism and divergence in *Drosophila simulans*. PLoS Biol. Public Library of Science; 2007;5:e310.

65. Hey J, Kliman RM. Population genetics and phylogenetics of DNA sequence variation at multiple loci within the *Drosophila melanogaster* species complex. Mol Biol Evol. 1993; 10:804-822

66. Tamura K, Subramanian S, Kumar S. Temporal patterns of fruit fly *(.Drosophila*) evolution revealed by mutation clocks. Mol Biol Evol. 2004;21: 36–44.

67. Russo CA, Takezaki N, Nei M. Molecular phylogeny and divergence times of drosophilid species. Mol Biol Evol. 1995; 12:391-404.

68. Sturtevant AH. Genetic Studies on *Drosophila simulanss.* I. Introduction. Hybrids with *Drosophila melanogaster*. Genetics; 1920;5: 488–500.

69. Muirhead CA, Presgraves DC. Hybrid Incompatibilities, local adaptation, and the genomic distribution of natural introgression between species. The American Naturalist. 2015;187: 249–261.

70. Charlesworth B, Coyne JA, Barton NH. The relative rates of evolution of sex chromosomes and autosomes. The American Naturalist; 2015;130: 113–146.

71. Martin SH, Davey JW, Jiggins CD. Evaluating the use of ABBA-BABA statistics to locate introgressed loci. Mol Biol Evol. 2015; 32: 244-257

72. Turissini DA, McGirr JA, Patel SS, David JR, Matute DR. The rate of evolution of postmating-prezygotic reproductive isolation in *Drosophila.* bioRxiv. Cold Spring Harbor Labs Journals; 2017;: 142059. doi:10.1101/142059

73. Purcell S, Neale B, Todd-Brown K, Thomas L, Ferreira MAR, Bender D, et al. PLINK: a tool set for whole-genome association and population-based linkage analyses. American Journal of Human Genetics. 2007;81: 559-575.

74. Corbett-Detig R, Jones M. SELAM: simulation of epistasis and local adaptation during admixture with mate choice. Bioinformatics. 2016; 32:3035-3037.

75. Pollard DA, Iyer VN, Moses AM, Eisen MB. Widespread discordance of gene trees with species tree in *Drosophila:* Evidence for incomplete lineage sorting. PLoS Genet. Public Library of Science; 2006;2: e173.

76. Joly S, McLenachan PA, Lockhart PJ. A Statistical approach for distinguishing hybridization and incomplete lineage sorting. The American Naturalist 2015;174: E54-E70.

77. Matute DR. Reinforcement of gametic isolation in *Drosophila*. PLoS Biol. Public Library of Science; 2010;8: el000341.

78. Matute DR, Coyne JA. Intrinsic reproductive isolation between two sister species of *Drosophila*. Evolution. 2010;64: 903–920.

79. Matute DR, Novak CJ, Coyne JA. Temperature-based extrinsic reproductive isolation in two species of *Drosophila*. Evolution. 2009;63: 595–612.

80. Matute DR. The magnitude of behavioral isolation is affected by characteristics of the mating community. Ecology and Evolution. 2014;4: 2945–2956.

81. Matute DR, Coyne JA. Intrinsic reproductive isolation between two sister species of *Drosophila*. Evolution. 2010;64: 903–920.

82. Orr HA. Haldane’s rule. Annual Review of Ecology and Systematics. 1997; 28(1):195–218.

83. Wu CI, Johnson NA, Palopoli MF. Haldane’s rule and its legacy: Why are there so many sterile males? Trends in Ecology & Evolution. 1996; 11: 281–284.

84. Delph LF, Demuth JP. Haldane’s rule: Genetic bases and their empirical support. Journal of Heredity. 2016.

85. Coyne JA. The genetic basis of Haldane’s rule. Nature. 1985; 314: 736–738.

86. Laurie CC. The weaker sex is heterogametic: 75 years of Haldane’s rule. Genetics. 1997.147: 937–951.

87. Sankararaman S, Mallick S, Patterson N, Reich D. The combined landscape of Denisovan and Neanderthal ancestry in present-day humans. Curr Biol. 2016;26:1241–1247.

88. Payseur BA, Krenz JG, Nachman MW. Differential patterns of introgression across the *X* chromosome in a hybrid zone between two species of house mice. Evolution. 2004;58: 2064–2078.

89. Carneiro M, Blanco Aguiar JA, Villafuerte R, Ferrand N, Nachman MW. Speciation in the European rabbit (*Oryctolagus cuniculus*): islands of differentiation on the *X* chromosome and autosomes. Evolution. B 2010;64: 3443-3460.

90. Carneiro M, Albert FW, Afonso S, Pereira RJ, Burbano H, Campos R, et al. The genomic architecture of population divergence between subspecies of the European rabbit. PLoS Genet. Public Library of Science; 2014;10: el003519.

91. Coyne JA, Orr HA. Two rules of speciation. In Otte O. & Endler JA. (eds.), Speciation and its Consequences. 1989 Sinauer Associates.

92. Coyne JA. Genetics and speciation. Nature. 1992;355: 511-515.

93. Masly JP, Presgraves DC. High-resolution genome-wide dissection of the two rules of speciation in *Drosophila.* PLoS Biol. Public Library of Science; 2007;5:e243.

94. Llopart A. The rapid evolution of X-linked male-biased gene expression and the large-Aeffect in *Drosophilayakuba, D. santomea,* and their hybrids. Mol Biol Evol. 2012.29:3873-86.

95. Pool JE, Nielsen R. Inference of historical changes in migration rate from the lengths of migrant tracts. Genetics; 2009;181: 711–719.

96. Gravel S. Population genetics models of local ancestry. Genetics; 2012;191: 607-619.

97. Liang M, Nielsen R. The Lengths of Admixture Tracts. Genetics; 2014; 197: 953–967.

98. Turissini DA, Comeault AA, Liu G, Lee YCG, Matute DR. The ability of *Drosophila* hybrids to locate food declines with parental divergence. Evolution. 2017;71: 960-973.

99. Matute DR. Noisy neighbors can hamper the evolution of reproductive isolation by reinforcing selection. The American Naturalist; 2015; 185: 253269.

100. Matute DR. Reinforcement can overcome gene flow during speciation in *Drosophila*. Curr Biol. 2010;20: 2229–2233.

101. Fraïsse C, Roux C, Welch JJ, Bierne N. Gene-Flow in a mosaic hybrid zone: is local introgression adaptive? Genetics; 2014;197: 939–951.

102. Yuan Q, Song Y, Yang C-H, Jan LY, Jan YN. Female contact modulates male aggression via a sexually dimorphic GABAergic circuit in *Drosophila*. Nature Neuroscience; 2014;17: 81–88.

103. Mast JD, De Moraes CM, Alborn HT, Lavis LD, Stern DL. Evolved differences in larval social behavior mediated by novel pheromones. Elife. 2014;3: e04205.

104. Thistle R, Cameron P, Ghorayshi A, Dennison L, Scott K. Contact Chemoreceptors mediate male-male repulsion and male-female attraction during *Drosophila* courtship. Cell. 2012;149:1140–1151.

105. Adewoye AB, Kyriacou CP, Tauber E. Identification and functional analysis of early gene expression induced by circadian light-resetting in *Drosophila*. BMC Genomics 2011 12:1. BioMed Central; 2015;16: 570.

106. Sone M, Suzuki E, Hoshino M, Hou D. Synaptic development is controlled in the periactive zones of *Drosophila synapses*. Development. 2000.127:41574168.

107. Brown EB, Layne JE, Zhu C, Jegga AG, Rollmann SM. Genome-wide association mapping of natural variation in odour-guided behaviour in *Drosophila.* Genes, Brain and Behavior. 2013;12: 503–515.

108. Lu H-L, Wang JB, Brown MA, Euerle C, Leger RJS. Identification of *Drosophila* mutants affecting defense to an entomopathogenic fungus. Scientific Reports. 2015;5: 697.

109. Morozova TV, Huang W, Pray VA, Whitham T, Anholt RRH, Mackay TFC. Polymorphisms in early neurodevelopmental genes affect natural variation in alcohol sensitivity in adult Drosophila. BMC Genomics 2011 12:1. 4 ed. BioMed Central; 2015; 16: 865.

110. Ni JD, Baik LS, Holmes TC, Montell C. A rhodopsin in the brain functions in circadian photoentrainment in *Drosophila*. Nature. 2017; 545:340-344.

111. Rumball W, Franklin IR, Frankham R, Sheldon BL. Decline in heterozygosity under full-sib and double first-cousin inbreeding in *Drosophila* melanogaster. Genetics; 1994;136:1039–1049.

112. David JR, Gibert P, Legout H, Pétavy G, Capy P, Moreteau B. Isofemale lines in *Drosophila:* an empirical approach to quantitative trait analysis in natural populations. Heredity (Edinb). 2005;94: 3-12.

113. Barton N, Bengtsson BO. The barrier to genetic exchange between hybridising populations. Heredity (Edinb). 1986;57: 357-376.

114. Uecker H, Setter D, Hermisson J. Adaptive gene introgression after secondary contact. J Math Biol. 2015;70:1523-1580.

115. Juric I, Aeschbacher S, Coop G. The strength of selection against Neanderthal introgression. PLoS Genet. Public Library of Science; 2016; 12: el006340.

116. Barton NH. Adaptation, speciation and hybrid zones. Nature. 1989;341: 497-503.

117. Barton NH, Hewitt GM. Analysis of hybrid zones. Annual review of Ecology and Systematics. 1985;16:113-48.

118. Barton NH. Does hybridization influence speciation? J Evol Biol. 2013;26: 267-269.

119. Moore WS. An evaluation of narrow hybrid zones in vertebrates. The Quarterly Review of Biology. 2015;52: 263-277. doi:10.1086/409995

120. Harris K, Nielsen R. Inferring demographic history from a spectrum of shared haplotype lengths. PLoS Genet. Public Library of Science; 2013;9: el003521.

121. Wall JD, Hammer MF. Archaic admixture in the human genome. Curr Opin Genet Dev. 2006; 16: 606–610.

122. Hellenthal G, Busby GBJ, Band G, Wilson JF, Capelli C, Falush D, et al. A genetic atlas of human admixture history. Science; 2014;343: 747–751.

123. Sugden LA, Ramachandran S. Integrating the signatures of demie expansion and archaic introgression in studies of human population genomics. Curr Opin Genet Dev. 2016;41: 140–149.

124. Deschamps M, Laval G, Fagny M, Itan Y, Abel L, Casanova J-L, et al. Genomic Signatures of selective pressures and introgression from archaic hominins at human innate immunity genes. The American Journal of Human Genetics. 2016;98:5–21.

125. McCoy RC, Wakefield J, Akey JM. Impacts of Neanderthal-introgressed sequences on the landscape of human gene expression. Cell. 2017;168: 916–927.el2.

126. Racimo F, Sankararaman S, Nielsen R, Huerta-Sánchez E. Evidence for archaic adaptive introgression in humans. Nat Rev Genet; 2015;16: 359371.

127. Eaton D, Ree RH. Inferring phylogeny and introgression using RADseq data: an example from flowering plants (Pedicularis: *Orobanchaceae*). Systematic biology. 2013; 62:689–706.

128. Senerchia N, Felber F, North B, Sarr A, Guadagnuolo R, Parisod C. Differential introgression and reorganization of retrotransposons in hybrid zones between wild wheats. Mol Ecol. 2016;25: 2518–2528.

129. Gross BL, Rieseberg LH. The ecological genetics of homoploid hybrid speciation. J Hered. 2005;96: 241–252.

130. Soltis PS, Soltis DE. The role of hybridization in plant speciation. Annual review of plant biology. 2009; 60: 561–588.

131. Winger BM. Consequences of divergence and introgression for speciation in Andean cloud forest birds. Evolution. 2017. doi:10.1111/evo.13251

132. Rheindt FE, Fujita MK, Wilton PR. Introgression and phenotypic assimilation in *Zimmerius* flycatchers (Tyrannidae): population genetic and phylogenetic inferences from genome-wide SNPs. Systematic Biology. 2014 63:134–152.

133. Moyle RG, Manthey JD, Hosner PA, Rahman M, Lakim M, Sheldon FH. A genome-wide assessment of stages of elevational parapatry in Bornean passerine birds reveals no introgression: implications for processes and patterns of speciation. PeerJ; 2017;5: e3335.

134. Toews DPL, Campagna L, Taylor SA, Balakrishnan CN, Baldassarre DT, Deane-Coe PE, et al. Genomic approaches to understanding population divergence and speciation in birds. The Auk; 2015;133:13–30.

135. Rheindt FE, Edwards SV. Genetic introgression: an integral but neglected component of speciation in birds. The Auk; 2011;128: 620–632.

136. Abbott R, Albach D, Ansell S, Arntzen JW, Baird SJE, Bierne N, et al. Hybridization and speciation. J Evol Biol. 2013;26: 229–246.

137. Mallet J. Hybridization as an invasion of the genome. Trends in Ecology & Evolution. 2005;20: 229–237.

138. Schwenk K, Brede N, Streit B. Introduction. Extent, processes and evolutionary impact of interspecific hybridization in animals. Philosophical Transactions of the Royal Society of London B: Biological Sciences. The Royal Society; 2008;363: 2805–2811.

139. Baack EJ, Rieseberg LH. A genomic view of introgression and hybrid speciation. Curr Opin Genet Dev. 2007;17: 513–518.

140. Phaff HJ. A new method of collecting *Drosophila* by means of sterile bait. The American Naturalist. 1955; 89:53–54.

141. Matute DR. noisy neighbors can hamper the evolution of reproductive isolation by reinforcing selection. The American Naturalist. 2015; 185: 253269.

142. Rogers RL, Cridland JM, Shao L, Hu TT. Landscape of standing variation for tandem duplications in *Drosophilayakuba* and *Drosophila simulans*. Molecular biology and evolution. 2014;31:1750–1766

143. Clark AG, Eisen MB, Smith DR, Bergman CM, Oliver B, Markow TA, étal. Evolution of genes and genomes on the *Drosophila* phylogeny. Nature. 2007;450:203–218.

144. Li H, Durbin R. Fast and accurate long-read alignment with BurrowsWheeler transform. Bioinformatics. Oxford University Press; 2010;26: 589595.

145. Li H, Handsaker B, Wysoker A, Fennell T, Ruan J, Homer N, et al. The Sequence Alignment/Map format and SAMtools. Bioinformatics. 2009;25: 2078–2079.

146. McKenna A, Hanna M, Banks E, Sivachenko A, Cibulskis K, Kernytsky A, et al. The Genome Analysis Toolkit: a MapReduce framework for analyzing next-generation DNA sequencing data. Genome Res. 2010;20: 1297–1303.

147. DePristo MA, Banks E, Poplin R, Garimella KV, Maguire JR, Hartl C, et al. A framework for variation discovery and genotyping using next-generation DNA sequencing data. Nat Genet. 2011;43: 491–498.

148. Jombart T. adegenet: a R package for the multivariate analysis of genetic markers. Bioinformatics. 2008;24:1403–1405.

149. Prüfer K, Racimo F, Patterson N, Jay F, Sankararaman S, Sawyer S, et al. The complete genome sequence of a Neanderthal from the Altai Mountains. Nature; 2014;505:43–49.

150. Busing FMTA, Meijer E, Van Der Leeden R. Delete-m Jackknife for Unequal m. Statistics and Computing; 1999;9: 3–8. 8

151. Presgraves DC. Sex chromosomes and speciation in *Drosophila*. Trends in Genetics; 2008;24: 336–343.

152. Kimura M, Ohta T. The Average Number of Generations until Fixation of a Mutant Gene in a Finite Population. Genetics; 1969;61: 763–771.

153. Gao Z, Przeworski M, Sella G. Footprints of ancient-balanced polymorphisms in genetic variation data from closely related species. Evolution. 2015;69: 431–446.

154. Comeron JM, Ratnappan R, Bailin S. The many landscapes of recombination in *Drosophila melanogaster*. PLoS Genet. Public Library of Science; 2012;8: e1002905.

